# Accounting for social feasibility in landscape-scale scenarios for restoring nature beyond protected areas

**DOI:** 10.64898/2026.09.11.750937

**Authors:** Alberto González-García, Margot Neyret, Adrián López-Tejedor, Marie Caroline Prima, Sara Si-Moussi, Sandra Lavorel

## Abstract

1. Global restoration targets, including the Kunming-Montreal Global Biodiversity Framework and the European Union’s Nature Restoration Regulation, call for extending restoration beyond protected area boundaries to reconnect isolated ecosystems, yet implementing these landscape-scale interventions forces trade-offs between biophysical effectiveness and social feasibility.

2. In the Grenoble region of the French Alps, we used a gradient analysis of 12 ecosystem services across protected area interfaces to design and compare two restoration scenarios. Both were built from the same set of co-selected interventions but differed only in their spatial allocation: a ‘Technical’ scenario targeting areas of low ecosystem service continuity around protected areas, and a ‘Participatory’ scenario co-designed with local stakeholders.

3. Both scenarios physically modified about 14.5% of the landscape, mostly through a forest-based adaptation strategy, yet their effects reached up to more than 60% of the territory for the most connectivity-dependent services, showing that benefits propagate well beyond the area directly modified.

4. The two scenarios diverged mainly in how they treated agricultural land. The Technical scenario reduced abrupt drops in ecosystem service provision at the protected area edge nearly twice as often as the Participatory one (46% versus 26% of the sharp declines it addressed), and at six locations removed them entirely. However, it concentrated over 99% of its non-forest effort in agricultural diversification, such as diversifying crops and adding hedgerows, the action stakeholders ranked hardest to implement, whereas the Participatory scenario favoured more feasible actions such as river restoration, which met less resistance but missed several critical discontinuities in ecosystem service supply.

5. These contrasts show that biophysical potential and social feasibility can pull restoration in different directions, and that the more effective spatial configuration is not necessarily the more implementable one.

6. Closing the implementation gap for nature-based solutions depended less on expanding the area restored than on where interventions were placed. Gradient-based diagnostics appear most useful not as fixed prescriptions but as tools to steer existing sectoral funding in agriculture, urban planning and forestry toward high-impact locations, an approach transferable to other regions working to reconnect protected areas with their surrounding landscapes as coupled social-ecological systems.

## 1. Introduction

The pervasive decline of biodiversity places planetary ecosystems at increasing risk (IPBES, 2019; Obura, 2023). This necessitates a fundamental shift in conservation strategies, moving from isolated protection toward landscape-scale restoration (Díaz et al., 2019). Although protected areas remain a cornerstone of conservation, they often function as disconnected elements in human-dominated landscapes (Watson et al., 2014). This isolation limits their effectiveness and the flow of ecosystem services for human well-being (Kremen and Merenlender, 2018). In response, major global and regional policy frameworks, such as the Kunming–Montreal Global Biodiversity Framework and the European Union’s Nature Restoration Regulation, call for integrated action. This framework explicitly highlights the need to enhance the connectivity of protected areas and substantially restore degraded ecosystems (Targets 2 and 3). Similarly, the EU regulation sets legally binding restoration targets, requiring coordinated interventions across working landscapes, including agricultural, forest and urban areas (Perissi, 2025). Achieving these objectives demands extending conservation efforts beyond protected areas boundaries and strategically intervening in the surrounding matrix (Chapman et al., 2025). This shift is needed to secure ecological connectivity and sustain the provision of ecosystem services from protected areas to adjacent agricultural, peri-urban and urban communities (Mitchell et al., 2015).

Nature-based solutions (NbS) promise to achieve large-scale restoration across working and urban landscapes (Seddon et al., 2021), acting as strategic interventions to bridge the gap between isolated protected areas and their surrounding matrix. NbS have gained traction as a pathway for transformative change (Palomo et al., 2021; Seddon et al., 2020; IPBES, 2024). Defined by their capacity to address societal challenges while benefiting human well-being and biodiversity (Cohen-Shacham et al., 2019), NbS are inherently multifunctional and can be cost-effective (González-García et al., 2025). For instance, implementing hedgerows enhances connectivity for biological control agents while improving carbon storage, generating multiple ecosystem services (Keesstra et al., 2018). Similarly, urban greening mitigates heatwaves, extending regulating ecosystem services from protected areas into populated zones (Raymond et al., 2023). However, NbS success relies on strategic landscape-scale deployment to maximize co-benefits and navigate trade-offs from competing land uses (Cohen-Shacham et al., 2019). This presents a dual challenge: quantifying the biophysical effectiveness of NbS in enhancing ecosystem services, while ensuring they align with diverse social values.

This challenge can only be addressed by a comprehensive assessment framework which integrates the spatial and social factors that modulate ecosystem service provision across complex interfaces such as protected area boundaries (Metzger et al., 2021). First, it must map ecosystem service supply and analyse bundles of multiple ecosystem services for diagnosing latent trade-offs and synergies in diverse landscapes (Raudsepp-Hearne et al., 2010; Saidi and Spray, 2018). Secondly, because stakeholder preferences influence prioritization, grouping ecosystem services by the priorities of different stakeholder groups, rather than by their biophysical category alone, supports management applicability (Martín-López et al., 2014; Plieninger et al., 2019; González-García et al., 2026). Framing bundles this way reflects how distinct actors perceive and prioritise the benefits a landscape provides, making trade-offs legible to the groups who manage them. Thirdly, assessing NbS feasibility is a prerequisite for implementation at scale. Implementation success hinges on enabling factors and the absence of barriers such as conflicting institutional objectives or polarized social acceptance (Cohen-Shacham et al., 2019; Bruley et al., 2025). Identifying these factors in conflict-prone areas, particularly at agricultural and urban interfaces adjacent to protected areas, is crucial for unlocking the transformative potential of NbS to reconnect the landscape.

Successful NbS implementation requires acknowledging intertwined people–nature relations (Locatelli et al., 2025). Interventions must be both “nature-based” and “people-based” (Welden et al., 2021), integrating intrinsic, instrumental, and relational values (Pascual et al., 2017; Chan et al., 2016). While technical analysis remains vital for diagnosing spatial limitations in ecosystem services (Metzger et al., 2021), a purely technical approach risks overlooking the heterogeneous preferences and power dynamics across landscapes (Nesshöver et al., 2017). These dynamics are often most pronounced at the very edges of protected areas, where conservation mandates collide with economic land uses, highlighting a critical research gap: determining whether the theoretical ecological benefits of technical optimization can overcome the implementation barriers arising from social conflicts (Bruley et al., 2025). Consequently, directly contrasting the biophysical effectiveness of a strategy prioritizing technical maximization with an alternative designed to maximize feasibility and reduce conflict at protected area interfaces represents a research priority for the successful scaling of NbS.

Building on the need to reconcile technical rigor with social legitimacy in NbS (Cousins, 2021; Nelson et al., 2020), this study addresses three central questions: (i) which implementation strategy more effectively enhances ecosystem services, contrasting a technical diagnosis based on expert assessments with a participatory approach grounded in local priorities; (ii) whether these strategies modify the functional structure of the landscape in different ways, including examining the role of protected areas within the wider matrix through ecosystem service gradients; and (iii) if the most ecologically effective strategy aligns with its feasibility for local actors. To answer these questions, we applied a coupled participatory and spatial modeling approach in the Grenoble region, a high-contrast interface between protected areas and human-dominated working landscapes. We contrasted two restoration scenarios: a Participatory scenario based on priorities and implementation barriers identified through stakeholder workshops, and a Technical scenario derived from an expert diagnosis of discontinuous ecosystem service gradients. To model biophysical dynamics, we quantified spatial patterns in 12 ecosystem services, incorporating human demand and grouping them into three bundles reflecting the management priorities of different stakeholder groups (Rural, Cultural, and Urban). Both scenarios were implemented using explicit spatial rules for land-use change validated by stakeholders. This integrated assessment allows us to directly examine the trade-off between technical optimization and social feasibility in landscape restoration, providing empirical insights into the ambition of NbS to enhance ecosystem service supply at landscape scale.

## 2. Methods

### 2.1 Study area and overall workflow

Our study was conducted in the Grenoble region of the French Alps, a well-studied landscape characterized by pronounced environmental and land-use gradients. The region encompasses a heterogeneous matrix stretching from densely populated, agriculturally intensive valleys to high-elevation alpine ecosystems (Vannier et al., 2019). From the diverse network of protected areas in this region, we selected 16 sites, excluding very small reserves to ensure a robust spatial analysis. This context provides an ideal setting for studying the interfaces between protected areas and human-dominated landscapes (see Supplementary Information A for a detailed description and map of the study area).

The research followed an integrated workflow consisting of a foundational diagnostic and two subsequent novel phases (Fig. 1). We relied on a recently developed spatial framework to characterize the current state of ecosystem service provision across protected area borders (González-García et al., 2026). In this prior analysis, 12 ecosystem services were modeled (Fig. 1A), ranging from carbon storage, food production, heatwave mitigation, flood regulation, erosion control, and outdoor recreation, to connectivity-dependent ecosystem services mediated by mobile species: pollination, biological control, seed dispersal, emblematic species, mosquito control, and hunting value. Following previous stakeholder-based assessments in the region (Vannier et al., 2019), these ecosystem services had been aggregated into three bundles reflecting the management priorities of different stakeholder groups (Rural, Cultural, and Urban) to analyze potential trade-offs. This previous work also utilized an automated process to generate perpendicular transects along protected area border segments (Fig. 1B), quantifying how the provision of each bundle changes from the interior to the exterior of the protected areas (Fig. 1C).

**Figure 1.**
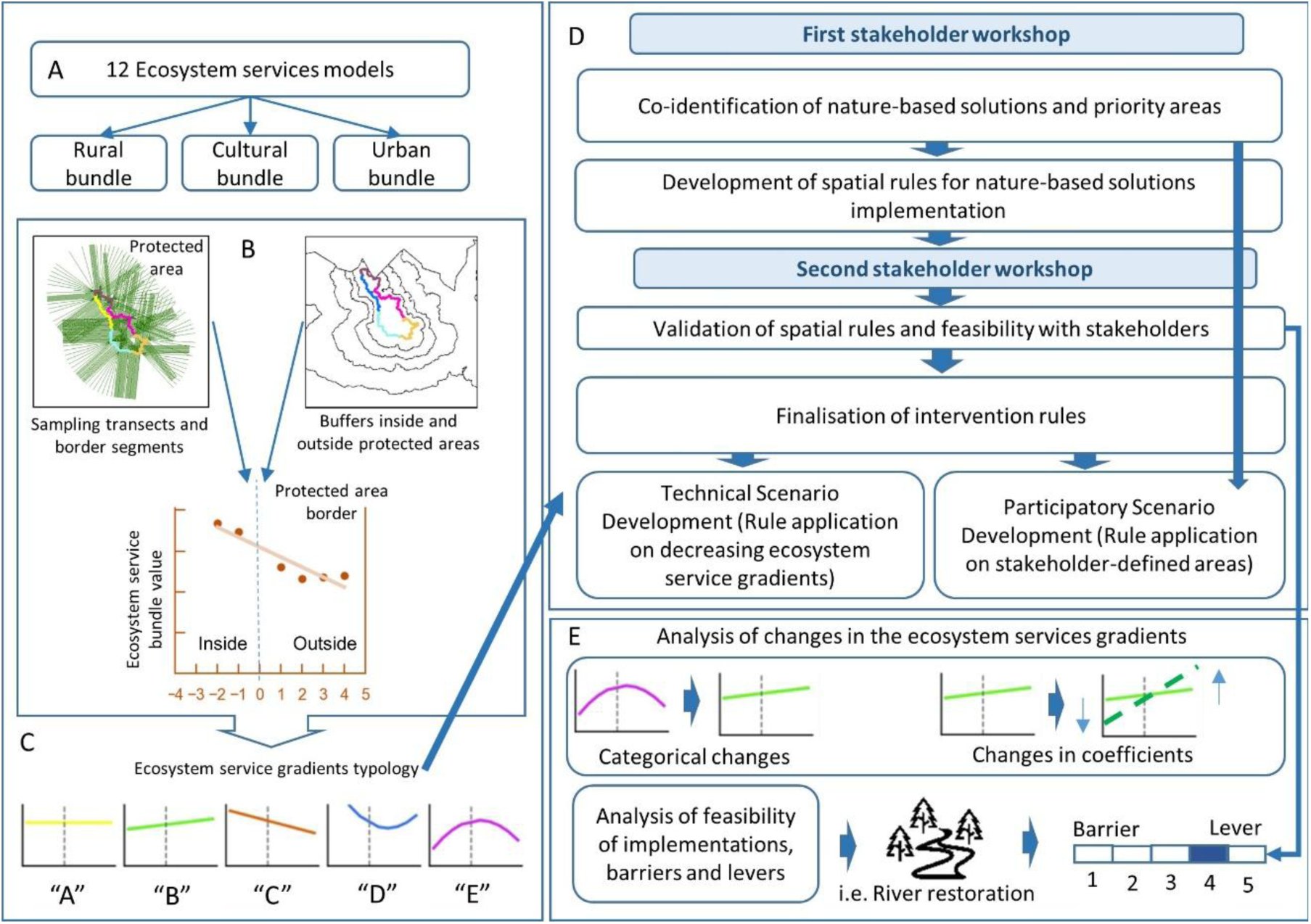
Conceptual workflow of the study. The research integrates a baseline assessment, scenario development, and comparative analysis. (A) We modeled 12 ecosystem services and aggregated them into three bundles. (B) We analyzed the gradients of these bundles across protected area borders using transects and structural buffers to generate a gradient profile for each border segment. (C) These profiles were classified into a five-category typology representing distinct patterns of ecosystem service provision across the border. (D) We developed two future scenarios through a process involving stakeholder workshops: a ‘Participatory’ scenario where interventions were placed in stakeholder-defined priority areas, and a ‘Technical’ scenario where interventions targeted border segments exhibiting sharp declines or discontinuities in the baseline gradient analysis. (E) Finally, we implemented a comparative analysis to assess the impacts of both scenarios, examining categorical changes in border patterns, modulations in the shape of gradients, and the trade-off with implementation feasibility as perceived by stakeholders.

Leveraging this established context, the core of the present study focuses on two new methodological steps. First, we developed two future scenarios for implementing NbS (Fig. 1D). To do so, we first conducted two workshops with local stakeholders to co-develop a set of spatial rules for actions such as implementing hedgerows in agricultural landscapes. Then, for the ‘Participatory’ scenario, stakeholders defined the specific locations for these interventions. For the ‘Technical’ scenario, the same set of rules was applied, but the locations were guided by a gradient analaysis targeting border segments with declining or discontinuous gradients. Third, we implemented a comparative analysis framework to quantify the impacts of these scenarios on ecosystem services provision and border gradient patterns (Fig. 1E), and to evaluate the trade-offs between their biophysical effectiveness and perceived implementation feasibility.

### 2.2 Baseline assessment: characterizing ecosystem service gradients

As the baseline ecosystem service gradients were fully developed in a preceding study (González-García et al., 2026), we briefly summarize that methodology here to contextualize our scenario simulations. We selected the 12 key ecosystem services based on their high relevance for the socio-ecological context of the French Alps, as identified in prior regional stakeholder assessments (Vannier et al., 2019). These ecosystem services were quantified using two main modeling approaches (Table 1; see Supplementary Information B for detailed model descriptions). Six services were modeled using established spatially-explicit tools such as InVEST and empirical models (Natural Capital Project, 2023; Lasseur et al., 2018; Byczek et al., 2018). The remaining six ecosystem services are provided by mobile organisms and were modeled with connectivity-based approaches that explicitly incorporate species movement. Pollination was represented through an insect-pollinator archetype (Schulp et al., 2014), whereas biological control, mosquito control, seed dispersal, hunting value and emblematic species were modeled from the movement of the relevant vertebrate groups, for example insectivores and predators for biological and mosquito control, using a multi-species connectivity framework (Prima et al., 2024) implemented with Omniscape (McRae et al., 2016). For all ecosystem services where relevant, we integrated spatially explicit human demand criteria to ensure management applicability (e.g., restricting the analysis of pollination to crop areas). All individual ecosystem services maps were normalized to a common 0-1 scale (see Supplementary Information C for normalization methods) and then aggregated into three bundles reflecting distinct societal value domains: Rural, Cultural, and Urban (Table 1).

**Table 1.**
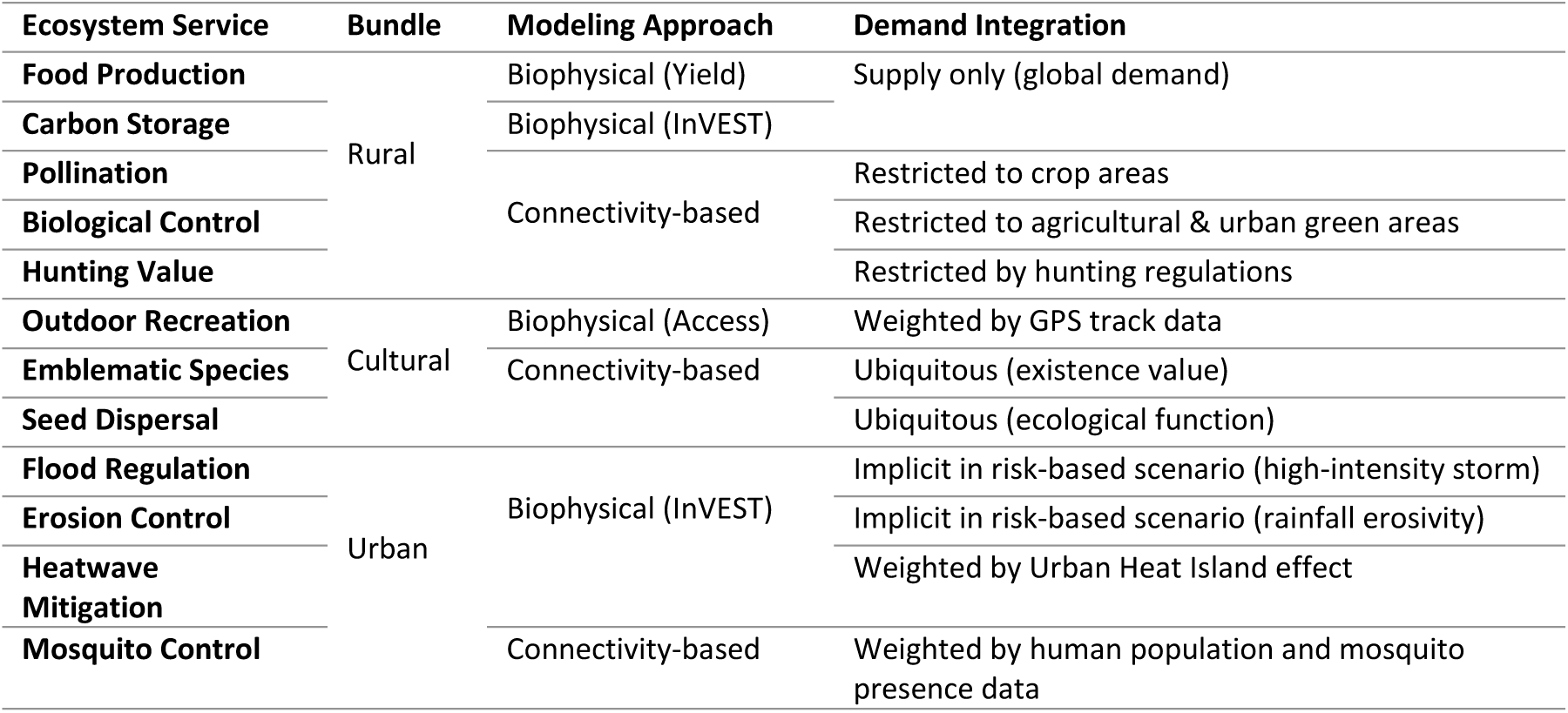
Overview of the 12 ecosystem services modeled in the study. The table details each service its assignment to one of the three bundles, reflecting distinct stakeholder management priorities, the general modeling approach used for its quantification, and the method for integrating human demand criteria into the final service map. Detailed model parameters and methodologies are provided in Supplementary Information B.

To characterise patterns of ES variation across protected area boundaries, we first divided each protected area border into segments as primary units of analysis. This process classified each border segment into one of five pattern types. Type A (’Flat Gradient’) indicates a seamless transition, i.e. similar ecosystem service supply in and outside the protected area. Type B (’Increasing Gradient’) shows that service provision is higher outside the protected area. Type C (’Decreasing Gradient’) represents a steady decline in services outside the protected area. The two non-linear patterns describe more complex dynamics: Type D (’Boundary Depression’) signifies a sharp drop in provision at the interface, while Type E (’Interface Peak’) represents a zone of enhanced provision at the border. This five-category typology serves as the basis for diagnosing the state of the borders and targeting interventions in the Technical scenario. Additional details of these categories are explained in the section 2.4 in the methods and in the Supplementary Information D, Figure S3, S4 and S5.

### 2.3 Future scenario co-development

We developed the ‘Participatory’ and ‘Technical’ scenarios through a multi-step process integrating expert-based diagnostics with local stakeholder knowledge. The scenarios are based on a common set of co-selected landscape interventions, but differ in their spatial allocation strategy.

The process began with two participatory workshops involving 16 local stakeholders in the first workshop and 10 in the second representing protected areas, agriculture, forestry, and local government and research (see Supplementary Information E for a detailed list). For the first workshop, we provided a preliminary diagnosis based on our baseline assessment, presenting maps that highlighted areas where ecosystem services bundles values fell within the lowest quintile (20%). These maps, alongside detailed land-use maps, served as the basis for discussion (see Supplementary Information E, Fig. S6). Working in three regional sub-groups, stakeholders identified key landscape challenges and opportunities. They then delineated polygons directly onto the land-use maps, proposing specific interventions to enhance ecosystem services provision and ecological connectivity. These interventions included diversifying some agricultural areas, restoring riverbanks, or creating ecological corridors between habitat patches.

Proposed interventions from the first workshop were translated into a set of spatially-explicit, algorithmic land-use change rules (see Supplementary Information E-a). In a second workshop, we presented these rules back to the stakeholders for validation and refinement. This collective review led to crucial adjustments to the rules’ technical parameters. For instance, the proposed width of hedgerows was reduced from 10m to 5m to better reflect on-the-ground constraints to agriculture (see Supplementary Information E, Fig S8). Stakeholders also refined the boundaries of the intervention polygons. Following this validation, we consolidated all proposed actions into five main management action groups: (1) Agricultural Diversification, (2) Ecological Corridors, (3) Forest Restoration & Management, (4) Urban Greening, and (5) River and Wetland Restoration. For each of these groups, stakeholders performed a feasibility assessment. Working in two sub-groups, participants first reached an internal consensus on an overall implementation score from 1 (Easy) to 5 (Hard). The final scores were then calculated as the average of the two sub-groups’ consensus scores. They listed key financial, social, knowledge, and governance factors underpinning these scores, rating each on a scale from 1 (strong barrier) to 5 (strong enabler/lever) (see Supplementary Information F for details). These factors were selected based on previous analyses in the region where stakeholders analysed barriers and levers for scaling NbS implementation (Bruley et al., 2025).

### 2.4 Scenario implementation and assessment

We used the refined set of intervention rules to model the two future scenarios. For the Participatory scenario, the rules were applied exclusively within the final priority polygons co-defined with the stakeholders (see Supplementary Information E, Fig. S7). For the Technical scenario, the same rule set was applied, but intervention locations were determined by the gradient analysis rather than by stakeholder polygons. We first selected all border segments classified as ‘Decreasing Gradient’ (Type C) or ‘Boundary Depression’ (Type D), where ecosystem service provision declines or drops sharply across the interface. Within the exterior structural buffer of each selected segment, we then assigned the intervention from the shared, stakeholder-validated rule set that matched the local land-use configuration, examining the underlying land-cover maps to determine which action was applicable at each location (for example, agricultural diversification where the buffer was dominated by cropland, or riverbank restoration along watercourses). These segments were prioritized to test whether interventions could mitigate the decline in ecosystem service provision from the interior to the exterior of protected areas, thereby extending their functional influence into the surrounding matrix.

To assess the impact of these scenarios, we re-evaluated the full suite of ecosystem services using the simulated future land-use maps. For most ecosystem services, whose provision is directly linked to land-use cover, the original models were applied to the new landscape configurations. The food production model, however, required a reparameterisation. Interventions such as agricultural diversification imply a shift towards agroecological practices that directly influence crop yields. We therefore adjusted the model’s yield parameters, ensuring our assessment captured changes in both land cover and land management intensity. A detailed explanation is provided in Supplementary Information E.d, Table S9.

### 2.5 Comparative analysis framework

To compare the impacts of the Participatory and Technical scenarios against the Present baseline, we implemented a multi-faceted analytical framework.

First, we quantified the spatial extent and leverage of ecosystem service changes relative to the planning effort. We calculated the the directly modified area (A_int_) as the percentage of the landscape where land-use categories changed. We then compared future provision maps against the baseline to quantify the percentage of the landscape showing ecosystem service improvement (A_imp_) or decline (A_dec_). From these values, we derived two Spatial Leverage Multipliers. The Positive Multiplier (M_pos_) represents the ratio between the area of service improvement and the direct area of land-use intervention (M_pos_=A_imp_/A_int_), quantifying the amplification of benefits. Conversely, the Negative Multiplier (M_neg_) represents the ratio of service decline to intervention area (M_neg_ = A_dec_/A_int_). A multiplier significantly greater than 1 indicates a leverage effect driven by landscape connectivity mechanisms.

Second, to understand how interventions altered the functional structure of the landscape, we used transition matrices to quantify the number of border segments that shifted between pattern types (e.g., from ‘Boundary Depression’ to ‘Decreasing Gradient’). For segments that maintained their classification, we analyzed quantitative changes in their gradient shape by extracting the polynomial regression coefficients before and after intervention. The statistical significance of these changes was assessed using paired t-tests or Wilcoxon signed-rank tests, depending on data normality, and linked to landscape configuration metrics (Supplementary Information G). Finally, to determine the biophysical mechanism driving these changes— specifically, to distinguish whether these quantitative modulations were driven by filling provision gaps or lowering internal values—we performed an absolute provision analysis for all segments that maintained their initial pattern classification. We calculated the mean normalized provision value (scale 0–1) for each structural buffer and computed the net differential (Δ) between future scenarios and the baseline across three functional zones: the protected area interior (Δ_Interior_), to assess integrity; the immediate border interface (Δ_Border_), to quantify connectivity improvements; and the distant matrix (Δ_Exterior_), to measure the spatial propagation of benefits.

Third, we integrated the quantitative outcomes of the scenarios with the qualitative data from the stakeholder workshops to analyze the trade-off between effectiveness and feasibility. We utilized the consensus difficulty scores (D_i_) assigned by stakeholders to each management action group on a scale from 1 (easiest) to 5 (most difficult). To compare the overall “social friction” of the strategies, we calculated an Area-Weighted Implementation Difficulty Score (S_w_) for each scenario using the formula:

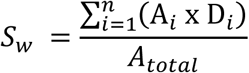

Where Ai is the area allocated to action group I and A_total_ is the total intervention area. We further decomposed this score to calculate the Contribution (C_i_) of each action group to the total landscape difficulty (C_i_ = (A_i_ × D_i_)/A_total_). This decomposition allows for identifying specific interventions that drive implementation risk. This integrated analysis allowed us to contextualize the biophysical outcomes of each scenario by explicitly quantifying the implementation effort required by their respective land-use configurations.

## 3. Results

### 3.1. Overall scenario impacts on ecosystem service provision and extent of change

The stakeholder-driven Participatory and the expert-driven Technical scenarios induced distinct changes in ecosystem service provision (Fig. 2A). Neither strategy was universally superior. The Technical scenario was nearly twice as effective at enhancing Pollination (Technical: +3.04%, Participatory: +1.67%), but resulted in larger declines in Food Production (Technical: -2.03%, Participatory: -1.34%) and Biological Control (Technical: -2.3%, Participatory: -0.3%). Conversely, both scenarios delivered comparable gains in Carbon Storage (Technical: +4.8%, Participatory: +4.8%) and substantially improved Outdoor Recreation, although the Technical approach yielded higher gains (Technical: +12.5%, Participatory: +4.9%). Heatwave Mitigation decreased slightly in both scenarios, driven by the forest thinning included in the intervention set.

**Figure 2.**
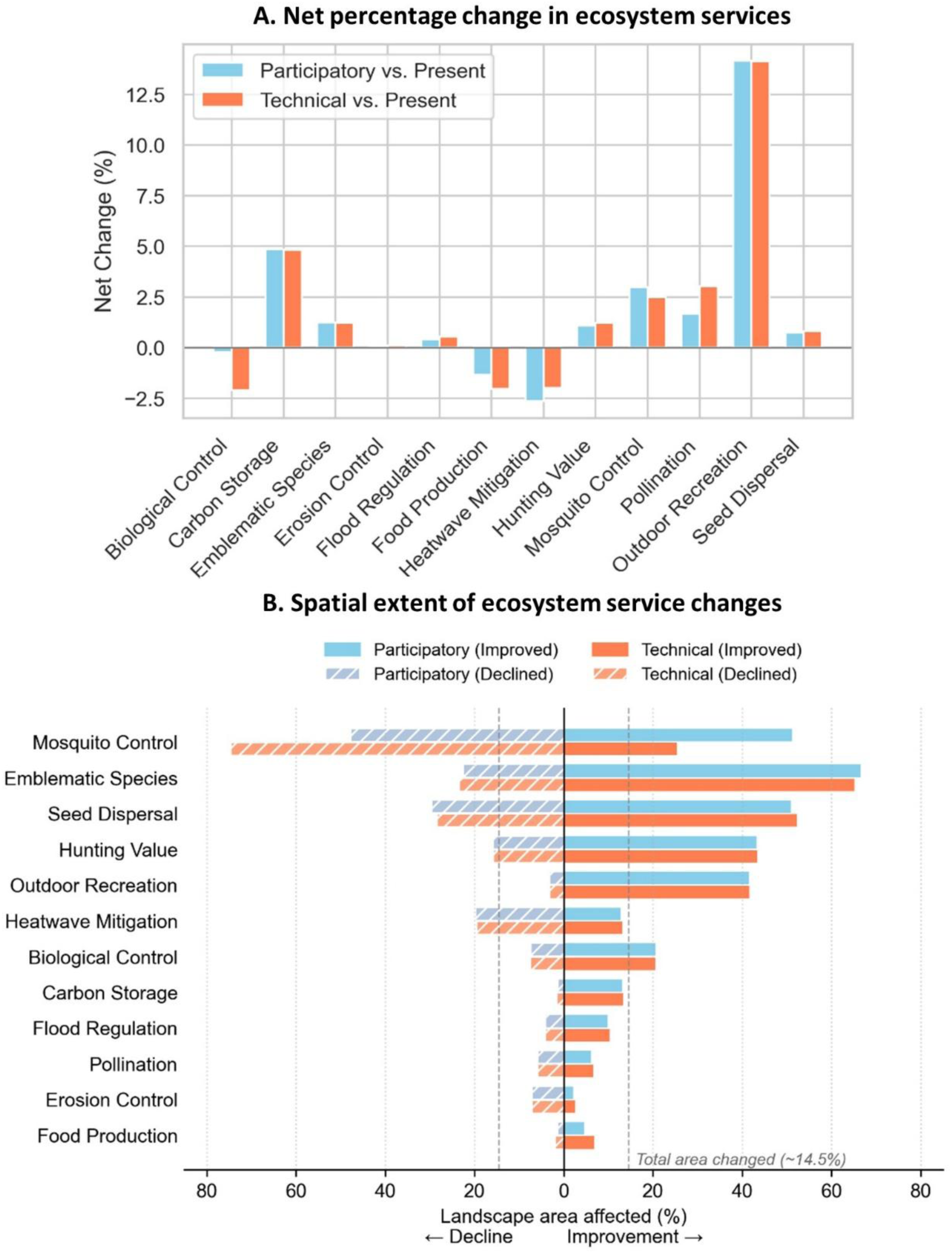
Net change and spatial extent of ecosystem service provision under future scenarios. (A) Net percentage change in the total landscape supply of 12 ecosystem services for the Participatory and Technical scenarios relative to the Present baseline, based on raw ecosystem service values prior to normalization. Positive values indicate a net gain in ecosystem service provision across the study area, while negative values indicate a net loss. (B) Spatial extent of ecosystem service changes. The bar chart shows the percentage of the total landscape area where ecosystem service provision either improved (solid bars) or declined (striped bars) for each scenario. The dashed vertical lines represent the total area directly modified by land-use change (∼14.5%). Ecosystem services with bars extending significantly beyond this threshold (e.g., Emblematic Species, Mosquito Control) demonstrate a high spatial propagation effect driven by connectivity mechanisms, whereas ecosystem services confined within the threshold (e.g., Carbon Storage) reflect impacts localized to the intervention footprint. Details on the specific spatial multipliers are provided in Supplementary Information Table S11.

The analysis of spatial leverage reveals that ecosystem service impacts propagated differently across the landscape (Fig. 2B, Supplementary Information G, Table S11). Although land-use interventions physically modified approximately 14.5% of the landscape in both scenarios, connectivity-dependent ecosystem services showed much broader spatial responses. Mosquito Control exhibited the most extensive declines. In the Technical scenario, for instance, declines propagated across 75% of the landscape (representing a 5-fold negative multiplier), whereas improvements were limited to 25% of the area (a 1.7-fold positive multiplier). Conversely, in both scenarios, Emblematic Species showed the most extensive improvements, with gains covering more than 65% of the landscape (a positive multiplier of ∼4.5) significantly outweighing declines. Seed Dispersal and Hunting Value also displayed high responsiveness across both scenarios, with spatial improvements covering between 43% and 52% of the territory (positive multipliers ranging from ∼3.0 to 3.5). Outdoor Recreation benefited substantially with minimal losses in both strategies (showing a positive multiplier near 2.9 versus a negative multiplier of 0.2). In contrast, Heatwave Mitigation showed a net spatial loss in both scenarios, with the area of decline (negative multiplier ∼1.3) exceeding the area of improvement (positive multiplier ∼0.9). Ecosystem services tied to biophysical stocks, such as Carbon Storage and Flood Regulation, tracked the directly modified area closely, showing balanced but limited propagation (multipliers ≤ 1.0). Finally, changes in Pollination and Erosion Control were confined to an area smaller than the total directly modified area (multipliers < 1.0 in both scenarios). This localized response occurs because these services depend strongly on specific crop interfaces or soil characteristics, which restricts their spatial propagation.

### 3.2. Sensitivity of border patterns at the individual service level

Gradients of ecosystem services dependent on landscape configuration proved sensitive to land-use changes (Table 2). Outdoor Recreation was the most responsive, with 22.1% of border segments changing pattern type. These transitions consistently involved smoothing abrupt patterns, specifically shifting from ‘Boundary Depression’ (Type D) to ‘Decreasing Gradient’ (Type C). Spatially, these transitions coincided with areas targeted for landscape diversification actions, such as adding hedgerows and green spaces, which enhance attractiveness near the border of protected areas. Mosquito Control and Pollination were also highly sensitive, frequently transitioning away from abrupt Type D patterns. Biological Control showed a distinct signature, with Type D patterns transitioning towards increasing or peaked gradients (Types B and E), indicating that agricultural interventions for this ecosystem service generate a complex restructuring of the gradient shape rather than simple smoothing.

**Table 2.** Sensitivity of border patterns at the individual ecosystem service level. The table shows the percentage of border segments (out of 104 total) that transitioned to a different category for each ecosystem service under the two future scenarios. Key changes (e.g., transitions from D to C, or C to A) are detailed to indicate the nature of the change.

| Bundle | Ecosystem Service | % Changed Tech | Main Improvement Tech | % Changed Participatory | Main Improvement Participatory |
| --- | --- | --- | --- | --- | --- |
| Cultural | Emblematic Species | 8.7 | D→C (4) | 7.7 | D→C (3) |
|  | Outdoor Recreation | 22.1 | D→C (13) C→A (1) | 22.1 | D→C (13) C→A (1) |
|  | Seed Dispersal | 8.7 | D→C (6) C→A (1) | 4.8 | D→C (2) C→A (1) |
| Rural | Biological Control | 8.7 | D→B (5) D→E (2) | 6.7 | D→C (1) C→A (1) |
|  | Carbon Storage | 5.8 | D→C (4) | 1.9 | D→C (1) |
|  | Food Production | 4.8 | D→C (1) | 2.9 | D→C (1) |
|  | Hunting Value | 4.8 | D→C (1) | 2.9 | D→C (1) C→A (1) |
|  | Pollination | 16.3 | D→C (5) | 9.6 | D→C (3) |
| Urban | Erosion Control | 1.9 | A→B (1) | 1 | D→C (1) |
|  | Flood Regulation | 4.8 | D→C (3) | 2.9 | D→C (1) |
|  | Heatwave Mitigation | 14.4 | D→C (5) | 10.6 | D→C (1) C→A (1) |
|  | Mosquito Control | 21.1 | D→C (1) C→A (2) | 7.4 | D→C (1) |

In contrast, ecosystem services tied to biophysical stocks remained highly stable. Erosion Control was the most stable ecosystem service, with fewer than 2% of segments transitioning in either scenario. Carbon Storage and Food Production also demonstrated high stability, with change rates below 6%. Even when transitions occurred, they were limited; for instance, the Technical scenario induced a shift away from the Type D pattern in only 4 of its 27 initial occurrences for Carbon Storage.

### 3.3. Transformation of border patterns for ecosystem service bundles

At the bundle level, border patterns showed high overall stability, with over 90% of segments maintaining their initial classification in both scenarios (Fig. 3). For the 10% remaining segments, the Technical scenario smoothed 46% of ‘Boundary Depression’ (Type D) patterns into ‘Decreasing Gradients’ (Type C), compared to 26% in the Participatory scenario. This smoothing was driven by distinct landscape mechanisms depending on the bundle. For the Rural bundle, the transition corresponded to a targeted increase in natural habitat immediately outside the protected area, whereas for the Cultural bundle, the D→C transition was associated with a widespread increase in landscape fragmentation (decrease in Moran’s I) across the border interface (Supplementary Information G, Fig. S11).

**Figure 3.**
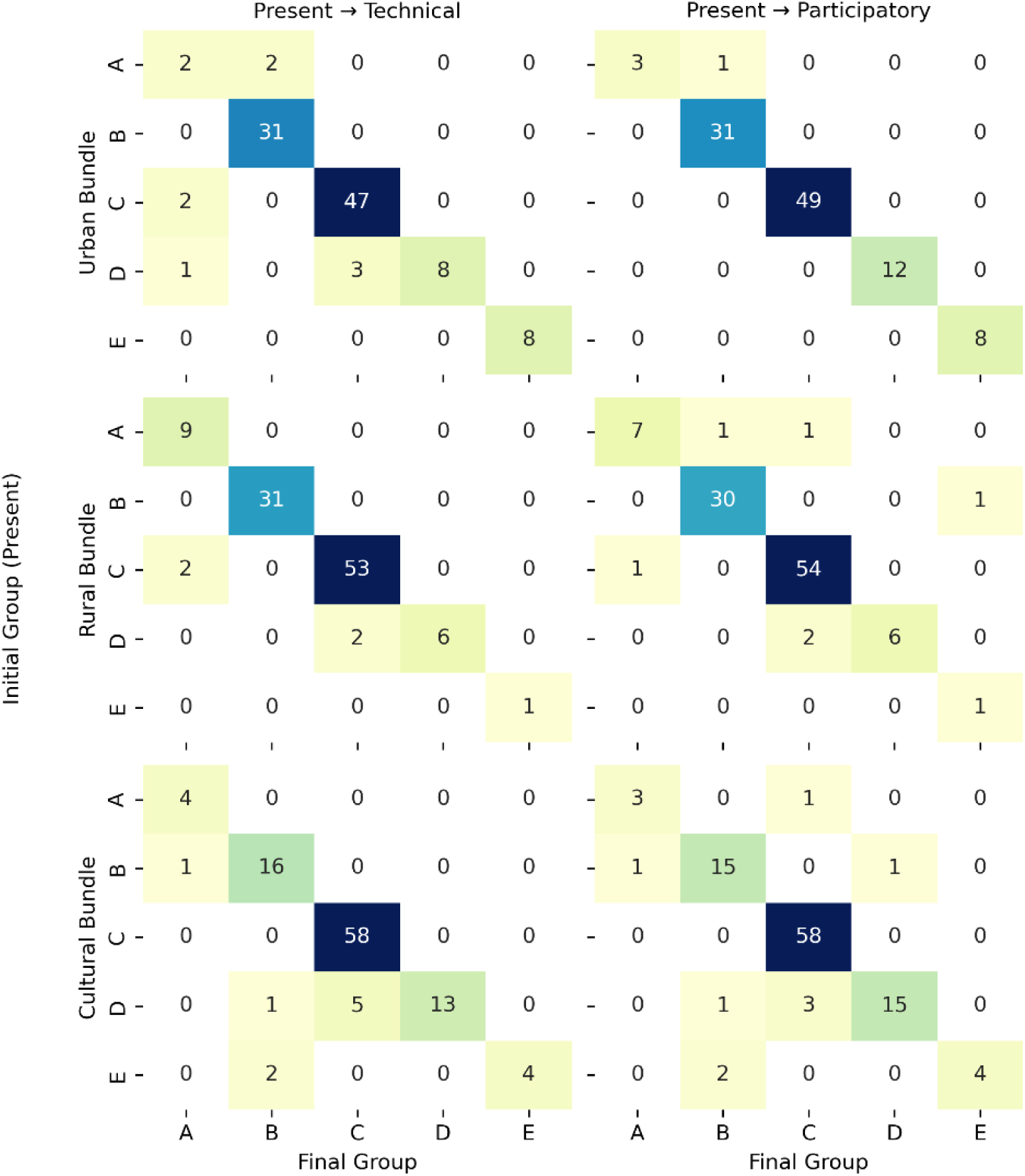
Transformation of aggregated border patterns. Transition matrices showing the number of border segments that transitioned between pattern types (A-E) from the Present baseline (rows) to the final state in the Technical (left) and Participatory (right) scenarios (columns). The diagonal (darker cells) represents the number of segments that remained stable in their initial category. Off-diagonal cells indicate categorical transformations, highlighting the distinct impacts of each scenario on the landscape’s structural patterns. The total number of segments analysed was 104 for each bundle.

The Technical scenario transitioned six ‘Decreasing Gradient’ (Type C) segments to ‘Seamless Transfer’ (Type A) patterns, an outcome not achieved by the Participatory approach. This transition was mechanistically tied to a sustained increase in landscape diversity (Shannon’s Diversity Index) across the matrix outside the protected area, primarily in the Rural bundle (Supplementary Information G, Fig. S12). These differences highlight the distinct structural impacts of each scenario. The Technical scenario induced a greater number and diversity of transitions (29 total, comprising D→C, D→B, and C→A shifts). In contrast, the 18 transitions in the Participatory scenario were almost exclusively concentrated on the D→C smoothing mechanism. This indicates that with an additional 0.5% of the landscape directly modified, the Technical scenario produced substantially more transitions and qualitatively distinct outcomes, such as the creation of Type A patterns. This suggests that a small increase in the directly modified area can enable a more substantial reorganization of landscape patterns, a finding we further explore in the context of implementation feasibility.

### 3.4. Modification of gradient shapes and absolute provision

Land-use interventions modified the profile of ecosystem service gradients, affecting both their geometric shape and absolute provision levels (Fig. S9 and S10; Fig. 4). For unchanged ‘Decreasing Gradients’ (Type C), the Technical scenario made the decline significantly shallower across all bundles (p < 0.001). The normalized profiles also showed a lower intercept (p < 0.01) (Supplementary Information G, Table S12), but the absolute profiles confirm that this smoothing came from an increase in ecosystem service provision across the agricultural matrix, not from a reduction inside the protected area (Fig. 4I and 4II). For the Rural bundle, this corresponds to a significant increase in landscape diversity (Shannon’s H) in the matrix outside the protected area. For the Cultural and Urban bundles, the same outcome was associated with a widespread increase in landscape fragmentation (reduction in Moran’s I) (Supplementary Information G, Fig. S13, Table S14). In contrast, the Participatory scenario also decreased the slope for the Cultural and Urban bundles (p < 0.01) but showed a mixed effect on the intercept, with no consistent statistical pattern.

**Figure 4.**
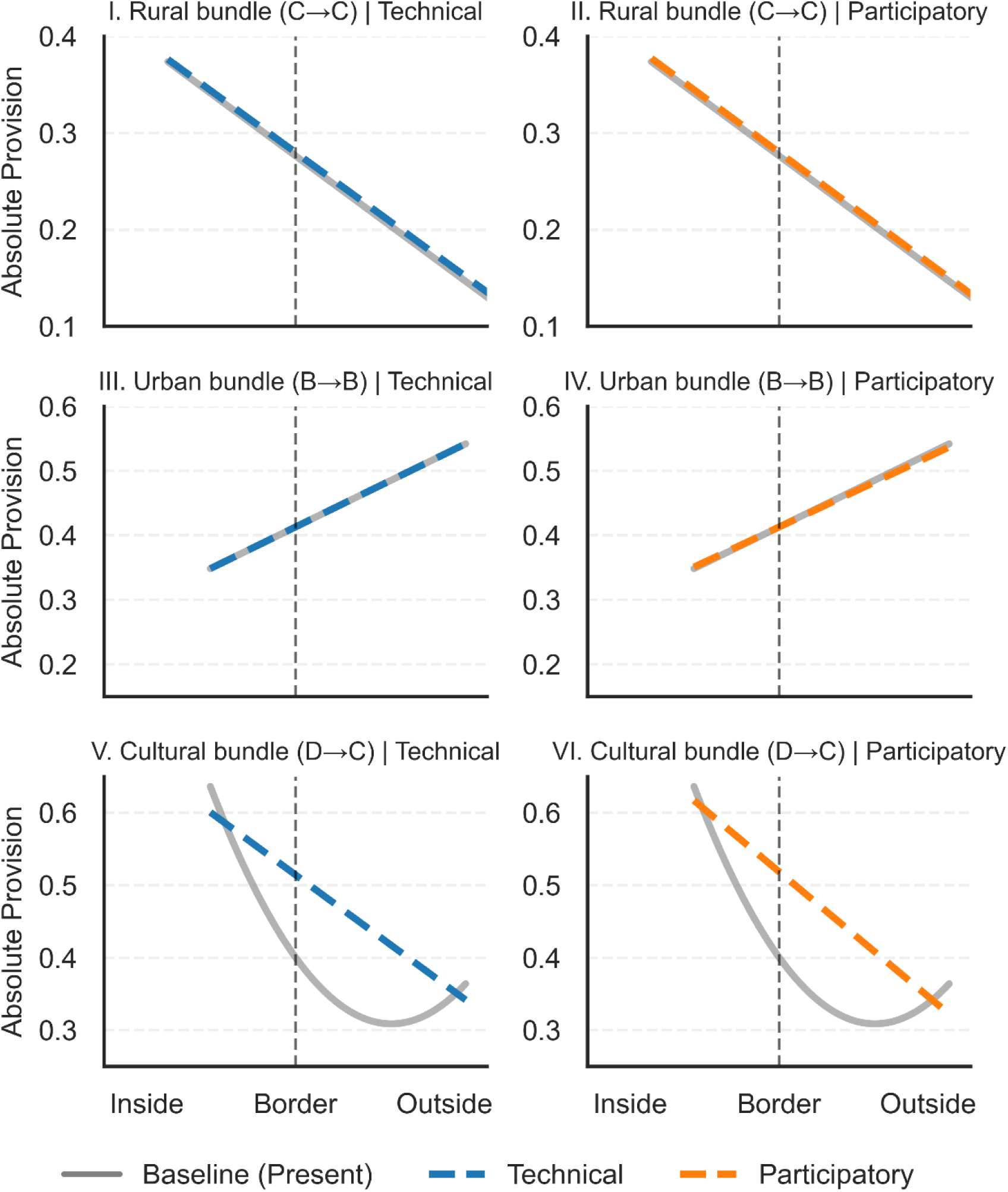
Biophysical signature of landscape interventions on ecosystem service gradients. The plots display the mean absolute ecosystem service provision (prior to the 0-1 normalization) across the protected area interface for three representative gradient transitions selected based on the Technical scenario classification (see Table S16 for full quantitative details). The x-axis denotes the distance in structural buffers, where 0 represents the protected area border. Solid grey lines indicate the Baseline; dashed colored lines indicate the Technical (blue) and Participatory (orange) scenarios. The panels illustrate three distinct biophysical mechanisms: (Top) For Rural ‘Decreasing Gradients’ (N=53), interventions induce a slight, statistically significant moderation of the slope, although absolute provision changes remain visually subtle. (Middle) Similarly, for Urban ‘Increasing Gradients’ (N=31), absolute provision levels show minimal visual deviation from the baseline, reflecting a highly stable magnitude despite significant statistical modulations of the gradient’s shape. (Bottom) For Cultural ‘Boundary Depressions’ (N=5), the Technical scenario successfully eliminates the boundary discontinuity by effectively filling the provision gap at the border, whereas the Participatory scenario (which achieved this transition in only N=3 cases, see Table S16) results in lower absolute provision at the interface for the plotted segments.

For consistent ‘Increasing Gradients’ (Type B), both scenarios resulted in a similar moderating effect, significantly reducing the slope and increasing the intercept for the Urban bundle (p < 0.05) (Supplementary Information G, Table S13). However, unlike the Rural bundle, the absolute profiles show that ecosystem services provision remained closely aligned with the baseline, with minimal deviation in total magnitude (Fig. 4III and 4IV). In terms of landscape pattern, this stability was associated with an increase in fragmentation (significant decrease in Moran’s I) immediately outside the protected area, a pattern confirmed for both the Cultural and Urban bundles (Supplementary Information G, Fig. S14, Table S15).

In contrast to the similar responses observed for Type B gradients, a sharp divergence between scenarios emerged in non-linear gradients, particularly for ‘Boundary Depression’ patterns (Type D) (Fig. 4V and 4VI). While the normalized analysis indicated that both scenarios flattened the valley shape, the absolute profiles demonstrate a sharp contrast in effectiveness. The Technical scenario achieved a structural transition to a ‘Decreasing Gradient’ (Type C) by substantially increasing ecosystem services provision at the border interface, effectively bridging the gap between the protected area and the matrix. Conversely, the Participatory scenario reduced the depth of the valley but maintained the original depression shape, resulting in lower absolute provision at the interface. This difference in magnitude explains why the Technical scenario was substantially more effective at generating categorical transitions (Table 2), successfully eliminating the boundary discontinuity through targeted changes in landscape configuration, whereas the Participatory approach limited its impact to gradient smoothing.

### 3.5. The effectiveness-feasibility trade-off

Our analysis reveals a fundamental tension between biophysical effectiveness and implementation feasibility (Fig. 5). While both scenarios intervene on a significant portion of the landscape (>14%), this area is unevenly distributed. The vast majority (>52,000 ha) corresponds to a shared forest adaptation strategy designed to diversify tree species composition. This action, primarily affecting public lands, was deemed moderately feasible by stakeholders (score 2.5/5), as it aligns with institutional mandates for climate resilience. Consequently, because forest interventions dominate the total area in both scenarios, the global Area-Weighted Difficulty Scores remain quantitatively similar (Technical: 2.83 vs. Participatory: 2.72).

**Figure 5.**
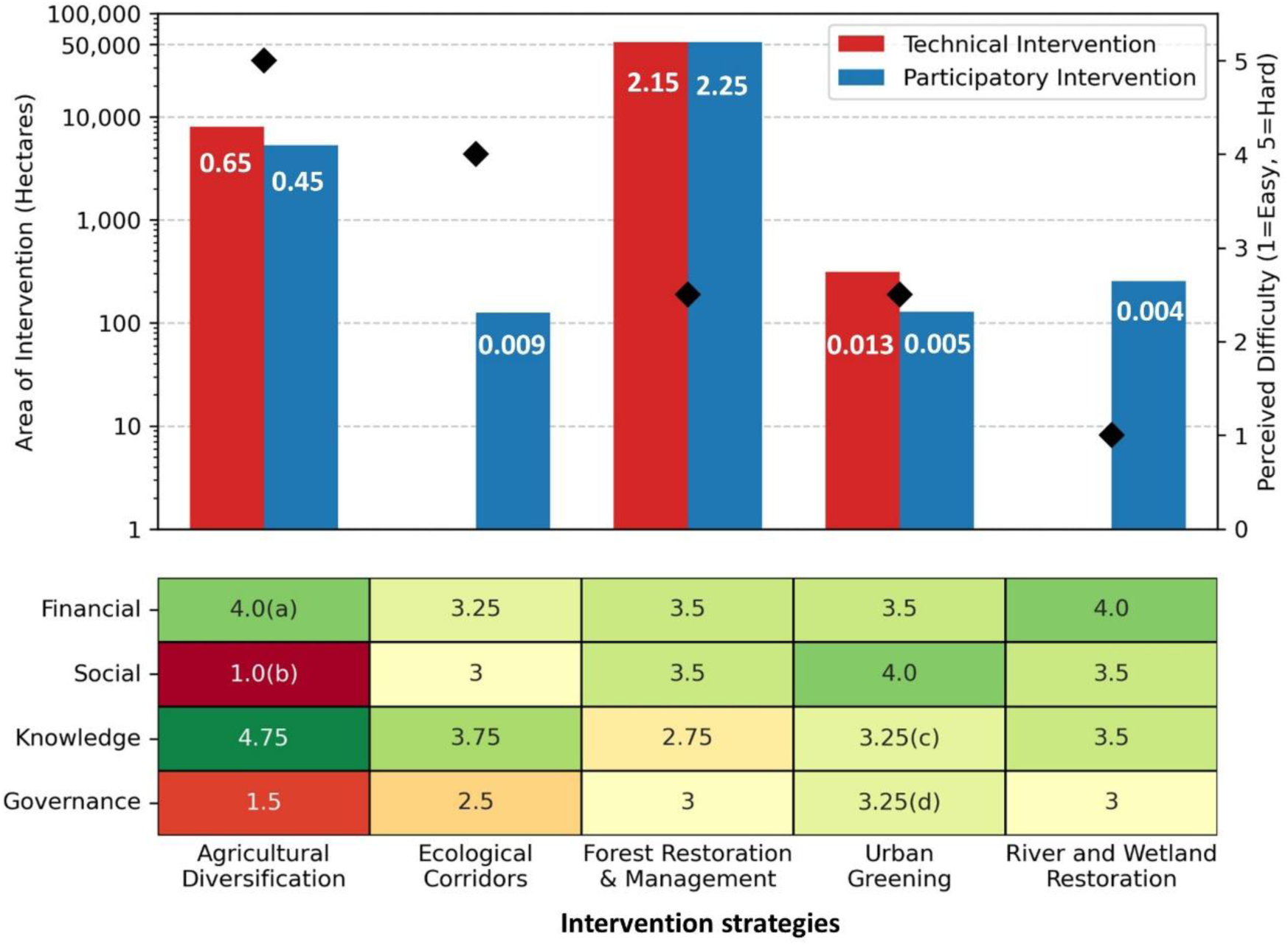
Intervention strategy versus implementation feasibility. The bar chart shows the total area (in hectares, logarithmic scale) allocated to each of the five management action groups under the Technical (red) and Participatory (blue) scenarios. Black diamonds indicate the mean perceived implementation difficulty for each action group, as ranked by stakeholders (1=Easy, 5=Hard). The white numbers on the bars represent the weighted contribution (Ci) of each action to the total scenario difficulty score, calculated as the product of the modified area and its difficulty ranking, normalized by total landscape area. The heatmap below deconstructs the difficulty score into four key feasibility domains (Financial, Social, Knowledge, Governance), where green indicates enabling factors (levers) and red indicates significant barriers. (a) Stakeholders noted that while financial support for agriculture is high, it can fund opposing goals, acting as both a lever and a barrier. (b) Social acceptance for agroecology is highly polarized, depending on the interest group. (c) Urban greening was described as complex, with available funding but implementation challenges due to space limitations and urban trade-offs. (d) Governance for urban greening was also noted as a potential barrier, as certain sectors (e.g., public health) may perceive an excess of green space as unsafe due to pest concerns.

However, decomposing these scores reveals critical divergences in the management of the non-forest matrix. The Technical scenario, guided by gradient analysis, focused almost exclusively on addressing discontinuities at the protected area interface. As a result, it allocated over 99% of its non-forest intervention area to Agricultural Diversification. While agroecological practices effectively restructured landscape patterns, stakeholders ranked them as the most difficult to implement (mean difficulty: 5/5), citing governance constraints and social barriers. This specific intervention contributed 0.65 points to the total difficulty score in the Technical scenario.

In contrast, the Participatory scenario, shaped by social license, reduced the specific contribution of agricultural interventions to 0.45 points. Instead, it allocated a significant portion of its non-forest effort to River and Wetland Restoration. This action was ranked as the easiest to implement (1/5) due to funding availability and lower social resistance. Thus, while the Technical scenario relies heavily on the most difficult interventions to achieve its structural outcomes, the Participatory scenario achieves a more balanced risk profile by incorporating socially preferred, lower-difficulty actions, even if they address fewer structural discontinuities identified by the gradient analysis.

## 4. Discussion

### 4.1 Landscape-scale assessment of nature-based solutions beyond quantitative targets

Our results indicate that evaluating nature-based solutions based solely on aggregate changes in ecosystem service supply can underestimate their structural impact. While stock-based ecosystem services such as Carbon Storage showed modest net gains (∼4.8%) tracking the area directly modified by the scenarios, connectivity-dependent ecosystem services exhibited widespread responses. This distinction aligns with the framework proposed by Metzger et al. (2021), who emphasize differentiating between impacts derived from the total area modified and those resulting from its spatial arrangement. Similarly, recent studies demonstrate that landscape configuration differentially modulates the provision of multiple ecosystem services (Lamy et al., 2016; Neyret et al., 2025), informing the design of multifunctional landscape templates where spatial pattern can mitigate trade-offs (Lavorel et al., 2022).

This perspective is particularly relevant for ecosystem services dependent on landscape connectivity. In our Technical scenario, land-use interventions facilitated a widespread improvement in habitat suitability for Emblematic Species. This supports the argument that the spatial arrangement of natural elements is a primary driver for organism mobility (Mitchell et al., 2015). By analyzing the interface between protected areas and their surroundings, we observed that specific interventions, such as agricultural diversification and hedgerow creation, smoothed abrupt ‘Boundary Depression’ gradients into more gradual transitions. This is consistent with evidence that woody elements added to the agricultural matrix improve functional connectivity only when strategically placed, for instance as stepping stones along river corridors, rather than through a simple increase in wooded area (Pineda-Zapata et al., 2026).

Our landscape metric analysis reveals the mechanisms behind these changes. The transformation of gradient shapes was associated with a localized increase in landscape diversity (Shannon Index) and heterogeneity (reduction in Moran’s I) immediately outside protected area boundaries. A marked spatial propagation emerged for connectivity-based ecosystem services. For instance, interventions modified ∼14.5% of the landscape but induced changes in Emblematic Species provision across more than 65% of the territory. While the net increase in supply was moderate, this widespread spatial response indicates a structural reorganization that effectively reduced the contrast between the protected area and the adjacent matrix. These findings reinforce the role of protected areas as ecological cornerstones that require effective integration with working landscapes (Kremen and Merenlender, 2018). While area-based targets such as those in the Nature Restoration Regulation remain important, our findings suggest that where interventions are placed can matter as much as how much area is restored, particularly for connectivity-dependent ecosystem services (Pineda-Zapata et al., 2026). This implies that restoration planning elsewhere would benefit from assessing structural, configuration-level outcomes alongside aggregate area or supply targets, rather than either in isolation.

### 4.2 Navigating trade-offs in conflict-prone agricultural and urban interfaces

Our comparative analysis reveals a sharp contrast in the implementation potential of NbS defined by a dichotomy between spatial availability and social feasibility. In the agricultural matrix, the Technical scenario identified a substantial biophysical potential. Excluding the large-scale forest management changes common to both scenarios, this scenario directed over 99% of its remaining effort toward agricultural diversification. This indicates that productive landscapes offer sufficient space to restore connectivity and support the continuity of ecosystem service provision (Grass et al., 2019). However, this potential clashes with socio-political reality, as stakeholders ranked these interventions as the most difficult to implement (score 5/5), citing conflicts with production models. This resistance aligns with findings that farmers often hesitate to adopt agroforestry due to tradition, lack of technical knowledge, and subsidy eligibility concerns (Rois-Díaz et al., 2018; Dubo et al., 2025; Tobin et al., 2026). Furthermore, this tension reflects the structural rigidity of the Common Agricultural Policy, which has struggled to incentivize biodiversity conservation on working lands (Pe’er et al., 2020). Our results reinforce the argument that technical prioritization alone is insufficient to overcome systemic barriers rooted in the competition between biodiversity targets and economic land-use demands (Chapman et al., 2025).

The urban interface presents an inverse challenge, where the binding constraint is not social acceptance but physical saturation. In dense urban fabrics, competition for space and the rigidity of the built environment restrict opportunities for retrofitting nature, limiting the scalability of NbS even where political will is strong (Diep and McPhearson, 2025; McPhearson et al., 2025). Our case illustrates this general limit: despite ambitious greening rules designed to maximize connectivity, the model modified only a negligible fraction of the landscape compared with the rural context, and urban ecosystem service gradients remained largely stable. Concretely, our scenarios added over 100 hectares of urban tree planting, less than a third of the roughly 370 hectares that Grenoble’s ‘Plan Canopée’ targets by 2030 to reach a 30% canopy cover.

However, the small area directly modified by urban interventions should not be conflated with limited impact. While our biophysical models show modest changes in total ecosystem service supply, literature suggests that in densely populated areas even small-scale interventions yield substantial social and health benefits due to the high density of beneficiaries (Cook et al., 2025; Raymond et al., 2017). Therefore, while agricultural restoration requires overcoming governance barriers to unlock its spatial potential, urban restoration requires targeted retrofitting. These interventions must integrate functional diversity within built environment constraints to maximize co-benefits such as heat mitigation and social cohesion where they are most needed (González-García et al., 2025).

### 4.3 Balancing technical optimization with social legitimacy in landscape restoration

Our results highlight a tension between the biophysical efficiency of technical planning and the social legitimacy required for implementation. The Technical scenario was more effective at restoring the structural continuity of the landscape. By targeting specific biophysical deficits, this approach smoothed 46% of the abrupt ‘Boundary Depression’ (Type D) gradients compared to 26% in the Participatory scenario. Furthermore, it uniquely generated ‘Seamless Transfer’ (Type A) patterns, eliminating the border discontinuity between protected areas and the matrix. Notably, these structural changes were achieved with a marginal increase in the directly modified area (0.5% of the landscape), aligning with literature proposing that spatial optimization effectively identifies high-leverage interventions to maximize ecological outcomes (Albert et al., 2019).

However, relying exclusively on technical criteria risks creating an implementation gap. While ecologically robust, the Technical scenario assumes land is a neutral substrate, overlooking the plural values local communities hold and the situated, place-based realities in which restoration is implemented (Pascual et al., 2017; Hall and van Rees, 2026). As noted by Diep and McPhearson (2025), technical approaches often fail to account for local capacities and power dynamics. In our study, stakeholders constrained the Participatory scenario by favoring actions with higher feasibility scores, reflecting a preference for interventions aligned with existing social norms even if they result in less transformative ecological outcomes. This creates a risk that technical plans remain theoretical if they lack the social support to navigate local conflicts (Andersson et al., 2025).

Furthermore, a purely technical approach raises concerns regarding procedural justice. Prioritizing restoration targets based solely on biophysical models can inadvertently marginalize local needs or place disproportionate burdens on specific groups, such as farmers in the agricultural matrix (Woroniecki et al., 2019; Rois-Díaz et al., 2018). Conversely, relying entirely on participatory processes carries limitations. Stakeholder-driven priorities in our study concentrated on areas with perceived local relevance, leading to a dispersed restoration pattern less aligned with key connectivity bottlenecks. This echoes the warning by Chapman et al. (2025) regarding “naive prioritization,” where planning lacking a strategic landscape perspective fails to achieve necessary biodiversity targets.

Therefore, mainstreaming NbS requires an iterative integration of both approaches. Technical diagnostics should serve as a navigational tool to inform stakeholders of critical deficits and leverage points rather than as a rigid prescription (Metzger et al., 2021). This enables knowledge co-production where technical precision guides discussion while social values determine the final design (Norström et al., 2020). Treating technical models and participatory processes as complementary makes it possible to design interventions that are both ecologically significant and socially feasible.

### 4.4 Harmonizing sectoral policies to boost the functional role of protected areas

Our results suggest that international frameworks like the Global Biodiversity Framework and the EU Nature Restoration Regulation should function as mechanisms to coordinate sectoral policies rather than solely as conservation mandates. This is particularly pressing given evidence that human land-use pressure inside protected areas is projected to rise substantially in coming decades, threatening the very targets these frameworks set (Li et al., 2026). Our Technical scenario demonstrates that integrating forestry, urban, and agricultural planning is biophysically feasible. By modifying approximately 14.5% of the landscape, predominantly through forest adaptation and targeted interventions, it is possible to restore protected area boundary patterns across large portions of the territory. This potential is reinforced by recent findings identifying suboptimal cropland areas as strategic bridges where revegetation can support climate and biodiversity goals without compromising food security (Hua et al., 2025). Similarly, evidence from agroecological transitions in India shows that biodiversity recovery can be achieved while maintaining yields, creating mutual benefits for wildlife and farmers (Berger et al., 2025).

Stakeholders identified funding availability as a facilitator for interventions like river restoration, challenging the assumption that financial resources are the primary limiting factor (Bruley et al., 2025). Instead, the challenge lies in policy alignment. As noted by zu Ermgassen et al. (2025), conflicting public policies often hinder investment, specifically when agricultural subsidies compete with and crowd out nature-based investments. Furthermore, the Scenar 2040 of the Common Agricultural Policy report highlights that focusing exclusively on environmental measures can reduce production, whereas productivity-oriented approaches increase emissions (Fellmann et al., 2025). Our findings point to an intermediate path. If restoration policies align existing sectoral funds toward specific connectivity elements like hedgerows, environmental goals can be advanced with optimized land use. Adopting this integrated perspective allows for generating synergies across multiple management priorities, linking urban needs, agricultural permeability, and protected area integrity.

Importantly, the approach we used to build these scenarios is not specific to our study area. Because the interventions are expressed as explicit, raster-based land-use rules co-selected and validated with local stakeholders, and are allocated using a gradient diagnostic that can be computed for any protected area interface, the same procedure can be transferred to other regions and policy contexts. What changes between sites is the local set of actions and priorities; the diagnostic-to-scenario workflow remains the same. This makes the framework a practical template for aligning sectoral policies with landscape connectivity beyond the specific case presented here.

#### Conclusions

This study establishes gradient analysis as a diagnostic tool for integrating protected area benefits into landscape-scale restoration. By moving beyond aggregate area-based metrics, this approach reveals structural discontinuities at the boundaries of protected areas that standard assessments often overlook. Our results demonstrate that systemic improvement relies on the propagation of benefits rather than just the scale of intervention. While our scenarios modified approximately 14.5% of the landscape, primarily through necessary forest adaptation and targeted agricultural changes, this intervention generated widespread gains in connectivity, improving connectivity-dependent ecosystem service provision across up to 65% of the territory. This confirms that the effectiveness of NbS depends not only on the sheer quantity of restored hectares but also on their spatial configuration and their capacity to reconnect isolated conservation areas with the surrounding matrix.

Yet there is a risk that implementation lags behind potential. The areas with the highest potential to restore ecological integrity, especially productive agricultural landscapes, tend to be constrained by strong socio-economic lock-ins. The mismatch between biophysical potential and social feasibility indicates that technical diagnostics are essential for identifying where interventions would be most effective, but they cannot dictate how those changes can be socially and politically implemented. Ultimately, scaling up NbS requires shifting the focus from funding availability to policy coherence and value changes (Bruley et al., 2025). Unlocking the restoration potential on working lands demands that environmental frameworks such as the Nature Restoration Regulation act as coordinating mechanisms that align existing sectoral incentives, particularly in forestry and agriculture, with the creation of landscape complexity. Transforming protected areas from isolated refuges into interconnected nodes within a multifunctional landscape can help global biodiversity goals translate into resilient local outcomes.

## Supporting information

Appendix_Gonzalez_Garcia_et_al_2026_scenarios

## Acknowledgments

We warmly thank the stakeholders who took part in the workshops for their time, engagement and generosity; their knowledge of the territory was essential to this work. This research was supported by Project RECONNECT, Biodiversa+, the European Biodiversity Partnership under the 2021–2022 BiodivProtect joint call for research proposals, with funding to AGG, MCP and AGG by the ANR Agence Nationale de la Recherche (ANR-22-EBIP-0009-06). MN received funding from the European Union’s Horizon 2020 research and innovation programme under the Marie Skłodowska-Curie grant agreement No 101104374.

## Notes

### Competing Interest Statement

The authors have declared no competing interest.

## References

Albert, C., Schröter, B., Haase, D., Brillinger, M., Henze, J., Herrmann, S., … & Matzdorf, B. (2019). Addressing societal challenges through nature-based solutions: How can landscape planning and governance research contribute?. Landscape and urban planning, 182, 12–21.

Andersson, E., Martin, R., Anderson, P., Brooks, S., Cortés Capano, G., González-García, A., … Raymond, C. M. (2025). Social-ecological connections and landscape approaches in conservation. BioScience.

Berger, I., Kamble, A., Morton, O., Raj, V., Nair, S. R., Edwards, D. P., Wauchop[1]e, H. S., Joshi, V., Basu, P., Smith, B., & Dicks, L. V. (2025). India’s agroecology programme, ‘Zero Budget Natural Farming’, delivers biodiversity and economic b[1]enefits withou[1]t lowering yields. Nature Ecology & Evolution, 9, 2057–2068.

Bruley, E., Palomo, I., Lavorel, S., Locatelli, B., & Dubo, T. (2025). Leverage points for scaling nature-based adaptation to climate change. People and Nature, 7, 2743–2758.

Byczek, C., Longaretti, P. Y., Renaud, J., & Lavorel, S. (2018). Benefits of crowd-sourced GPS information for modelling the recreation ecosystem service. PloS one, 13(10), e0202645.

Chan, K. M. A., Balvanera, P., Benessaiah, K., Chapman, M., Díaz, S., Gómez-Baggethun, E.,… Klain, S. (2016). Opinion: why protect nature? Rethinking values and the environment. Proceedings of the National Academy of Sciences, 113(6), 1462–1465.

Chapman, M., Jung, M., Leclère, D., Boettiger, C., Augustynczik, A. L. D., Gusti, M., Ringwald, L., & Visconti, P. (2025). Meeting European Union biodiversity targets under future land-use demands. Nature Ecology & Evolution. 10.1038/s41559-025-02671-1

Cohen-Shacham, E., Andrade, A., Dalton, J., Dudley, N., Jones, M., Kumar, C.,… Walters, G. (2019). Core principles for successfully implementing and upscaling nature-based solutions. Environmental Science & Policy, 98, 20–29.

Cook, E. M., Kim, Y., Grimm, N. B., McPhearson, T., Anderson, P., Bulkeley, H., Collier, M. J., Diep, L., Morató, J., & Zhou, W. (2025). Nature-based solutions for urban sustainability. Proceedings of the National Academy of Sciences, 122(29), e2315909122. 10.1073/pnas.2315909122

Cousins, J. J. (2021). Justice in nature-based solutions: Research and pathways. Ecological economics, 180, 106874.

Díaz, S., Settele, J., Brondízio, E. S., Ngo, H. T., Agard, J., Arneth, A.,… Willis, K. J. (2019). Pervasive human-driven decline of life on Earth points to the need for transformative change. Science, 366(6471), eaax3100.

Diep, L., & McPhearson, T. (2025). Empowering cities globally: Four levers for transformative urban adaptation with nature-based solutions. Proceedings of the National Academy of Sciences, 122(29), e2315912121. 10.1073/pnas.2315912121

Dubo, T., Palomo, I., Zingraff-Hamed, A., Bruley, E., Collain, G., & Lavorel, S. (2023). Levers for transformative nature-based adaptation initiatives in the Alps. PLOS Climate, 2(11), e0000193.

Fellmann, T., Tassinari, G., Lasarte Lopez, J., Rey Vicario, D., Beber, C., Barbosa, A. L., De Jong, B., Ferrari, E., Gocht, A., Isbasoiu, A., Klinnert, A., Kremmydas, D., M’Barek, R., Philippidis, G., Rokicki, B., Tillie, P., Weiss, F., & Genovese, G. (2025). Scenar 2040: A scenario study on the Common Agricultural Policy. Publications Office of the European Union.

González-García A, Neyret M, López-Tejedor A, Prima MC, Si-Moussi S, Renaud J, Gueguen M, Lavorel S (2026) Ecosystem service gradients at protected area borders reveal multiple patterns and prevalent management conflicts. Conservation Biology.

González-García, A., Palomo, I., Codemo, A., Rodeghiero, M., Dubo, T., Vallet, A., & Lavorel, S. (2025). Co-benefits of nature-based solutions exceed the costs of implementation. Cell Reports Sustainability, 2(3).

Grass, I., Loos, J., Baensch, S., Batáry, P., Librán-Embid, F., Ficiciyan, A., … & Tscharntke, T. (2019). Land-sharing/-sparing connectivity landscapes for ecosystem services and biodiversity conservation. People and Nature, 1(2), 262–272.

Hall, D. M., & van Rees, C. B. (2026). Implementing nature-based solutions requires distinguishing place from space. Ambio, 1–18.

Herzon, I., Birge, T., Allen, B., Povellato, A., Vanni, F., Hart, K., … & Pražan, J. (2018). Time to look for evidence: Results-based approach to biodiversity conservation on farmland in Europe. Land use policy, 71, 347–354.

Hua, T., Hu, X., Austrheim, G., Speed, J. D., van Oort, B., & Cherubini, F. (2025). Reconciling crop production, climate action and nature conservation in Europe by agricultural intensification and extensification. Nature Communications, 16(1), 10289.

IPBES (2019). Global assessment report on biodiversity and ecosystem services of the Intergovernmental Science-Policy Platform on Biodiversity and Ecosystem Services, eds Brondizio ES, Settele J, Díaz S, Ngo HT (IPBES secretariat, Bonn, Germany), p 1148.

IPBES (2024). Thematic Assessment Report on the Interlinkages among Biodiversity, Water, Food and Health of the Intergovernmental Science-Policy Platform on Biodiversity and Ecosystem Services. Harrison, P. A., McElwee, P. D., and van Huysen, T. L. (eds.). IPBES secretariat, Bonn, Germany.

Keesstra, S., Nunes, J., Novara, A., Finger, D., Avelar, D., Kalantari, Z., & Cerdà, A. (2018). The superior effect of nature based solutions in land management for enhancing ecosystem services. Science of the Total Environment, 610-611, 997–1009.

Kremen, C., & Merenlender, A. M. (2018). Landscapes that work for biodiversity and people. Science, 362(6412), eaau6020.

Lamy, T., Liss, K. N., Gonzalez, A. & Bennett, E. M. Landscape structure affects the provision of multiple ecosystem services. Environ. Res. Lett. 11, 124017 (2016)

Lasseur R, Vannier C, Lefebvre J, Longaretti PY, Lavorel S (2018) Landscape-scale modeling of agricultural land use for the quantification of ecosystem services. J Appl Remote Sens 12, 046024.

Lavorel, S., Grigulis, K., Richards, D. R., Etherington, T. R., Law, R. M., & Herzig, A. (2022). Templates for multifunctional landscape design. Landscape Ecology, 37(3), 913–934.

Li, G., Wang, Z., Sun, S., Qi, W., & Watson, J. E. (2026). Projected human land-use pressures and natural habitat conversion risk within global terrestrial protected areas. Nature Ecology & Evolution, 10(2), 281–292.

Locatelli, B., Lavorel, S., Colloff, M. J., Crouzat, E., Bruley, E., Fedele, G.,… Walters, G. (2025). Intertwined people–nature relations are central to nature-based adaptation to climate change. Philosophical Transactions of the Royal Society B: Biological Sciences, 380, 20230213.

Martín-López, B., Gómez-Baggethun, E., García-Llorente, M., & Montes, C. (2014). Trade-offs across value-domains in ecosystem services assessment. Ecological Indicators, 37A, 220–228.

Martín-López, B., Iniesta-Arandia, I., García-Llorente, M., Palomo, I., Casado-Arzuaga, I., Del Amo, D. G.,… Montes, C. (2012). Uncovering ecosystem service bundles through social preferences. PLoS ONE, 7(6), e38970.

McPhearson, T., Frantzeskaki, N., Ossola, A., Diep, L., Anderson, P. M. L., Blatch, T., Collier, M. J., Cook, E. M., Culwick Fatti, C., Grabowski, Z. J., Grimm, N. B., Haase, D., Herreros-Cantis, P., Kavonic, J., Lin, B. B., Lopez Meneses, D. H., Matsler, A. M., Moglia, M., Morató, J., … Zhou, W. (2025). Global synthesis and regional insights for mainstreaming urban nature-based solutions. Proceedings of the National Academy of Sciences, 122(29), e2315910121. 10.1073/pnas.2315910121

McRae BH, et al. (2016) Conserving nature’s stage: mapping omnidirectional connectivity for resilient terrestrial landscapes in the pacific northwest (The Nature Conservancy, Portland, Oregon).

Metzger, J. P., Villarreal-Rosas, J., Suárez-Castro, A. F., López-Cubillos, S., González-Chaves, A., Runting, R. K.,… Rhodes, J. R. (2021). Considering landscape-level processes in ecosystem service assessments. Science of the Total Environment, 796, 149028.

Mitchell, M. G., Bennett, E. M., & Gonzalez, A. (2013). Linking landscape connectivity and ecosystem service provision: current knowledge and research gaps. Ecosystems, 16(5), 894–908.

Mitchell, M. G., Bennett, E. M., Gonzalez, A., Lechowicz, M. J., Rhemtulla, J. M., Cardille, J. A., … & Dancose, K. (2015). The Montérégie Connection: linking landscapes, biodiversity, and ecosystem services to improve decision making. Ecology and Society, 20(4). Natural Capital Project (2023) InVEST 0.0. (Stanford University, et al.). Available at: https://naturalcapitalproject.stanford.edu/software/invest.

Nesshöver, C., Assmuth, T., Irvine, K. N., Rusch, G. M., Waylen, K. A., Delbaere, B., … & Wittmer, H. (2017). The science, policy and practice of nature-based solutions: An interdisciplinary perspective. Science of the total environment, 579, 1215–1227.

Neyret, M., Richards, D., Prima, M. C., Etherington, T. R., & Lavorel, S. (2025). One cannot have it all: Trading-off ecosystem services and biodiversity bundles in landscape connectivity restoration. Biological Conservation, 302, 110946.

Norström, A. V., Cvitanovic, C., Löf, M. F., West, S., Wyborn, C., Balvanera, P., … Österblom, H. (2020). Principles for knowledge co-production in sustainability research. Nature Sustainability, 3(3), 182–190. 10.1038/s41893-019-0448-2

Obura, D. (2023). The Kunming-Montreal global biodiversity framework: business as usual or a turning point?. One Earth, 6(2), 77–80.

Palomo, I., Locatelli, B., Otero, I., Colloff, M., Crouzat, E., Cuni-Sanchez, A.,… Lavorel, S. (2021). Assessing nature-based solutions for transformative change. One Earth, 4(5), 730–741.

Pascual, U., Balvanera, P., Díaz, S., Pataki, G., Roth, E., Stenseke, M.,… Wittmer, H. (2017). Valuing nature’s contributions to people: The IPBES approach. Current Opinion in Environmental Sustainability, 26, 7–16.

Pe’er, G., Bonn, A., Bruelheide, H., Dieker, P., Eisenhauer, N., Feindt, P. H., … & Lakner, S. (2020). Action needed for the EU Common Agricultural Policy to address sustainability challenges. People and nature, 2(2), 305–316.

Perissi, I. (2025). Assessing the EU27 Potential to Meet the Nature Restoration Law Targets. Environmental Management, 75(4), 711–729.

Pineda-Zapata, S., Morán-Ordoñez, A., Mola-Yudego, B., & Duflot, R. (2026). Strategic placement of plantations enhances forest connectivity for birds in agricultural landscapes. Landscape Ecology, 41(3), 54.

Plieninger, T., Torralba, M., Hartel, T., & Fagerholm, N. (2019). Perceived ecosystem services synergies, trade-offs, and bundles in European high nature value farming landscapes. Landscape Ecology, 34(7), 1565–1581.

Prima, M. C., Renaud, J., Witté, I., Suarez, L., Rouveyrol, P., Fernando, M., … & Thuiller, W. (2024). A comprehensive framework to assess multi-species landscape connectivity. Methods in Ecology and Evolution, 15(12), 2385–2399..

Raudsepp-Hearne, C., Peterson, G. D., & Bennett, E. M. (2010). Ecosystem service bundles for analyzing tradeoffs in diverse landscapes. Proceedings of the National Academy of Sciences, 107(11), 5242–5247.

Raymond, C. M., Frantzeskaki, N., Kabisch, N., Berry, P., Breil, M., Nita, M. R., Geneletti, D., & Calfapietra, C. (2017). A framework for assessing and implementing the co-benefits of nature-based solutions in urban areas. Environmental Science & Policy, 77, 15–24. 10.1016/j.envsci.2017.07.008

Rois-Díaz, M., Lovric, N., Lovric, M., Ferreiro-Domínguez, N., Mosquera-Losada, M. R., Den Herder, M., … & Burgess, P. (2018). Farmers’ reasoning behind the uptake of agroforestry practices: evidence from multiple case-studies across Europe. Agroforestry Systems, 92(4), 811–828.

Saidi, N., & Spray, C. (2018). Ecosystem services bundles: challenges and opportunities for implementation and further research. Environmental Research Letters, 13(11), 113001.

Schulp, C. J. E., Lautenbach, S., & Verburg, P. H. (2014). Quantifying and mapping ecosystem services: Demand and supply of pollination in the European Union. Ecological Indicators, 36, 131–141.

Seddon, N., Chausson, A., Berry, P., Girardin, C. A. J., Smith, A., & Turner, B. (2020). Understanding the value and limits of nature-based solutions to climate change and other global challenges. Philosophical Transactions of the Royal Society B: Biological Sciences, 375(1794), 20190120.

Seddon, N., Smith, A., Smith, P., Key, I., Chausson, A., Girardin, C., … & Turner, B. (2021). Getting the message right on nature-based solutions to climate change. Global change biology, 27(8), 1518–1546.

Sharp, R., Douglass, J., Wolny, S., Arkema, K., Bernhardt, J., Bierbower, W.,… Glowinski, K. (2020). InVEST 3.8.7. User’s Guide. The Natural Capital Project.

Tobin, D., Pfaff, A., Vincent, J. R., Vanamamalai, A., & Karanth, K. K. (2026). Guiding private afforestation to raise public goods provision: Understanding farmers’ multi-dimensional preferences for trees in India. Ecological Economics, 247, 109023.

Vannier, C., Lasseur, R., Crouzat, E., Byczek, C., Lafond, V., Cordonnier, T., … & Lavorel, S. (2019). Mapping ecosystem services bundles in a heterogeneous mountain region. Ecosystems and People, 15(1), 74–88.

Watson, J. E. M., Dudley, N., Segan, D. B., & Hockings, M. (2014). The performance and potential of protected areas. Nature, 516, 67–73.

Welden, E. A., Chausson, A., & Melanidis, M. S. (2021). Leveraging nature-based solutions for transformation: Reconnecting people and nature. People and Nature, 3(5), 966–977.

Zu Ermgassen, S. O., Hawkins, I., Lundhede, T., Liu, Q., Thorsen, B. J., & Bull, J. W. (2025). The current state, opportunities and challenges for upscaling private investment in biodiversity in Europe. Nature Ecology & Evolution, 1–10.

