## Appendix_Gonzalez_Garcia_et_al_2026_scenarios for "Accounting for social feasibility in landscape-scale scenarios for restoring nature beyond protected areas": Gonzalez_Garcia_et_al_Appendix_Reconnect_scenarios_01_09_2026.pdf

### A. Study Area

Our study was conducted in the Grenoble region of the French Alps, a well-studied landscape of approximately 4,500 km<sup>2</sup> characterized by a strong socio-ecological heterogeneity.

The region's land use is structured along a steep elevation gradient (Fig. S1). The lowlands are dominated by two main agricultural plains, where intensive cropping (e.g., maize and walnuts) and grasslands are interspersed with a dense urban fabric, including the Grenoble metropolitan area (Fig. S1A). These human-dominated valleys contrast sharply with the mountain areas, which are primarily covered by extensive forests, sub-alpine grasslands, and alpine ecosystems. While these mountain landscapes are central to the region's recreational activities, they also support traditional pastoralism and forestry.

This heterogeneous matrix hosts a diverse network of protected areas. For this analysis, we selected 16 sites that are representative of the region's conservation efforts, ranging from large regional parks in mountain areas to smaller reserves in the agricultural lowlands (Fig. S1B). The region's pronounced topography, with elevations spanning from lowland valleys to high peaks, is a key driver of this landscape structure (Fig. S1C). This combination of intensive agriculture, a dynamic urban population, and high-value natural areas makes the region an ideal natural laboratory for studying the complex interfaces between conservation and production landscapes.

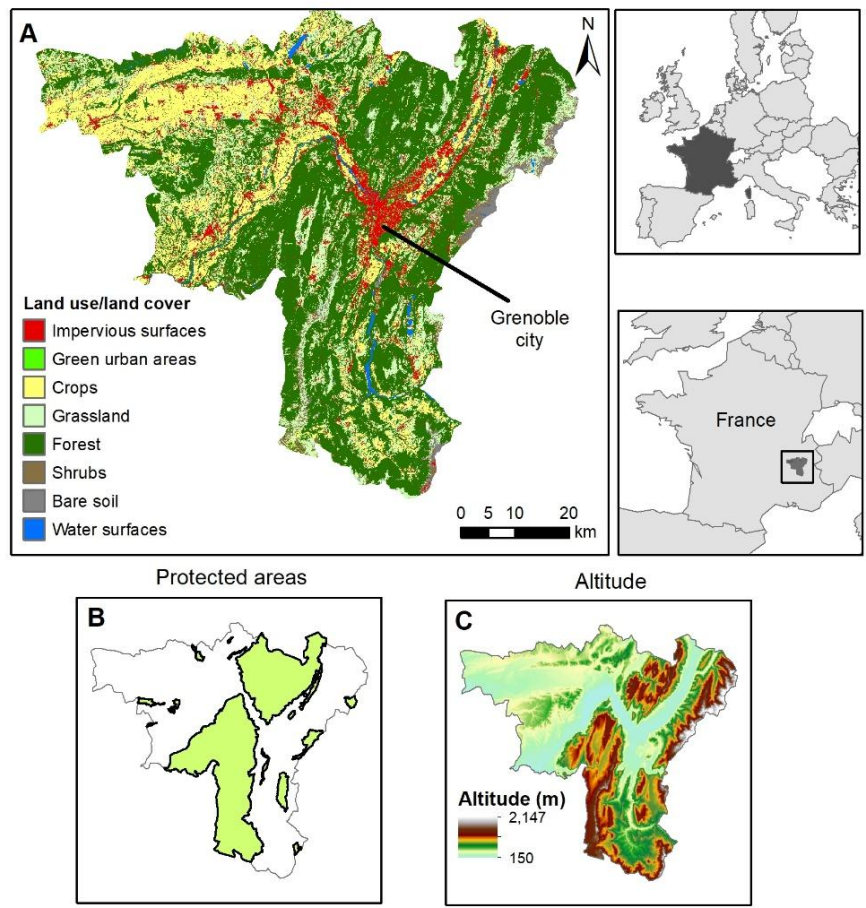

Figure S 1. Description of the study area. The panels illustrate the geographical and environmental context of the Grenoble region. (A) A detailed land use/land cover map showing the heterogeneous mosaic of urban centres (e.g., Grenoble city), agricultural valleys, and forested mountain slopes. (B) The location of the 16 protected areas selected for the analysis. (C) An altitude map highlighting the region's pronounced topography, from the lowland valleys to the high-elevation mountain ranges.

### **B. Baseline ecosystem service modelling**

This section details the methodologies used to model the 12 ecosystem services for the baseline assessment, as summarized in the main text. The selection of these ecosystem services was guided by their high relevance to the socio-ecological dynamics of the French Alps, as identified in previous regional stakeholder workshops (Vannier et al., 2019). All final raster maps were produced at a 5-meter resolution using a regional Land Use/Land Cover map (Marsoner et al., 2023) as a common base.

#### **a. Modelling Approaches**

Six ecosystem services were modelled using established biophysical approaches, primarily from the InVEST 4.0 software suite (Sharp et al., 2020), which combines land use/land cover maps with specific biophysical parameters to quantify ecosystem functions. Key input data included the 5m land use/land cover map, a 5m Digital Elevation Model (DEM), and regional soil and climate data. Each model's specific parameterization is detailed in the individual ecosystem service descriptions below.

For the six ecosystem services dependent on mobile organisms, we developed and applied a comprehensive framework to translate species' potential presence into a spatially explicit map of ecosystem service provision. This framework integrates species distribution modelling, fine-scale habitat information, and landscape connectivity theory.

##### **Step 1: European-scale habitat suitability modelling (1 km resolution)**

The foundation of the framework is a set of habitat suitability models for 224 vertebrate species known to be present in the region. These models were developed for the entire European continent at a 1 km resolution using an ensemble species distribution modelling approach (Si-Moussi & Thuiller, 2024). The ensemble model combines predictions from multiple machine-learning algorithms (Random Forest, XGBoost, Neural Networks) trained on species occurrence data (GBIF) and a wide range of environmental predictors (e.g., climate, topography, soil properties). This approach yields robust, consensus-based predictions of suitable environmental conditions for each species at a continental scale.

##### **Step 2: Downscaling to local landscape (5 m resolution)**

To bridge the scale gap from the coarse 1 km predictions to our fine-scale 5m landscape, we implemented a downscaling procedure. First, we established a correspondence between each species' documented habitat preferences (O'Connor et al., 2024), originally defined for GlobCover classes, and the classes of our regional EUSALP land use/land cover map using a crosswalk table (Table S1). This allowed us to create a binary "local habitat mask" at 5m resolution for each species, identifying all pixels corresponding to a suitable land use/land cover type. We then resampled the 1 km European suitability map to 5m resolution and multiplied it by this binary mask. The resulting 5m habitat suitability map combines the broad environmental suitability from the species distribution modelling with the fine-grained constraints imposed by the local landscape structure.

*Table S 1. Crosswalk references between EUSALP land use cover used for the study and GlobCover.*

| **Code** | **EUSALP** | **GlobCover** |
| --- | --- | --- |
| **11000** | Artificial surfaces and constructions | 190 |
| **11100** | Dense settlement area | 190 |
| **11200** | Low density settlement area | 190 |
| **11300** | Builtup area | 190 |
| **11400** | Open settlement area | 190 |
| **12100** | Industrial and commercial zones | 190 |
| **12210** | Roads motorways and trunks | 190 |
| **12220** | Roads primary and secondary | 190 |
| **12221** | Roads tertiary and others | 190 |
| **12230** | Railways train tracks | 190 |
| **12240** | Unpaved Roads and Tracks | 190 |
| **14100** | Green urban areas | 190 |
| **21000** | Cultivated areas - Arable Land - Annual Crops | 10,11,13,14,15 |
| **21211** | Common wheat | 10,15 |
| **21212** | Durum wheat | 10,15 |
| **21213** | Barley | 10,15 |
| **21214** | Rye | 10,15 |
| **21215** | Oats | 10,15 |
| **21216** | Maize | 10,13 |
| **21217** | Rice | 10,11 |
| **21218** | Triticale | 10,15 |
| **21219** | Other cereals | 10,15 |
| **21221** | Potatoes | 10,14 |
| **21222** | Sugar beet | 10,14 |
| **21223** | Other root crops | 10,14 |
| **21230** | Other non permanent industrial crops | 10,14,15 |
| **21231** | Sunflower | 10,14 |
| **21232** | Rape and turnip rape | 10,14 |
| **21233** | Soya | 10,15 |
| **21240** | Dry pulses | 10,14,15 |
| **21250** | Fodder crops (cereals and leguminous) | 10,14,15 |
| **21290** | Bare arable land | 10,14,15 |
| **22000** | Permanent Crops | 10,16 |
| **22100** | Vinyard | 10,16 |
| **22200** | Orchard | 10,16 |
| **23100** | Managed grassland - Pastures | 20, 21, 30 |
| **23200** | Seminatural grassland - Meadows | 140,141,144,151,110,120 |
| **31100** | Broadleaf tree cover | 40, 41, 50, 60, 100, 101, 32 |
| **31102** | Broadleaf tree cover 30-60% | 40, 41, 50, 60, 100, 101, 32 |
| **31103** | Broadleaf tree cover 60-100% | 40, 41, 50, 60, 100, 101, 32 |
| **31200** | Coniferous tree cover | 70, 90, 91, 92, 100, 101, 32 |
| **31202** | Coniferous tree cover 30-60% | 70, 90, 91, 92, 100, 101, 32 |
| **31203** | Coniferous tree cover 60-100% | 70, 90, 91, 92, 100, 101, 32 |
| **31300** | Mixed tree cover | 40, 41, 50, 60, 100, 101, 32, 70, 90, 91, 92, 100, 101, 32 |
| **31400** | Tree cover in agricultural context | 32 |
| **31450** | Tree cover in urban context | 190 |
| **31500** | Green linear elements - linear woody features | 130,131,132,133,134,136 |
| **31600** | Patchy woody features | 130,131,132,133,134,136 |
| **31610** | Additional woody features | 130,131,132,133,134,136 |
| **32000** | Scrub and shrubland | 130,131,132,133,134,136,152, 110,120 |
| **32100** | Alpine and sub-alpine natural grassland | 140,141,144,151,110,120 |
| **32200** | Moors and Heathland - other scrubland | 130,131,132,133,134,136,152, 110,120 |
| **32300** | Sclerophyllous vegetation | 130,131,132,133,134,136,152, 110,120 |
| **33100** | Beaches, dunes, sands | 200, 202,150,151,152 |
| **33200** | Bare rocks and rock debris | 200, 202,150,151,152 |
| **33300** | Sparsely vegetated land | 150,151,152 |
| **33500** | Permanent snow covered surfaces | 220 |
| **41000** | Wetland (permanent wet areas) - inland marshes | 180,185 |
| **41200** | Peatbogs | 180,185 |
| **42100** | Coastal salt marshes | 180,185 |
| **42200** | Intertidal flats | 180,185 |
| **51000** | Water bodies | 210 |
| **51100** | Rivernetwork | 210 |
| **51200** | Riverbed > 10m width | 210 |
| **52100** | Lagoons and Estuaries | 210 |

#### Step 3: Functional landscape connectivity modelling (Omniscape)

To explicitly account for species movement, we modelled functional landscape connectivity using the Omniscape algorithm (McRae et al., 2016). Based on circuit theory, Omniscape calculates "current flow" across the landscape, which represents the probability of landscape use by a moving organism. The algorithm requires two main inputs derived from the 5m habitat suitability map:

• Source map: Pixels with a habitat suitability probability exceeding a species-specific optimal threshold were defined as "sources" from which movement originates.
• Resistance map: The habitat suitability map was converted into a resistance surface using a non-linear transformation, where higher suitability corresponds to lower resistance to movement.

Omniscape then calculates cumulative current flow using a moving window radius defined by each species' mean dispersal distance. The output is a "cumulative current" map, representing a connectivity-corrected probability of landscape use. For species with very broad dispersal capacities, landscape connectivity was assumed to be complete at the scale of our study area, and their downscaled habitat suitability map was used directly.

##### Step 4: Aggregation and ecosystem service quantification

Finally, the connectivity-corrected maps of all species contributing to a given ecosystem service (e.g., all pest predators for "Biological Control") were averaged with equal weighting to create a single, composite ecosystem service map. This approach assumes functional redundancy, where the presence of a diverse assemblage is key to ecosystem service provision. The resulting map represents the overall potential for a given function across the landscape, ready for the final integration of human demand criteria as described for each ecosystem service below.

##### b. Biophysical ecosystem service models

###### i. Food Production

We calculated food production as the energy equivalent (GJ/ha) of the average annual crop yield, based on regional statistics (Lasseur et al., 2018). The model uses a table of energy values for each crop type present in the land use/land cover map (Table S2). As food produced in the region is primarily traded on national and international markets, demand was considered delocalized, and thus only supply was mapped.

*Table S 2. Yields used for the food production model. "Yield (100 kg/ha)" represents the direct value from public authorities for each crop type. "Tn/ha" is the conversion to tons per hectare. "Energy (MJ/kg)" represents the transformation of yields into energy, and "GJ/5m" is the final conversion based on a 5-meter pixel size.*

| CODE | DESC | Yield<br>100kg/ha | Tn/ha | Energy<br>Mj/kg | GJ/5m |
| --- | --- | --- | --- | --- | --- |
| **21000** | Cultivated areas - Arable land - Annual crops | 58.7 | 5.9 | 11.3 | 3.3 |
| **21211** | Common wheat | 54.0 | 5.4 | 11.4 | 3.1 |
| **21212** | Durum wheat | 53.0 | 5.3 | 11.4 | 3.0 |
| **21213** | Barley | 68.0 | 6.8 | 11.5 | 3.9 |
| **21214** | Rye | 44.0 | 4.4 | 12.1 | 2.7 |
| **21215** | Oats | 43.0 | 4.3 | 10.2 | 2.2 |
| **21216** | Maize | 98.0 | 9.8 | 11.0 | 5.4 |
| **21218** | Triticale | 51.0 | 5.1 | 11.4 | 2.9 |
| **21221** | Potatoes | 422.0 | 42.2 | 2.7 | 5.8 |
| **21222** | Sugar beet | 428.0 | 42.8 | 2.4 | 5.1 |
| **21230** | Other non-permanent industrial crops | 52.7 | 5.3 | 37.1 | 9.8 |
| **21231** | Sunflower | 25.0 | 2.5 | 15.3 | 1.9 |
| **21232** | Rape and turnip rape | 32.0 | 3.2 | 15.3 | 2.4 |
| **21233** | Soya | 24.0 | 2.4 | 10.2 | 1.2 |
| **21240** | Dry pulses | 28.0 | 2.8 | 14.0 | 2.0 |
| **21250** | Fodder crops (cereals and leguminous) | 138.0 | 13.8 | 2.0 | 1.4 |
| **21290** | Bare arable land | 0.0 | 0.0 | 0.0 | 0.0 |
| **22000** | Permanent crops | 12.0 | 1.2 | 15.6 | 0.9 |
| **22100** | Vinyard | 83.0 | 8.3 | 2.9 | 1.2 |
| **22200** | Orchard | 286.6 | 28.7 | 6.9 | 2.8 |
| **23100** | Managed grassland - Pastures | 83.0 | 8.3 | 3.8 | 1.6 |
| **23200** | Seminal natural grassland - Meadows | 58.0 | 5.8 | 3.8 | 1.1 |
| **32100** | Alpine and sub-alpine natural grassland | 146.0 | 14.6 | 3.8 | 2.7 |

### ii. Carbon Storage

We used the InVEST Carbon Storage and Sequestration model to estimate the total carbon stored in four pools: aboveground biomass, belowground biomass, soil organic matter, and dead organic matter. The model was parameterized using a table of carbon stock values for each land use/land cover class, compiled from regional literature (Table S3). As a globally relevant ecosystem service, only supply was mapped.

*Table S 3. Carbon pools in CO<sub>2</sub> eq. used for the InVEST model. C<sub>soil</sub> is the organic carbon in soil, C<sub>Above</sub> is the carbon in aboveground biomass, C<sub>Below</sub> is the carbon in belowground biomass, C<sub>litter</sub> is the carbon in dead matter.*

| CODE | Description | C<sub>soil</sub> | C<sub>Above</sub> | C<sub>Below</sub> | C<sub>litter</sub> | References |
| --- | --- | --- | --- | --- | --- | --- |
| 11000 | Artificial surfaces and constructions | 0.18 | 0.10 | 0 | 0 | González-García et al. (2020) |
| 11100 | Dense settlement area | 0.18 | 0.10 | 0 | 0 |  |
| 11200 | Low density settlement area | 64.04 | 16.86 | 8.57 | 0 | Dorendorf et al. (2015) |
| 11300 | Builtup area | 0.18 | 0.10 | 0 | 0 | González-García et al. (2020) |
| 11400 | Open settlement area | 64.04 | 16.86 | 8.57 | 0.00 | Dorendorf et al. (2015) |
| 12100 | Industrial and commercial zones | 12.34 | 14.44 | 0 | 0 |  |
| 12210 | Roads motorways and trunks | 0.18 | 0.10 | 0 | 0 | González-García et al. (2020) |
| 12220 | Road networks | 0.18 | 0.10 | 0 | 0 |  |
| 12221 | Roads tertiary and others | 0.18 | 0.10 | 0 | 0 |  |
| 12230 | Railways train tracks | 0.18 | 0.10 | 0 | 0 |  |
| 12240 | Unpaved roads and tracks | 0.18 | 0.10 | 0 | 0 |  |
| 14100 | Green urban areas | 258.82 | 135.8 | 50.62 | 0 | Dorendorf et al. (2015) |
| 21000 | Cultivated areas - Arable land - Annual crops | 81.68 | 5.07 | 0 | 0 | Dorendorf et al. (2015) |
| 21211 | Common wheat | 81.68 | 5.07 | 0 | 0 |  |
| 21212 | Durum wheat | 81.68 | 5.07 | 0 | 0 |  |
| 21213 | Barley | 81.68 | 5.07 | 0 | 0 |  |
| 21214 | Rye | 81.68 | 5.07 | 0 | 0 |  |
| 21215 | Oats | 81.68 | 5.07 | 0 | 0 |  |
| 21216 | Maize | 81.68 | 5.07 | 0 | 0 |  |
| 21218 | Triticale | 81.68 | 5.07 | 0 | 0 |  |
| 21221 | Potatoes | 81.68 | 5.07 | 0 | 0 |  |
| 21222 | Sugar beet | 81.68 | 5.07 | 0 | 0 |  |
| 21230 | Other non-permanent industrial crops | 81.68 | 5.07 | 0 | 0 |  |
| 21231 | Sunflower | 81.68 | 5.07 | 0 | 0 |  |
| 21232 | Rape and turnip rape | 81.68 | 5.07 | 0 | 0 |  |
| 21233 | Soya | 81.68 | 5.07 | 0 | 0 |  |
| 21240 | Dry pulses | 81.68 | 5.07 | 0 | 0 |  |
| 21250 | Fodder crops (cereals and leguminous) | 81.68 | 5.07 | 0 | 0 |  |
| 21290 | Bare arable land | 81.68 | 0 | 0 | 0 |  |
| **22000** | Permanent crops | 151.48 | 52.923 | 20.22 | 0 | Cardinael et al. (2017) |
| **22100** | Vineyard | 153.63 | 6.2 | 8.57 | 0 | Chiti, et al. (2012) |
| **22200** | Orchard | 161.7 | 0 | 0 | 0 | Chiti, et al. (2012) |
| **23100** | Managed grassland - Pastures | 277.97 | 7.13 | 13.2 | 0 | Bahn et al. (2008); Guidi et al. 2014 (managed grassland) |
| **23200** | Seminatural grassland - Meadows | 277.97 | 7.13 | 13.2 | 0 |  |
| **31100** | Broadleaf tree cover | 296.75 | 176.41 | 67.42 | 18.11 | INFC 2005, Tonolli & Salvagni, (2007) |
| **31102** | Broadleaf tree cover 30-60% | 296.75 | 52.92 | 20.22 | 5.43 |  |
| **31103** | Broadleaf tree cover 60-100% | 296.75 | 105.84 | 40.45 | 10.86 |  |
| **31200** | Coniferous tree cover | 258.88 | 267.96 | 76.52 | 30.09 |  |
| **31202** | Coniferous tree cover 30-60% | 258.88 | 80.38 | 22.95 | 9.02 |  |
| **31203** | Coniferous tree cover 60-100% | 258.88 | 160.77 | 45.91 | 18.05 |  |
| **31300** | Mixed tree cover | 277.81 | 222.18 | 71.97 | 24.1 | Cardinael et al. (2017) |
| **31400** | Tree cover in agricultural context | 151.48 | 52.92 | 20.22 | 0 |  |
| **31450** | Tree cover in urban context | 0.18 | 135.8 | 50.62 | 0 | Dorendorf et al. (2015) |
| **31500** | Green linear elements - linear woody features | 84.19 | 6.20 | 8.57 | 4.90 | Sørensen et al., (2018) |
| **31600** | Patchy woody features | 84.19 | 6.20 | 8.57 | 4.90 |  |
| **31610** | Additional woody features | 84.19 | 6.20 | 8.57 | 4.90 |  |
| **32000** | Scrub and shrubland | 84.19 | 6.20 | 8.57 | 4.90 |  |
| **32100** | Alpine and sub-alpine natural grassland | 277.97 | 7.13 | 13.2 | 0 | Fonseca et al. (2012) |
| **32200** | Moors and heathland - other scrubland | 84.19 | 6.20 | 8.57 | 4.90 |  |
| **32300** | Sclerophyllous vegetation | 97.8 | 5.69 | 7.5 | 1.53 |  |
| **33100** | Beaches, dunes, sands | 18.51 | 0 | 0 | 0 |  |
| **33200** | Bare rocks and rock debris | 0 | 0 | 0 | 0 | González-García et al. (2020) |
| **33300** | Sparsely vegetated land | 12.34 | 6.2 | 8.57 | 4.9 |  |
| **33500** | Permanent snow covered surfaces | 0 | 0 | 0 | 0 | IPCC |
| **41000** | Wetland (permanent wet areas) - inland marshes | 539 | 43 | 0 | 0 |  |
| **51000** | Water bodies | 320.1 | 0.44 | 0.25 | 1.83 | González-García et al. (2020) |
| **51100** | Rivernetwork | 320.1 | 0.44 | 0.25 | 1.83 |  |

#### iii. Outdoor Recreation

We assessed outdoor recreation potential using a regional model adapted from Byczek et al. (2018), which is based on the Recreation Opportunity Spectrum framework. The model calculates a Recreation Potential Index by combining multiple spatial indicators that represent landscape attractiveness, avoidance of disturbances, and accessibility.

The Landscape Attractiveness component was calculated based on three main factors:

• Proximity to natural and aquatic features: Pixels were assigned higher values based on their proximity to natural land covers (e.g., forests, alpine grasslands) and aquatic elements

(rivers, lakes). We used a 500m buffer to model the positive influence of nearby water bodies.

• Degree of nature protection: We incorporated a pre-calculated layer from Byczek et al. (2018) that scores the level of nature protection, assigning higher values to areas with stricter conservation statuses (e.g., national parks over regional parks), as this is often correlated with higher perceived naturalness.
• Scenic value: We included a scenic value index, also from Byczek et al. (2018), which was derived from a viewshed analysis combined with measures of landscape heterogeneity. This factor assigns higher values to pixels from which a greater diversity of landscape elements is visible, serving as a proxy for panoramic quality.

The Avoidance of Disturbances component integrates two factors that detract from the recreational experience:

• Road and noise avoidance: We used a layer that assigns lower values to areas close to major roads, railways, and airports, based on buffer distances reflecting the impact of noise pollution.
• Avoidance of artificialized areas: Pixels corresponding to urban, industrial, or intensively managed agricultural land received a value of zero, effectively masking out areas with low recreational appeal.

Finally, Accessibility was the key component for integrating human demand. Instead of using simple distance metrics, we incorporated a comprehensive accessibility layer from Byczek et al. (2018) derived from a massive dataset of crowd-sourced GPS tracks from platforms like skitour.fr and vttour.fr. This layer represents the observed density of recreational use across the landscape for a wide range of activities (hiking, biking, skiing, etc.). By using this data, the final model is weighted not just by theoretical potential but by demonstrated visitor presence and preferences, providing a robust, demand-integrated measure of the outdoor recreation ecosystem service (Fig. S2).

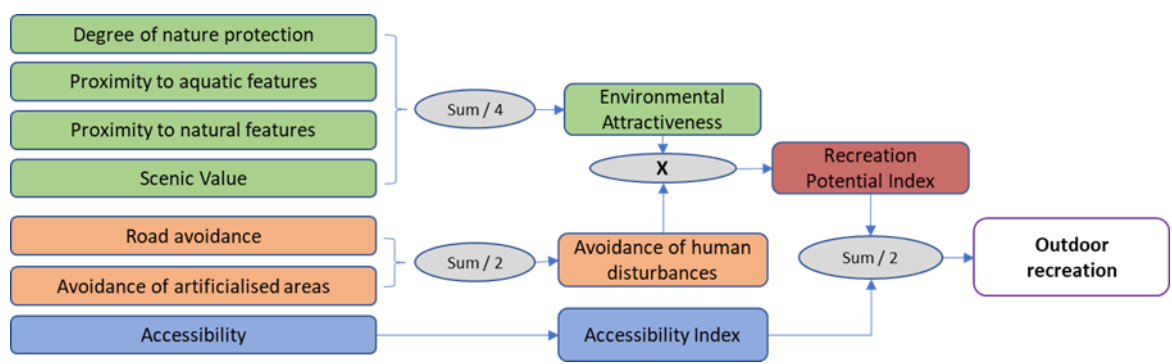

Figure S 2. Conceptual design of the outdoor recreation model

##### iv. Flood Regulation

We used the InVEST Urban Flood Regulation model to quantify the capacity of landscapes to mitigate flood risk by retaining runoff during storm events. The model calculates the amount of runoff generated per pixel, and the ecosystem service is quantified as the volume of water retained by the ecosystem.

The model's calculations are based on the widely used curve number method, which estimates direct runoff from a given rainfall event. The primary inputs for the model were:

• Rainfall depth: We set a reference rainfall depth of 100 mm for the storm event (Goswami et al., 2006). This value was chosen to represent a high-intensity, low-frequency storm event (i.e., a significant flood risk scenario), aligning with common hydrological modelling practices for risk assessment in European mountainous regions. The use of a risk-based scenario implicitly integrates human demand, as the ecosystem service is most relevant under conditions where a societal need for flood mitigation is highest.
• Soil hydrologic group: We used a 250m resolution map of soil hydrologic groups for Europe (Ross et al., 2018). This dataset classifies soils into four groups (A, B, C, D) based on their infiltration capacity, from high (A) to low (D).
• Land Use/Land Cover: We used our 5m resolution regional land use/land cover map.

The core of the model is the biophysical table, where each land use/land cover class is assigned a specific curve number for each soil hydrologic group (Table S4). These curve number values, which range from 0 to 100, represent the runoff potential of a given land use/land cover-soil combination. The values were derived from the standard tables provided by the USDA Natural Resources Conservation Service (NRCS), adapted to our specific LULC classes following the equivalencies established by Cronshey (1986) and recent European studies (Jaafar et al., 2019; Tedela et al., 2012).

The model computes the runoff retention for each pixel by subtracting the generated runoff from the initial rainfall depth. The final output is a map of runoff retention volume per pixel (in m<sup>3</sup>), where higher values indicate a greater capacity of the landscape to mitigate flooding during a major storm event.

*Table S 4. Biophysical table associated with the flood regulation model. CN values represent the curve numbers for each soil hydrologic group (A, B, C, or D). The "Ref" field indicates the equivalencies used from the main sources related to land use/land cover categories.*

| CODE | Description | CN_a | CN_b | CN_c | CN_d | Reference | Ref |
| --- | --- | --- | --- | --- | --- | --- | --- |
| 11000 | Artificial surfaces and constructions | 98 | 98 | 98 | 98 | Cronshey (1986) | Impervious areas |
| 11100 | Dense settlement area | 98 | 98 | 98 | 98 |  |  |
| 11200 | Low density settlement area | 61 | 75 | 83 | 87 |  | 1/4 acre (38% imp.) |
| 11300 | Builtup area | 98 | 98 | 98 | 98 |  | Impervious areas |
| 11400 | Open settlement area | 61 | 75 | 83 | 87 |  | 1/4 acre (38% imp.) |
| 12100 | Industrial and commercial zones | 98 | 98 | 98 | 98 |  | Impervious areas |
| 12210 | Roads motroways and trunks | 98 | 98 | 98 | 98 |  |  |
| 12220 | Road networks | 98 | 98 | 98 | 98 |  |  |
| 12221 | Roads tertiary and others | 98 | 98 | 98 | 98 |  |  |
| 12230 | Railways train tracks | 98 | 98 | 98 | 98 |  |  |
| 12240 | Unpaved roads and tracks | 77 | 86 | 91 | 94 |  | Bare soil |
| 14100 | Green urban areas | 39 | 61 | 74 | 80 |  | Open space (Good condition) |
| **21000** | Cultivated areas -<br>Arable land -<br>Annual crops | 60 | 72 | 80 | 84 |  | Small grain SR+CR good |
| **21211** | Common wheat | 60 | 72 | 80 | 84 |  |  |
| **21212** | Durum wheat | 60 | 72 | 80 | 84 |  |  |
| **21213** | Barley | 60 | 72 | 80 | 84 |  |  |
| **21214** | Rye | 60 | 72 | 80 | 84 |  |  |
| **21215** | Oats | 60 | 72 | 80 | 84 |  |  |
| **21216** | Maize | 65 | 75 | 82 | 86 |  | Row crops contoured |
| **21218** | Triticale | 60 | 72 | 80 | 84 |  | Small grain SR+CR good |
| **21221** | Potatoes | 32 | 58 | 72 | 79 |  | Orchard |
| **21222** | Sugar beet | 32 | 58 | 72 | 79 |  |  |
| **21230** | Other non<br>permanent<br>industrial crops | 60 | 72 | 80 | 84 |  | Small grain SR+CR good |
| **21231** | Sunflower | 64 | 75 | 82 | 85 | Cronshey<br>(1986) | Row crops SR+CR |
| **21232** | Rape and turnip<br>rape | 64 | 75 | 82 | 85 |  | Close-seeded or broadcast<br>legumes or rotation<br>meadow |
| **21233** | Soya | 58 | 72 | 81 | 85 |  |  |
| **21240** | Dry pulses | 58 | 72 | 81 | 85 |  |  |
| **21250** | Fodder crops<br>(cereals and<br>leguminous) | 58 | 72 | 81 | 85 |  |  |
| **21290** | Bare arable land | 77 | 86 | 91 | 94 |  | Bare arable land |
| **22000** | Permanent crops | 43 | 65 | 76 | 82 |  | Agro-forestry areas |
| **22100** | Vinyard | 67 | 78 | 85 | 89 |  | Straight row |
| **22200** | Orchard | 32 | 58 | 72 | 79 |  | Orchard |
| **23100** | Managed<br>grassland -<br>Pastures | 8 | 79 | 86 | 89 |  | Pasture |
| **23200** | Seminatural<br>grassland -<br>Meadows | 30 | 58 | 71 | 78 |  | Meadow |
| **31100** | Broadleaf tree<br>cover | 30 | 55 | 70 | 77 |  | Good condition |
| **31102** | Broadleaf tree<br>cover 30-60% | 36 | 60 | 73 | 79 |  | Fair condition |
| **31103** | Broadleaf tree<br>cover 60-100% | 30 | 55 | 70 | 77 |  | Good condition |
| **31200** | Coniferous tree<br>cover | 33 | 58 | 72 | 78 | Jaafar et al.<br>(2019); Tedela<br>et al. (2012) | Good condition |
| **31202** | Coniferous tree<br>cover 30-60% | 40 | 63 | 75 | 80 |  | Fair condition |
| **31203** | Coniferous tree<br>cover 60-100% | 33 | 58 | 72 | 78 |  | Good condition |
| **31300** | Mixed tree cover | 31.5 | 56.5 | 71 | 77.5 |  | Average broadleaf and<br>conifer |
| **31400** | Tree cover in<br>agricultural<br>context | 43 | 65 | 76 | 82 | Cronshey<br>(1986) | Woods—grass<br>combination (orchard or<br>tree farm).D (Fair) |
| **31450** | Tree cover in<br>urban context | 39 | 61 | 74 | 80 |  | Open space (Good<br>condition) |
| **31500** | Green linear<br>elements - linear<br>woody features | 43 | 65 | 76 | 82 |  | Woods—grass<br>combination (orchard or<br>tree farm).D |
| **31600** | Patchy woody features | 43 | 65 | 76 | 82 |  |  |
| **31610** | Additional woody features | 43 | 65 | 76 | 82 |  |  |
| **32000** | Scrub and shrubland | 43 | 65 | 76 | 82 |  |  |
| **32100** | Alpine and sub-alpine natural grassland | 39 | 61 | 74 | 80 |  | Brush—brush-weed-grass mixture with brush the major element.B (Fair) |
| **32200** | Moors and heathland - other scrubland | 35 | 56 | 70 | 77 |  | Brush—brush-weed-grass mixture with brush the major element.B |
| **32300** | Sclerophyllous vegetation | 0 | 62 | 74 | 85 |  | Herbaceosu for arid rangelands |
| **33100** | Beaches, dunes, sands | 63 | 77 | 85 | 88 |  | Natural desert landscaping (pervious areas only) |
| **33200** | Bare rocks and rock debris | 63 | 77 | 85 | 88 |  |  |
| **33300** | Sparsely vegetated land | 68 | 79 | 86 | 89 |  | Poor condition (grass cover < 50%) |
| **33500** | Permanent snow covered surfaces | 99 | 99 | 99 | 99 |  |  |
| **41000** | Wetland (permanent wet areas) - inland marshes | 49 | 69 | 79 | 84 | Chen et al. (2014) | Wet tussock grassland with herbs, sedges or rushes, herblands or ferns |
| **51000** | Water bodies | 99 | 99 | 99 | 99 |  |  |
| **51100** | Rivernetwork | 99 | 99 | 99 | 99 |  |  |
| **51200** | Riverbed > 10m width | 99 | 99 | 99 | 99 |  |  |

##### v. Erosion Control

We used the InVEST Sediment Delivery Ratio model to quantify the capacity of the landscape to retain soil, a ecosystem service critical for preventing land degradation and protecting water quality. The model estimates the amount of annual soil loss per pixel and the proportion of that sediment that is delivered to the stream network. The ecosystem service is quantified as the amount of sediment retained by the landscape.

The model's core calculations are based on the Revised Universal Soil Loss Equation (RUSLE), which requires several spatial inputs:

• Rainfall erosivity (R factor): We used a high-resolution dataset of rainfall erosivity for Europe (Panagos et al., 2017), which represents the potential of rainfall to cause soil erosion. This factor implicitly integrates human demand, as the ecosystem service is most needed in areas where the erosive force of rain is highest.
• Soil erodibility (K factor): We used the corresponding European soil erodibility dataset from the European Soil Data Centre (ESDAC) (Panagos et al., 2014), which quantifies the susceptibility of different soil types to erosion.
• Topographic factor (LS factor): This factor, which combines slope length (L) and slope steepness (S), was calculated by the model directly from our 5m Digital Elevation Model. We set the maximum L value to 122m as per model recommendations.

• Cover-management factor (C factor) and support practice factor (P factor): These factors represent the effect of land cover and agricultural practices on erosion. We derived them from a table assigning specific C and P values to each land use/land cover class, based on literature values.

To model sediment transport and deposition, the SDR model requires additional parameters that define the connectivity between hillslopes and the stream network:

Threshold flow accumulation (TFA): We used a TFA of 400 pixels to define the threshold at which overland flow begins to concentrate and form channels. This value determines the extent of the stream network generated by the model.

• Borselli $k$ and $IC_0$ parameters: These are calibration parameters that relate the landscape's hydrological connectivity to its sediment trapping efficiency. We used the default recommended values of $k=2$ and $IC_0=0.8$, which are commonly used in European contexts.
• The model's final output is a map of sediment retained per pixel (in tons/pixel/year). This map directly represents the ecosystem service of erosion control, with higher values indicating a greater capacity of the landscape to prevent soil from reaching waterways.

##### vi. Heatwave Mitigation

We used the InVEST Urban Cooling model to estimate the cooling capacity of natural and semi-natural features during extreme heat events. The model quantifies the cooling effect of green spaces based on their shading, evapotranspiration, and albedo, and how this effect extends into the surrounding landscape.

The model required several spatial and non-spatial inputs:

• Land Use/Land Cover: We used our 5m resolution regional LULC map.
• Reference evapotranspiration ($ET_0$): We used a raster layer of reference evapotranspiration from the CHELSA database (Karger et al., 2021).
• Biophysical table: Each LULC class was assigned specific parameters: the crop coefficient ($K_c$), albedo, and shade (Table S5), compiled from an extensive literature review.
• Air blending distance: We set this parameter, which defines the distance over which cooling effects blend, to 500m, following model recommendations.
• Maximum cooling distance: This parameter defines the maximum distance over which a large green area (>2 ha) exerts a cooling effect. We used a standard value of 450m.
• Reference air temperature and urban heat island intensity: To ground the model in local climatic conditions, we used data from a recent, detailed study of the Grenoble urban heat island (Foissard et al., 2024). Based on this work, we set the reference rural air temperature to 30 °C and the maximum UHI intensity to 4.4 °C, representing typical heatwave conditions.

By incorporating the empirically measured local heat island intensity, the model's output is implicitly weighted by demand. The cooling capacity is calculated as being higher in areas with higher ambient temperatures (i.e., within the urban heat island), thus reflecting the locations where this ecosystem service is most critically needed. The final output is a map of temperature mitigation (in °C), representing the cooling contribution of each pixel to the surrounding landscape.

Table S 5. Biophysical table associated with the heatwave mitigation model. Albedo represents the albedo of each land use/land cover type. Kc refers to the evapotranspiration coefficients for each land use/land cover type. Green area indicates whether the area is natural or not. Shade represents the shade ratio for each land use/land cover type.

| CODE | Description | Albedo | Kc | Green area | Shade | References |
| --- | --- | --- | --- | --- | --- | --- |
| 11000 | Artificial surfaces and constructions | 0.143 | 0.325 | 0 | 0 | Trlica et al., (2017) |
| 11100 | Dense settlement area | 0.143 | 0.325 | 0 | 0 |  |
| 11200 | Low density settlement area | 0.147 | 0.25 | 0 | 0 |  |
| 11300 | Builtup area | 0.143 | 0.325 | 0 | 0 |  |
| 11400 | Open settlement area | 0.147 | 0.25 | 0 | 0 |  |
| 12100 | Industrial and commercial zones | 0.132 | 0.35 | 0 | 0 |  |
| 12210 | Roads motorways and trunks | 0.139 | 0.3 | 0 | 0 |  |
| 12220 | Road networks | 0.139 | 0.3 | 0 | 0 |  |
| 12221 | Roads tertiary and others | 0.139 | 0.3 | 0 | 0 |  |
| 12230 | Railways train tracks | 0.139 | 0.3 | 0 | 0 |  |
| 12240 | Unpaved roads and tracks | 0.132 | 0.31 | 0 | 0 |  |
| 14100 | Green urban areas | 0.136 | 0.27 | 1 | 0 | average (31100, 31200, 31300, 32000, 32100) |
| 21000 | Cultivated areas - Arable land - Annual crops | 0.180 | 0.912 | 1 | 0 | Average annual crops |
| 21211 | Common wheat | 0.186 | 0.89 | 1 | 0 | Sieber et al. (2022), Kang et al. (2003) |
| 21212 | Durum wheat | 0.186 | 0.8 | 1 | 0 | Lhomme et al. (2009) |
| 21213 | Barley | 0.176 | 1.07 | 1 | 0 | Sieber et al. (2022), Attarod et al. (2009) |
| 21214 | Rye | 0.181 | 0.87 | 1 | 0 | Sieber et al. (2022), |
| 21215 | Oats | 0.172 | 0.867 | 1 | 0 | Andréasson, (2023), Moteva et al. (2014) |
| 21216 | Maize | 0.184 | 0.973 | 1 | 0 | Bsaibes et al. (2009), Kang et al. (2003) |
| 21218 | Triticale | 0.183 | 0.978 | 1 | 0 | Moteva et al. (2014) |
| 21221 | Potatoes | 0.214 | 0.65 | 1 | 0 | Paredes et al. (2018) |
| 21222 | Sugar beet | 0.179 | 0.918 | 1 | 0 | Sieber et al. (2022), Moteva et al. (2014) |
| 21230 | Other non-permanent industrial crops | 0.180 | 0.912 | 1 | 0 | 21000 |
| 21231 | Sunflower | 0.225 | 0.79 | 1 | 0 | Srivastava et al. (1998), Mila et al. (2016) |
| 21232 | Rape and turnip rape | 0.188 | 0.945 | 1 | 0 | Sieber et al. (2022), Moteva et al. (2014) |
| 21233 | Soya | 0.239 | 0.86 | 1 | 0 | Weiss et al. (2001), Attarod et al. (2009) |
| 21240 | Dry pulses | 0.214 | 0.8 | 1 | 0 | Nandi et al. (2024) |
| 21250 | Fodder crops (cereals and leguminous) | 0.214 | 0.884 | 1 | 0 | Weiss et al. (2001), Moteva et al. (2014) |
| 21290 | Bare arable land | 0.23 | 0.275 | 1 | 0 | 33100 |
| 22000 | Permanent crops | 0.115 | 1.075 | 1 | 0.7 | Schwaab et al., (2015) |
| 22100 | Vineyard | 0.2 | 0.86 | 1 | 0 | Galleguillos et al. (2011), Cancela et al. (2010) |
| 22200 | Orchard | 0.179 | 0.918 | 1 | 0 | Sieber et al. (2022), Moteva et al. (2014) |
| 23100 | Managed grassland - Pastures | 0.196 | 0.93 | 1 | 0 | Rosset et al. (2001), |
| 23200 | Seminal grassland - Meadows | 0.168 | 0.93 | 1 | 0 | Ambrosi et al. (2024) |
| **31100** | Broadleaf tree cover | 0.126 | 1.55 | 1 | 1 | Schwaab et al., (2015) |
| **31102** | Broadleaf tree cover 30-60% | 0.126 | 1.55 | 1 | 0.45 |  |
| **31103** | Broadleaf tree cover 60-100% | 0.126 | 1.55 | 1 | 0.8 |  |
| **31200** | Coniferous tree cover | 0.104 | 1 | 1 | 1 |  |
| **31202** | Coniferous tree cover 30-60% | 0.104 | 1 | 1 | 0.45 |  |
| **31203** | Coniferous tree cover 60-100% | 0.104 | 1 | 1 | 0.8 |  |
| **31300** | Mixed tree cover | 0.115 | 1.4 | 1 | 1 |  |
| **31400** | Tree cover in agricultural context | 0.115 | 1.075 | 1 | 0.7 | 14100 |
| **31450** | Tree cover in urban context | 0.136 | 0.27 | 1 | 0.7 |  |
| **31500** | Green linear elements - linear woody features | 0.132 | 0.975 | 1 | 0.4 | Aartsma et al., (2020) |
| **31600** | Patchy woody features | 0.132 | 0.975 | 1 | 0 |  |
| **31610** | Additional woody features | 0.132 | 0.975 | 1 | 0 |  |
| **32000** | Scrub and shrubland | 0.132 | 0.975 | 1 | 0 |  |
| **32100** | Alpine and sub-alpine natural grassland | 0.194 | 1.125 | 1 | 0 | Tian et al., (2014);<br>Blumthaler and Ambach, (1988) |
| **32200** | Moors and heathland - other scrubland | 0.132 | 0.975 | 1 | 0 | Aartsma et al., (2020) |
| **32300** | Sclerophyllous vegetation | 0.132 | 0.65 | 1 | 0 |  |
| **33100** | Beaches, dunes, sands | 0.23 | 0.275 | 1 | 0 | Blumthaler and Ambach, (1988) |
| **33200** | Bare rocks and rock debris | 0.3 | 0.125 | 1 | 0 | Al Fahdawi, (2021) |
| **33300** | Sparsely vegetated land | 0.13 | 0.55 | 1 | 0 | Sieber et al., (2022) |
| **33500** | Permanent snow covered surfaces | 0.507 | 0.95 | 1 | 0 | Kalitin, (1930) |
| **41000** | Wetland (permanent wet areas) - inland marshes | 0.159 | 0.625 | 1 | 0 | Trlica et al., (2017) |
| **51000** | Water bodies | 0.091 | 0.95 | 1 | 0 | Blumthaler and Ambach, (1988) |
| **51100** | Rivernetwork | 0.091 | 0.95 | 1 | 0 |  |
| **51200** | Riverbed > 10m width | 0.091 | 0.95 | 1 | 0 |  |

### vii. Pollination

The pollination ecosystem service was modeled to represent the potential for wild pollinators to provide pollination to crops, using a connectivity-based approach centered on a pollinator archetype. This archetype represents a typical wild bee in the region, defined by its habitat preferences and foraging behaviour, with parameters detailed in Table S6.

First, we created a categorical habitat suitability map at a 5m resolution. Following Schulp et al. (2014), we classified each land use/land cover pixel into one of three categories: 'full habitat' (e.g., scrublands, diverse grasslands), 'partial habitat' (e.g., managed pastures, forest edges within 10m of open areas), or 'non-habitat' (e.g., urban areas, dense forests, croplands).

Second, we converted this habitat suitability map into a resistance surface for pollinator movement. Instead of a simple reclassification, the resistance of 'non-habitat' pixels was calculated as a function of their distance to the nearest source of pollinators. We used an exponential decay function where resistance increases with distance from the nearest 'full' or 'partial' habitat pixel. The decay rate ($r \approx 1.4$) was calibrated so that the function's value drops by half at a distance of 500m, corresponding to the assumed median foraging distance of our pollinator archetype.

Third, we modeled landscape connectivity using the Omniscape algorithm. Using the resistance surface described above, Omniscape calculated the cumulative current flow across the landscape, with a moving window radius set to 500m to reflect the pollinator's average foraging range. The resulting map represents the probability of pollinator passage, or "pollinator flux," across every pixel in the landscape.

Finally, we integrated human demand. The pollinator flux map was multiplied by a demand layer that represents the degree to which different crops depend on insect pollination. Following Klein et al. (2007) and Schulp et al. (2014), we classified each crop land use/land cover class as having full, partial, or no dependence on pollination. The final ecosystem service map thus represents the realized pollination service, showing the flux of pollinators specifically in the areas where it is needed by agriculture.

*Table S 6. Suitability habitat for pollinators and demand for pollination scores used for the pollination model.*

| **CODE** | **DESC** | **Pollinator** | **Demand of pollination** |
| --- | --- | --- | --- |
| **21222** | Sugar beet | 0 | 1 |
| **21231** | Sunflower | 0 | 1 |
| **21232** | Rape and turnip rape | 0 | 1 |
| **21233** | Soya | 0 | 1 |
| **21240** | Dry pulses | 0 | 1 |
| **21250** | Fodder crops (cereals and leguminous) | 0 | 1 |
| **21290** | Bare arable land | 0 | 0 |
| **22000** | Permanent crops | 0 | 1 |
| **22100** | Vinyard | 0 | 1 |
| **22200** | Orchard | 0.5 | 1 |
| **23100** | Managed grassland - Pastures | 0.5 | 0.5 |
| **23200** | Seminal grassland - Meadows | 0.5 | 0.5 |
| **31100** | Broadleaf tree cover | 0.5 | 0.5 |
| **31102** | Broadleaf tree cover 30-60% | 0.5 | 0.5 |
| **31103** | Broadleaf tree cover 60-100% | 0.5 | 0.5 |
| **31200** | Coniferous tree cover | 0.5 | 0.5 |
| **31202** | Coniferous tree cover 30-60% | 0.5 | 0.5 |
| **31203** | Coniferous tree cover 60-100% | 0.5 | 0.5 |
| **31300** | Mixed tree cover | 0.5 | 0.5 |
| **32000** | Scrub and shrubland | 1 | 0.5 |
| **32100** | Alpine and sub-alpine natural grassland | 0.5 | 0.5 |
| **32200** | Moors and heathland - other scrubland | 1 | 0.5 |
| **32300** | Sclerophyllous vegetation | 1 | 0.5 |

##### c. Specific connectivity-based models: Species selection and demand integration

The following five ecosystem services were modelled using the general connectivity-based framework described in S2.1.2. While the core connectivity modelling was consistent, each ecosystem service differed in its constituent species assemblage and, crucially, in how human demand was conceptualized and integrated.

###### i. Mosquito control

This model is based on averaging suitability-weighted connectivity (or suitability alone) maps of a group of 81 species potentially linked to mosquito control (see Table S7 for species). To update the model and incorporate the demand and access of this ecosystem service, we used mosquito observation points for *Aedes albopictus* and *Culex pipiens*. These two species were selected because they are the most directly associated with potential human nuisance in the region. While other species may also cause nuisance, they are more commonly related to animals than to humans.

We contacted the Interdepartmental Mosquito Control Authority in Grenoble (EID Rhône-Alpes) to obtain data on the presence of these two species. This institution is responsible for monitoring mosquito populations and notifying potential health risks linked to mosquitoes. We obtained two datasets: one for the presence of *Aedes albopictus*, which included an Excel file with 22,958 recorded observation points, and another for *Culex pipiens*, with 3,204 recorded points.

After clipping the data to fit the study area, we retained a total of 4,318 points for *Aedes albopictus* and 892 for *Culex pipiens*. We merged both datasets and applied a habitat modelling approach using Biomod2 (Thuiller et al., 2016) following the methodology in Sherpa et al. (2020) to create a raster layer representing the habitat suitability for both species as a measure of demand for mosquito control.

Additionally, we created a 1 km buffer around urban areas, recognizing that human mobility is a key factor in mosquito exposure. This buffer distance was informed by a participatory mapping approach conducted in Grenoble, where over 600 participants were asked about various ecosystem-related benefits, including their willingness to travel to nearby ecosystems for regular recreational activities. To create the buffer, we used the land use-land cover layer and selected the codes 11100, 11200, 11300, 11400, 12100 corresponding with areas where people live or work. Finally, we multiplied the resulting layer by the connectivity map for mosquito control species to obtain the final mosquito control map.

### ii. Biological control

This ecosystem service represents the regulation of agricultural and garden pests by their natural predators.

• Species assemblage: The model aggregated the connectivity maps of 78 vertebrate species (including various insectivorous birds, bats, and small mammals) known to prey on common pests in the region (Table S7).
• Demand integration: The ecosystem service is only relevant where pests have a direct impact on human activities. Therefore, demand was modelled as spatially explicit and concentrated. We created a binary mask from the land use/land cover map, selecting all pixels corresponding to agricultural lands (e.g., croplands, orchards) and urban green spaces (e.g., parks, private gardens). The final ecosystem service map was generated by multiplying the aggregated species connectivity map by this demand mask, thus restricting the ecosystem service to areas where it is explicitly needed.

### iii. Hunting value

This ecosystem service represents the opportunity for recreational hunting, a culturally significant activity in the region.

• Species assemblage: The model included 24 species that are officially listed as game species by regional hunters' associations (Table S7).
• Demand integration: The value of this ecosystem service is determined not only by the presence of game species but also by the legal and physical accessibility for hunters. Demand was therefore integrated as a multi-layered weighting scheme based on regional hunting regulations. We created a raster where different zones were assigned different weights: areas where hunting is explicitly prohibited (e.g., core zones of nature reserves) received a weight of 0.1; hunting reserves with restricted access received a weight of 0.2; regulatory buffers around buildings (150m) received a weight of 0.5; and all other permissible areas retained their full value (weight of 1.0). The final ecosystem service map was created by multiplying the aggregated connectivity map by this demand layer.

##### iv. Emblematic species

This ecosystem service represents the cultural, aesthetic, and existence value derived from the presence of iconic and charismatic species.

• Species assemblage: We selected 96 species considered emblematic based on a combination of criteria, including their conservation status (e.g., listed in the EU Habitats and Birds Directives), their importance for nature-based tourism, and high public interest as indicated by citizen science observation rates (O'Connor et al., in revision).
• Demand integration: The value people derive from emblematic species is not confined to specific locations of observation but extends across the entire landscape as part of the region's natural heritage (i.e., existence value). Consequently, demand for this ecosystem service was considered ubiquitous and non-spatial. No demand mask or weighting was applied, and the aggregated species connectivity map was used directly as the final ecosystem service map.

##### v. Seed dispersal

This ecosystem service represents a foundational ecological process that underpins the resilience and regeneration of plant communities across the landscape.

• Species assemblage: The model aggregated the connectivity maps of 81 frugivorous species (birds and mammals) known to play a key role in dispersing native plant seeds (Table S7).
• Demand integration: Similar to emblematic species, seed dispersal provides a benefit that is not targeted at a specific human activity but is essential for the long-term health of all natural and semi-natural ecosystems. Its importance is distributed across the landscape wherever ecosystem regeneration is needed. Therefore, demand was also considered ubiquitous and non-spatial. The aggregated species connectivity map was used directly as the final ecosystem service map.

Table S 7. Species used for each ecosystem service based on connectivity algorithms. "Total ES" indicates the number of times a species appears in different ecosystem services.

| Species | Hunting value | Biological control | Seed dispersal | Mosquito control | Emblematic species | Total ES |
| --- | --- | --- | --- | --- | --- | --- |
| *Acanthis hornemanni* |  |  | x |  |  | 1 |
| *Accipiter gentilis* |  | x |  |  | x | 2 |
| *Accipiter nisus* |  |  |  |  | x | 1 |
| *Aegolius funereus* |  | x |  | x | x | 3 |
| *Aegyptius monachus* |  |  |  |  | x | 1 |
| *Alauda arvensis* |  | x |  |  |  | 1 |
| *Alcedo atthis* |  |  |  |  | x | 1 |
| *Alectoris graeca* |  |  | x |  | x | 2 |
| *Alectoris rufa* | x |  | x |  |  | 2 |
| *Alytes obstetricans* |  | x |  |  | x | 2 |
| *Anas platyrhynchos* | x |  | x |  |  | 2 |
| *Anthus campestris* |  |  |  |  | x | 1 |
| *Anthus pratensis* |  | x |  |  |  | 1 |
| *Anthus spinoletta* |  |  | x |  |  | 1 |
| *Apodemus alpicola* |  |  | x |  |  | 1 |
| *Apodemus flavicollis* |  |  | x |  |  | 1 |
| *Apodemus sylvaticus* |  |  | x |  |  | 1 |
| *Apus apus* |  | x |  | x |  | 2 |
| *Apus melba* |  | x |  | x |  | 2 |
| *Aquila chrysaetos* |  | x |  |  | x | 2 |
| *Aquila fasciata* |  |  |  |  | x | 1 |
| *Ardea purpurea* |  |  |  |  | x | 1 |
| *Asio otus* |  | x |  |  |  | 1 |
| *Athene noctua* |  | x |  | x |  | 2 |
| *Aythya ferina* | x |  |  |  |  | 1 |
| *Barbastella barbastellus* |  |  |  | x | x | 2 |
| *Bombina variegata* |  |  |  | x | x | 2 |
| *Botaurus stellaris* |  |  |  |  | x | 1 |
| *Bubo bubo* |  | x |  |  | x | 2 |
| *Burhinus oedicephalus* |  | x |  |  | x | 2 |
| *Buteo buteo* |  | x |  | x |  | 2 |
| *Canis lupus* |  | x |  |  | x | 2 |
| *Capra ibex* |  |  |  |  | x | 1 |
| *Capreolus capreolus* | x |  |  |  | x | 2 |
| *Caprimulgus europaeus* |  | x |  | x | x | 3 |
| *Carduelis carduelis* |  |  | x |  |  | 1 |
| *Castor fiber* |  |  |  |  | x | 1 |
| *Cecropis daurica* |  | x |  | x |  | 2 |
| *Certhia brachydactyla* |  |  |  |  | x | 1 |
| *Cervus elaphus* | x |  | x |  | x | 3 |
| *Chloris chloris* |  |  | x |  |  | 1 |
| *Chroicocephalus ridibundus* |  |  |  | x |  | 1 |
| *Circaetus gallicus* |  |  |  |  | x | 1 |

|  |  |  |  |  |  |
| --- | --- | --- | --- | --- | --- |
| *Circus aeruginosus* |  | x |  | x | 2 |
| *Circus cyaneus* |  | x |  | x | 2 |
| *Circus pygargus* |  | x |  | x | 2 |
| *Columba livia* | x |  | x |  | 2 |
| *Columba oenas* | x |  | x |  | 2 |
| *Columba palumbus* | x |  | x | x | 3 |
| *Coronella austriaca* |  |  |  | x | 1 |
| *Corvus corax* |  | x | x | x | 3 |
| *Corvus corone* |  | x | x |  | 2 |
| *Corvus frugilegus* |  | x | x |  | 2 |
| *Corvus monedula* |  | x | x |  | 2 |
| *Coturnix coturnix* | x |  | x |  | 2 |
| *Crex crex* |  |  |  | x | 1 |
| *Crocidura russula* |  | x |  |  | 1 |
| *Crocidura suaveolens* |  | x |  |  | 1 |
| *Cuculus canorus* |  | x |  | x | 2 |
| *Cyanistes caeruleus* |  |  | x |  | 1 |
| *Dama dama* | x |  |  | x | 2 |
| *Delichon urbicum* |  | x |  | x | 2 |
| *Dendrocopos major* |  |  | x | x | 2 |
| *Dendrocoptes medius* |  |  | x |  | 1 |
| *Dryocopus martius* |  |  |  | x | 1 |
| *Egretta garzetta* |  |  |  | x | 1 |
| *Eliomys quercinus* |  |  | x |  | 1 |
| *Emberiza calandra* |  | x |  |  | 1 |
| *Emberiza cia* |  |  |  | x | 1 |
| *Emberiza cirius* |  |  | x |  | 1 |
| *Emberiza citrinella* |  | x |  |  | 1 |
| *Emberiza hortulana* |  | x |  | x | 3 |
| *Emberiza schoeniclus* |  |  | x | x | 2 |
| *Emys orbicularis* |  |  |  | x | 1 |
| *Epidalea calamita* |  | x |  |  | 1 |
| *Eptesicus nilssonii* |  |  |  | x | 2 |
| *Eptesicus serotinus* |  |  |  | x | 2 |
| *Erinaceus europaeus* |  | x |  |  | 1 |
| *Erithacus rubecula* |  |  | x |  | 1 |
| *Falco peregrinus* |  |  |  | x | 1 |
| *Falco subbuteo* |  | x |  | x | 2 |
| *Falco tinnunculus* |  | x |  |  | 1 |
| *Felis silvestris* |  | x |  | x | 2 |
| *Ficedula hypoleuca* |  |  |  | x | 1 |
| *Gallinula chloropus* |  |  | x |  | 1 |
| *Garrulus glandarius* |  |  | x | x | 2 |
| *Glaucidium passerinum* |  | x |  | x | 2 |
| *Glis glis* |  | x | x |  | 2 |
| *Gyps fulvus* |  |  |  | x | 1 |

|  |  |  |  |  |
| --- | --- | --- | --- | --- |
| *Hierophis viridiflavus* | x |  |  | 1 |
| *Himantopus himantopus* |  |  | x | 1 |
| *Hippolais polyglotta* |  | x |  | 1 |
| *Hirundo rustica* |  | x |  | 1 |
| *Hyla arborea* |  |  | x | 1 |
| *Hypsugo savii* |  | x | x | 2 |
| *Ixobrychus minutus* |  |  | x | 1 |
| *Jynx torquilla* |  | x |  | 1 |
| *Lacerta agilis* | x |  | x | 2 |
| *Lagopus muta* |  | x | x | 2 |
| *Lanius collurio* | x | x | x | 3 |
| *Lanius excubitor* | x | x |  | 2 |
| *Lanius senator* | x | x |  | 2 |
| *Larus michahellis* | x | x |  | 2 |
| *Lepus europaeus* | x | x |  | 2 |
| *Lepus timidus* | x |  |  | 1 |
| *Linaria cannabina* |  | x |  | 1 |
| *Lissotriton vulgaris* |  | x | x | 2 |
| *Lullula arborea* | x |  | x | 2 |
| *Luscinia megarhynchos* |  | x |  | 1 |
| *Luscinia svecica* |  | x | x | 2 |
| *Lutra lutra* | x |  | x | 2 |
| *Lynx lynx* | x |  | x | 2 |
| *Marmota marmota* |  |  | x | 1 |
| *Martes foina* | x | x |  | 2 |
| *Martes martes* | x |  |  | 1 |
| *Meles meles* | x | x |  | 2 |
| *Merops apiaster* | x | x | x | 3 |
| *Micromys minutus* |  | x |  | 1 |
| *Microtus agrestis* |  | x |  | 1 |
| *Microtus arvalis* |  | x |  | 1 |
| *Microtus duodecimcostatus* |  | x |  | 1 |
| *Milvus migrans* | x | x | x | 3 |
| *Milvus milvus* | x | x | x | 3 |
| *Miniopterus schreibersii* |  | x | x | 2 |
| *Monticola saxatilis* |  | x |  | 1 |
| *Montifringilla nivalis* |  | x |  | 1 |
| *Motacilla alba* |  | x |  | 1 |
| *Motacilla cinerea* |  | x |  | 1 |
| *Motacilla flava* |  | x |  | 1 |
| *Muscardinus avellanarius* |  | x | x | 2 |
| *Muscicapa striata* |  | x |  | 1 |
| *Mustela erminea* | x |  |  | 1 |
| *Mustela nivalis* | x |  |  | 1 |
| *Mustela putorius* | x |  |  | 1 |
| *Myotis alcathoe* |  | x | x | 2 |
| *Myotis bechsteinii* |  | x | x | 2 |
| *Myotis blythii* |  | x | x | 2 |
| *Myotis brandtii* |  | x | x | 2 |
| *Myotis capaccinii* |  | x | x | 2 |
| *Myotis daubentonii* |  | x | x | 2 |
| *Myotis emarginatus* |  | x | x | 2 |
| *Myotis myotis* |  |  | x | 1 |
| *Myotis mystacinus* |  | x | x | 2 |
| *Myotis nattereri* |  |  | x | 1 |
| *Nucifraga caryocatactes* |  | x | x | 2 |
| *Numenius arquata* |  | x |  | 1 |
| *Nyctalus lasiopterus* |  |  | x | 2 |
| *Nyctalus leisleri* |  |  | x | 2 |
| *Nyctalus noctula* |  |  | x | 2 |
| *Nycticorax nycticorax* | x |  | x | 2 |
| *Oenanthe oenanthe* | x |  | x | 2 |
| *Oriolus oriolus* |  | x |  | 1 |
| *Oryctolagus cuniculus* | x |  | x | 2 |
| *Otus scops* |  | x | x | 2 |
| *Parus major* |  | x | x | 2 |
| *Passer domesticus* |  | x |  | 1 |
| *Passer montanus* |  | x | x | 2 |
| *Perdix perdix* | x |  | x | 3 |
| *Periparus ater* |  | x | x | 2 |
| *Pernis apivorus* | x |  | x | 3 |
| *Petronia petronia* |  | x |  | 1 |
| *Phasianus colchicus* | x |  | x | 2 |
| *Phoenicurus ochruros* |  | x | x | 2 |
| *Phoenicurus phoenicurus* |  | x | x | 2 |
| *Phylloscopus bonelli* |  |  | x | 1 |
| *Phylloscopus sibilatrix* |  | x |  | 1 |
| *Phylloscopus trochilus* |  | x |  | 1 |
| *Pica pica* | x | x | x | 3 |
| *Pipistrellus kuhlii* |  | x | x | 2 |
| *Pipistrellus nathusii* |  | x | x | 2 |
| *Pipistrellus pipistrellus* |  | x | x | 2 |
| *Plecotus auritus* |  | x | x | 2 |
| *Plecotus austriacus* |  | x | x | 2 |
| *Podarcis muralis* |  |  | x | 1 |
| *Poecile montanus* |  | x | x | 2 |
| *Porzana porzana* |  |  | x | 1 |
| *Ptyonoprogne rupestris* |  |  | x | 1 |
| *Pyrrhocorax graculus* |  | x |  | 1 |
| *Pyrhacorax pyrrhacorax* |  | x | x | 2 |
| *Pyrrhula pyrrhula* |  | x |  | 1 |
| *Rallus aquaticus* |  | x |  | 1 |

|  |  |  |  |  |  |
| --- | --- | --- | --- | --- | --- |
| *Rana dalmatina* |  |  |  | x | 1 |
| *Rhinolophus euryale* |  |  | x | x | 2 |
| *Rhinolophus ferrumequinum* |  |  | x | x | 2 |
| *Rhinolophus hipposideros* |  |  | x | x | 2 |
| *Riparia riparia* |  |  | x |  | 1 |
| *Rupicapra rupicapra* | x |  |  | x | 2 |
| *Saxicola rubetra* |  | x | x |  | 2 |
| *Sciurus vulgaris* |  |  | x |  | 1 |
| *Scolopax rusticola* | x |  | x |  | 2 |
| *Sitta europaea* |  |  | x |  | 1 |
| *Sorex araneus* |  | x |  |  | 1 |
| *Sorex coronatus* |  | x |  |  | 1 |
| *Spatula clypeata* | x |  |  |  | 1 |
| *Spatula querquedula* | x |  |  |  | 1 |
| *Spinus spinus* |  |  | x |  | 1 |
| *Streptopelia decaocto* | x |  | x |  | 2 |
| *Streptopelia turtur* | x |  | x |  | 2 |
| *Strix aluco* |  | x |  | x | 2 |
| *Sturnus vulgaris* |  | x | x |  | 2 |
| *Sus scrofa* | x |  | x | x | 3 |
| *Sylvia atricapilla* |  |  |  | x | 1 |
| *Sylvia borin* |  |  | x | x | 2 |
| *Sylvia communis* |  |  | x |  | 1 |
| *Tadarida teniotis* |  |  | x | x | 2 |
| *Talpa europaea* |  | x |  |  | 1 |
| *Tetrao tetrix* | x |  | x |  | 2 |
| *Tetrastes bonasia* | x |  | x |  | 2 |
| *Tetrax tetrax* |  |  | x | x | 2 |
| *Timon lepidus* |  | x |  |  | 1 |
| *Triturus cristatus* |  |  |  | x | 2 |
| *Troglodytes troglodytes* |  |  | x | x | 3 |
| *Turdus merula* |  | x |  |  | 1 |
| *Turdus philomelos* |  |  | x |  | 1 |
| *Turdus pilaris* |  | x | x | x | 3 |
| *Turdus torquatus* |  |  | x |  | 1 |
| *Turdus viscivorus* |  | x | x | x | 3 |
| *Tyto alba* |  | x |  |  | 1 |
| *Upupa epops* |  | x |  |  | 1 |
| *Vanellus vanellus* |  | x |  |  | 1 |
| *Vespertilio murinus* |  |  |  | x | 1 |
| *Vespertilio murinus.* |  |  | x |  | 1 |
| *Vipera aspis* |  | x |  |  | 1 |
| *Vulpes vulpes* |  | x | x |  | 2 |
| *Zamenis longissimus* |  | x |  |  | 1 |
| *Zootoca vivipara* |  |  |  | x | 1 |
| *Zootoca vivipara.* |  | x |  |  | 1 |

#### C. Normalization methods

To enable direct quantitative comparison and aggregation of the diverse ecosystem service indicators, which originated from different models with varying units and ranges, a rigorous normalization protocol was implemented. This protocol ensures all indicators are converted to a common, dimensionless scale from 0 to 1, allowing for valid comparisons across ecosystem services and scenarios. Prior to normalization, all raster layers were spatially harmonized to a common grid (5m resolution, EPSG:2154) to ensure perfect pixel alignment.

##### a. Normalization strategy: Global linear decile normalization

A simple layer-by-layer normalization (e.g., min-max scaling for each individual map) was discarded as it would prevent valid comparisons across scenarios; a value of 1.0 in the Present-day scenario would not represent the same absolute level of ecosystem service as a 1.0 in a future scenario.

Instead, we applied a Global Linear Decile Normalization with Range Clipping (D1-D9). This method establishes a common and robust scale for each ecosystem service "family" across all scenarios (Present, Participatory, and Technical). The protocol was as follows:

• Data aggregation: For each ecosystem service (e.g., all "Carbon Storage" layers), all valid pixel values from all three scenarios were combined into a single global dataset.
• Global Threshold Calculation: From this aggregated dataset, the lower decile (D1, 10th percentile) and the upper decile (D9, 90th percentile) were calculated. Using deciles instead of absolute minimum and maximum values avoids the influence of extreme outliers and provides a robust range of variation for each ecosystem service.
• Application of normalization: Each individual layer within the ecosystem service family was then normalized using these same fixed D1 global and D9 global thresholds according to the following rules:
  ○ If original value ≤ D1 global, then normalized value = 0.
  ○ If original value ≥ D9 global, then normalized value = 1.
  ○ If D1 global < original value < D9 global, the value was linearly scaled to the [0, 1] range using the formula: $\text{normalized value} = (\text{original value} - \text{D1 global}) / (\text{D9 global} - \text{D1 global})$

This global approach ensures that a normalized value (e.g., 0.8) represents the same absolute level of ecosystem service provision in any scenario, allowing for valid cross-scenario and temporal comparisons.

##### b. Treatment of special cases

During the analysis, two specific data distributions required special treatment to avoid biased results:

• Zero-inflated data: For ecosystem services such as "Erosion Control" or "Mosquito Control," where a large number of pixels with a value of zero represented the "absence of service," the calculation of the global deciles was performed exclusively on the subset of pixels with values strictly greater than zero. This prevented the abundance of zeros from distorting the

thresholds. Pixels with an original value of zero were maintained as zero in the final normalized layer.

• Categorical or pre-normalized data: The "Pollination" service exhibited discrete values (0, 0.5, and 1). In this case, the decile-based normalization did not alter the data (D1=0, D9=1), and the original values were accepted as an indicator already standardized on a 0-to-1 scale.

##### **D. Spatial gradient analysis (detailed methods)**

This section details the multi-step framework used to quantify and classify ecosystem service gradients across protected area borders.

###### **a. Border segmentation and transect generation**

To analyse the variable interfaces of protected areas, we first simplified the protected area polygons using the Douglas-Peucker algorithm (simplification tolerance = 300) to facilitate subsequent geometric operations. The border of each simplified polygon was then divided into a set number of equal-length segments. These segments serve as our primary unit of analysis, allowing us to capture how gradient patterns change along a perimeter. To maintain a comparable level of detail across sites of varying sizes, the number of segments was scaled to the protected area's size: 20 segments for large (>10,000 ha), 10 for medium (1,000-10,000 ha), 5 for small (100-1,000 ha), and 3 for very small (<100 ha) protected areas.

Within each segment, we then generated high-density perpendicular transects at 25m intervals, extending a maximum of 4 km both inwards and outwards from the border. These transects served as fine-scale sampling lines. Before data extraction, all transects were subjected to a rigorous automated filtering process to ensure their spatial validity. The filtering rules included: discarding transects shorter than a 20m minimum length; truncating transects that intersected with the structural buffers of neighbouring protected areas; and discarding transects with geometric anomalies (e.g., crossing their own protected area polygon more than once).

###### **b. Structural buffer creation**

Conventional linear-distance buffers are poorly suited for heterogeneous landscapes, as they fail to capture real-world complexity in accessibility and ecological connectivity. To overcome this, we developed structural buffers based on landscape resistance to movement. These buffers define concentric zones of equal "cost of travel," ensuring that all points within a given buffer zone are functionally equidistant from the protected area border in terms of landscape permeability.

The entire cost-distance analysis was performed iteratively using Python libraries (NumPy, GDAL, scikit-image, and SciPy). The process was conducted twice to assess these structural gradients both outwards from (positive buffers) and inwards towards (negative buffers) protected areas.

###### **Generation of the structural resistance surface**

First, a raster representing landscape resistance was configured. We assigned a value of zero within the source area (e.g., inside the protected area for the outward iteration), designating it as the least-cost origin. For all other pixels, the resistance value was calculated as a composite of land cover and topographic factors using the following formula:

$$\text{Final Resistance} = 2 \times (\text{Land-Cover Resistance}) + \text{Topographic Resistance}$$

Where:

• Land-Cover Resistance: Each LULC class was assigned a specific resistance value (ranging from 0.1 to 1.0) reflecting its permeability to general movement (see Table S10 for detailed values).
• Topographic Resistance: Derived from a 5m Digital Elevation Model (DEM), classified into three categories based on slope steepness (<50%, 50-100%, and >100%) to penalize movement across steep terrain.

##### Cumulative cost calculation and buffer delineation

Following the creation of the resistance surface, we executed the Geometric Minimum Cost Path algorithm from scikit-image (skimage.graph.mcp) to generate a cumulative cost raster. This raster represents the total accumulated resistance to travel from the protected area boundary to any given pixel. Finally, this cumulative cost raster was used to delineate five discrete structural buffers. The zones were defined based on the following cumulative resistance thresholds, creating zones of progressively increasing isolation:

• Buffer 1 / -1: Cumulative Resistance $\leq 100$
• Buffer 2 / -2: Cumulative Resistance 101 – 200
• Buffer 3 / -3: Cumulative Resistance 201 – 400
• Buffer 4 / -4: Cumulative Resistance 401 – 800
• Buffer 5 / -5: Cumulative Resistance $> 800$

These non-linear intervals were selected to provide higher resolution near the border (where gradients change rapidly) while capturing the broader landscape context in the outer zones.

*Table S 8. Resistances assigned to each land use/land cover class to create the resistance layer for structural buffer calculations.*

| CODE | DESC | Cost |
| --- | --- | --- |
| 11000 | Artificial surfaces and constructions | 1 |
| 11100 | Dense settlement area | 1 |
| 11200 | Low density settlement area | 0.9 |
| 11300 | Builtup area | 1 |
| 11400 | Open settlement area | 0.9 |
| 12100 | Industrial and commercial zones | 1 |
| 12210 | Roads motroways and trunks | 1 |
| 12220 | Road networks | 1 |
| 12221 | Roads tertiary and others | 1 |
| 12230 | Railways train tracks | 0.9 |
| 12240 | Unpaved roads and tracks | 0.8 |
| 14100 | Green urban areas | 0.7 |
| **21000** | Cultivated areas - Arable land - Annual crops | 0.5 |
| **21211** | Common wheat | 0.5 |
| **21212** | Durum wheat | 0.5 |
| **21213** | Barley | 0.5 |
| **21214** | Rye | 0.5 |
| **21215** | Oats | 0.5 |
| **21216** | Maize | 0.5 |
| **21218** | Triticale | 0.5 |
| **21221** | Potatoes | 0.5 |
| **21222** | Sugar beet | 0.5 |
| **21230** | Other non-permanent industrial crops | 0.5 |
| **21231** | Sunflower | 0.5 |
| **21232** | Rape and turnip rape | 0.5 |
| **21233** | Soya | 0.5 |
| **21240** | Dry pulses | 0.5 |
| **21250** | Fodder crops (cereals and leguminous) | 0.5 |
| **21290** | Bare arable land | 0.5 |
| **22000** | Permanent crops | 0.5 |
| **22100** | Vinyard | 0.5 |
| **22200** | Orchard | 0.5 |
| **23100** | Managed grassland - Pastures | 0.4 |
| **23200** | Seminatural grassland - Meadows | 0.4 |
| **31100** | Broadleaf tree cover | 0.1 |
| **31102** | Broadleaf tree cover 30-60% | 0.1 |
| **31103** | Broadleaf tree cover 60-100% | 0.1 |
| **31200** | Coniferous tree cover | 0.1 |
| **31202** | Coniferous tree cover 30-60% | 0.1 |
| **31203** | Coniferous tree cover 60-100% | 0.1 |
| **31300** | Mixed tree cover | 0.1 |
| **31400** | Tree cover in agricultural context | 0.3 |
| **31450** | Tree cover in urban context | 0.8 |
| **31500** | Green linear elements - linear woody features | 0.2 |
| **31600** | Patchy woody features | 0.2 |
| **31610** | Additional woody features | 0.2 |
| **32000** | Scrub and shrubland | 0.1 |
| **32100** | Alpine and sub-alpine natural grassland | 0.1 |
| **32200** | Moors and heathland - other scrubland | 0.1 |
| **32300** | Sclerophyllous vegetation | 0.1 |
| **33100** | Beaches, dunes, sands | 0.1 |
| **33200** | Bare rocks and rock debris | 0.2 |
| **33300** | Sparsely vegetated land | 0.2 |
| **33500** | Permanent snow covered surfaces | 0.4 |
| **41000** | Wetland (permanent wet areas) - inland marshes | 0.1 |
| **51000** | Water bodies | 0.3 |
| **51100** | Rivernetwork | 0.3 |
| **51200** | Riverbed > 10m width | 0.2 |

#### c. Gradient Profile Construction and Typology Generation

For each border segment and ecosystem service bundle, we constructed a single, representative gradient profile. This was achieved by averaging the normalized bundle values from all valid transects within that segment for each of the 10 discrete structural buffers. This process yields a series of 10 data points representing the average gradient.

Finally, we developed an automated method to classify the shape of these profiles using polynomial regression. For each profile, we fitted both first-degree (linear) and second-degree (quadratic) models, selecting the best fit based on minimizing the Mean Squared Error (MSE) (Fig. S3). A first-degree polynomial was considered a good fit if $MSE < 0.05$. Border segments were then classified into one of five types based on the selected model's coefficients:

• Type A ('Flat Gradient'): Linear fit with an absolute slope $< 0.01$.
• Type B ('Increasing Gradient'): Linear fit with a slope $\geq 0.01$.
• Type C ('Decreasing Gradient'): Linear fit with a slope $\leq -0.01$.
• Type D ('Boundary Depression'): Quadratic fit with a positive quadratic term (concave-up).
• Type E ('Interface Peak'): Quadratic fit with a negative quadratic term (concave-down).

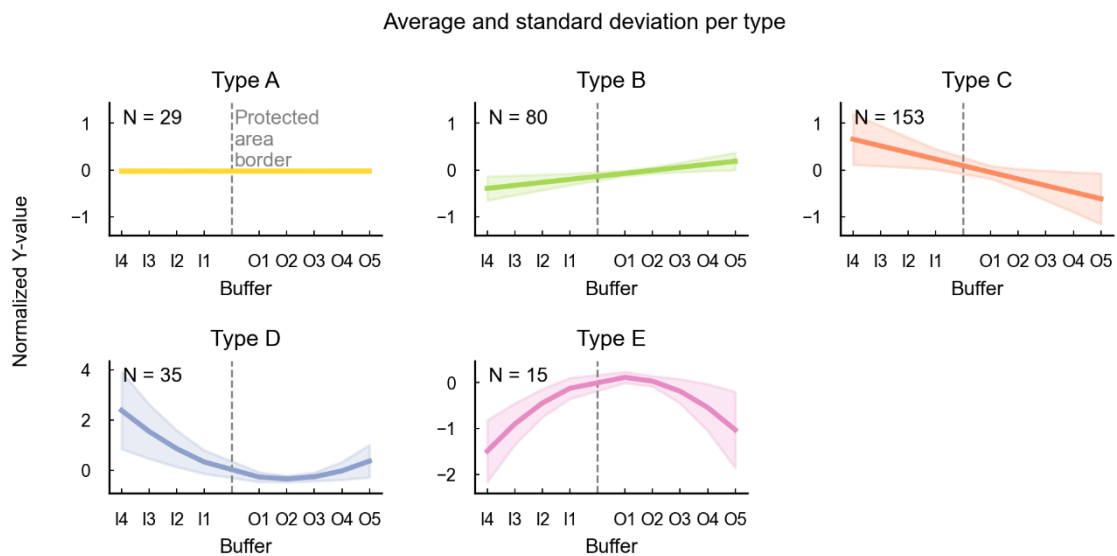

Figure S3. Typology of protected area borders based on ecosystem service bundle profiles (from González-García et al., 2026). The profiles show normalized ecosystem service bundle values (Y-axis) across structural buffer zones from inside to outside protected areas (X-axis). Y-axis values represent the average bundle value in each buffer zone, normalized by the average value along the entire border-segment transect line. X-axis labels use "I" to indicate zones inside protected areas and "O" for zones outside. The number of border segments classified into each typology (N) is shown in each panel. Because profiles are provided for the three analysed bundles (rural, cultural, and urban), the figure includes 312 cases in total (104 border segments × 3 bundles). Figure S3 in the Supporting Information provides an example of the polynomial regressions generated for all bundles and border segments, including statistically significant values and detailed mean square errors. Figure from González-García et al. (2026).

#### d. Illustrative outputs of the gradient analysis

To provide a visual example of the analysis outputs, we include two figures. Figure S4 shows the full set of polynomial regressions generated for a single, representative border segment across all 12

ecosystem services and 3 bundles. This illustrates how the classification algorithm works in practice, showing the fitted curves, the resulting classification (e.g., Type D), and the associated statistical fit (MSE). Figure S5 provides a spatial representation of the results for an entire protected area. The figure displays heatmaps illustrating the distribution of ecosystem services and bundle values as a function of distance from the border for each segment, allowing for a visual interpretation of the spatial patterns of ecosystem service provision inside and outside the protected area.

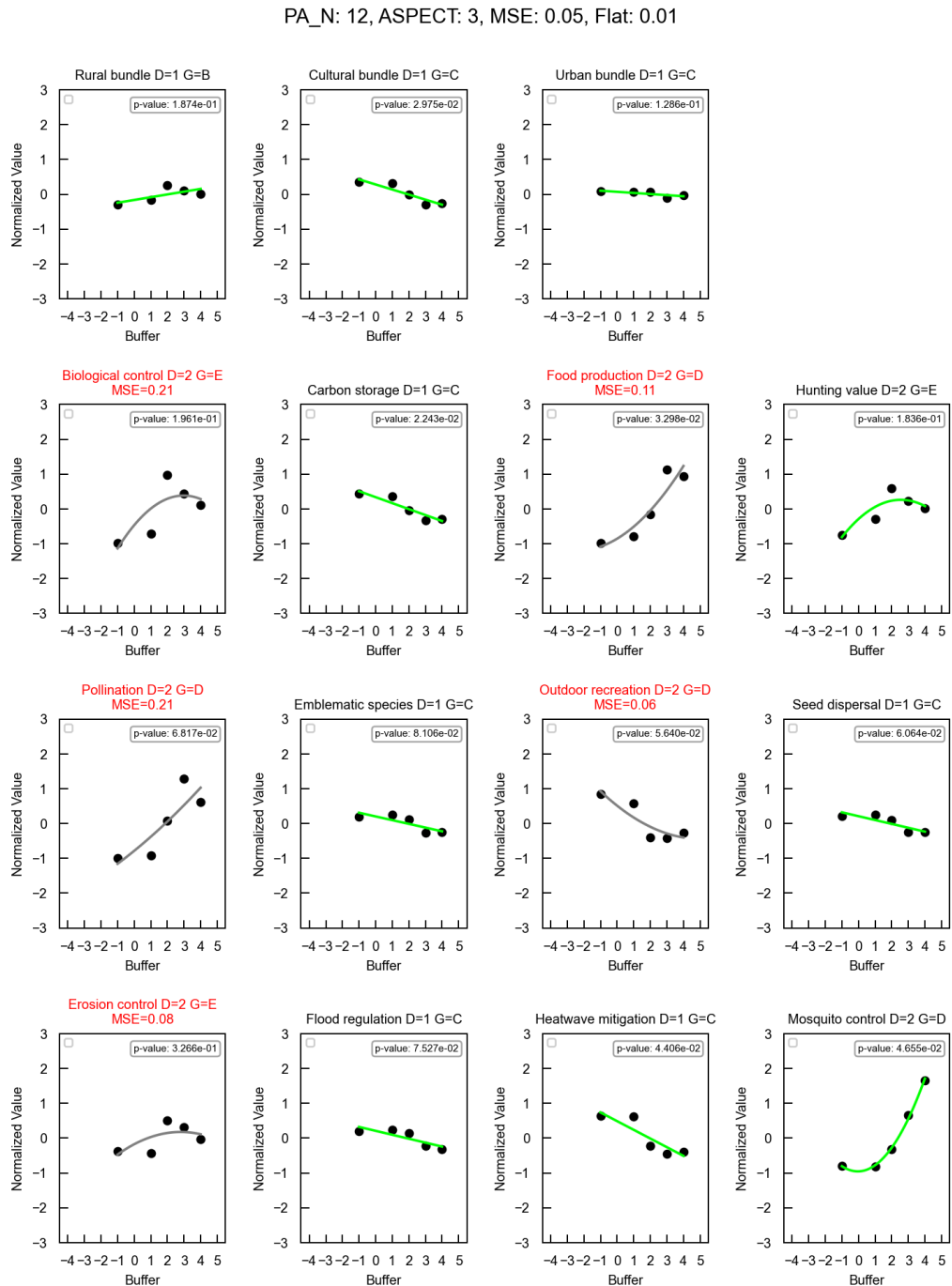

Figure S 4. Example of the gradient classification for a single border segment. The figure shows the set of polynomial regressions generated for a single border segment (Protected Area N: 12, Segment: 3) across all 12 ecosystem services and 3 aggregated bundles. Each panel displays the mean normalized ecosystem service value (Y-axis) across the 10 structural buffers (X-axis, from -4 inside to +5 outside). The algorithm fits both linear and quadratic models and selects the best fit. The title of each panel indicates the degree of the selected polynomial (D) and the resulting gradient type

classification (G, e.g., 'C' for Decreasing). The Mean Squared Error (MSE) is shown in red when it exceeds the goodness-of-fit threshold (0.05). Green lines indicate a good fit according to the defined criteria, while grey lines represent fits where the MSE exceeds the threshold. This figure illustrates the heterogeneity in gradient patterns for different ecosystem services even within the same geographical location. Figure from González-García et al. (2026).

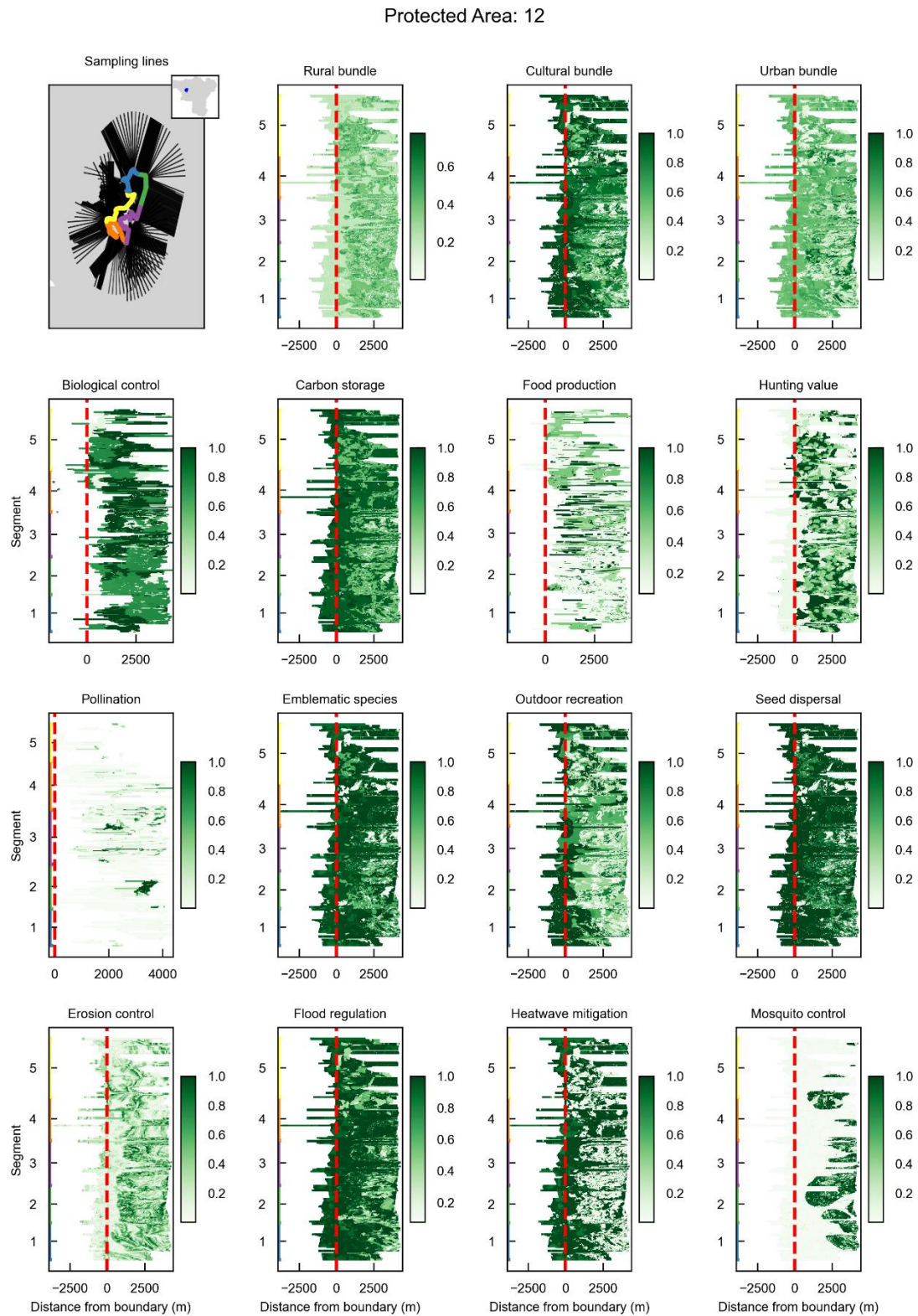

Figure S 5. Example of ecosystem service and bundle value distributions around a protected area. This figure, automatically generated by our methodology, provides a spatial interpretation of ecosystem service provision for a single protected area (Protected Area 12). The top-left panel shows the protected area's border segments (coloured

lines) and the sampling transects used for data extraction. The remaining panels are heatmaps that visualize the normalized values (colour scale, from low in white to high in dark green) for each ecosystem services and bundle. Each heatmap's Y-axis represents the different border segments, while the X-axis shows the distance from the protected area boundary (in meters). The red dashed line at $x=0$ marks the boundary, with negative distances representing the area inside the protected area and positive distances the area outside. This visualization allows for a detailed assessment of how ecosystem service provision varies both across the border and along its perimeter. Figure from González-García et al. (2026).

### E. Future Scenario Development

This section provides a detailed account of the methods used to develop the 'Participatory' and 'Technical' scenarios, complementing the summary provided in the main text. The process was centered around two participatory workshops designed to integrate local knowledge.

Participant profiles:

1. Geomatics database administrator working with metropolitan spatial data management (Grenoble metropolitan authority).
2. Foresight and strategic planning officer at the Grenoble metropolitan authority.
3. Regional coordinator for nature-based solutions for climate adaptation at the French Biodiversity Office.
4. Territorial planning officer involved in regional development processes, environmental assessment of spatial planning strategies, and topics related to risk, resilience, agriculture, and biodiversity.
5. Urban forestry and tree-planting programme officer (metropolitan "Canopy Plan").
6. Territorial planning and climate adaptation expert from a national public agency supporting governments and local authorities (Cerema).
7. Agronomist specialized in forestry, forest ecology, and plant ecophysiology.
8. Researcher studying risk perception, behavioural vulnerability, and societal resilience to hydro-climatic events.
9. Policy officer responsible for metropolitan mountain policy and relations with regional natural parks (Grenoble Metropolitan authority).
10. Researcher working on institutional change, sustainability transitions, and governance of socio-ecological systems.
11. Regional NGO representative engaged in biodiversity conservation and wildlife protection.
12. Biodiversity officer from a regional mountain territory organization.
13. Representative from a regional land conservation organization focused on habitat and biodiversity protection.
14. Representative from the regional chamber of agriculture.
15. Researcher in restoration ecology, plant community ecology, and ecological compensation.
16. Social scientist specialized in water governance and water-related socio-ecological systems.

For the first workshop, we provided stakeholders with a preliminary diagnostic that included large-format maps (A0-A2) of land use, ecosystem service bundles, and highlighted "coldspot" areas where ecosystem service provision fell within the lowest quintile (Figure S6). Working in groups for geographically distinct sub-regions (Vannier et al. 2019 ), stakeholders used these materials to identify landscape challenges and to delineate polygons directly onto the maps for their propositions of specific interventions. Researchers then translated their suggestions to nature-based solutions interventions. In a second workshop, the algorithmic rules derived from these proposals were presented for validation. This collaborative review led to crucial adjustments to the rules' technical parameters (e.g., reducing the width of new hedgerows from 10m to 5m) and a final refinement of the boundaries of intervention polygons.

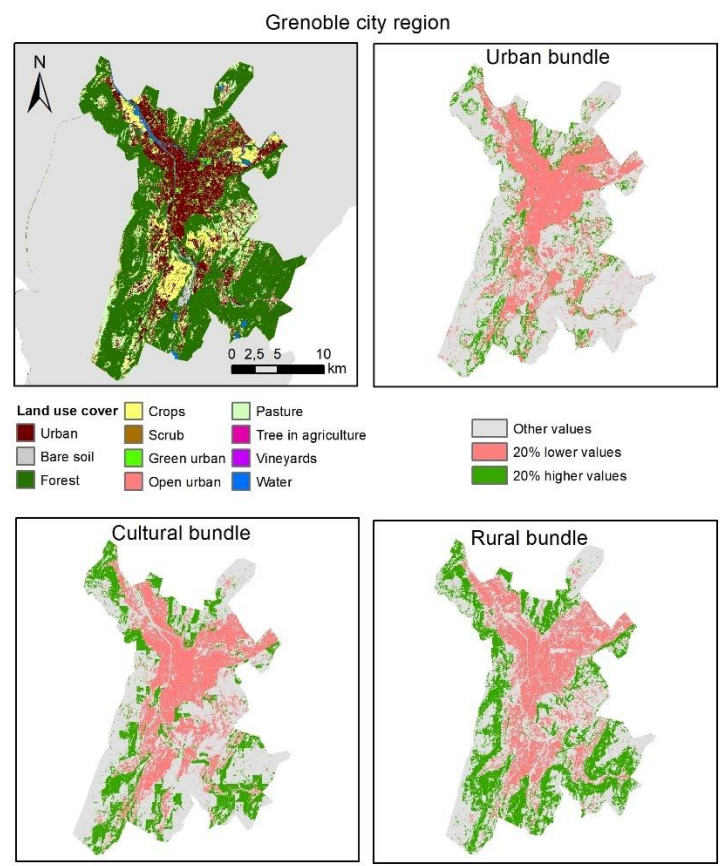

Figure S 6. Example of diagnostic maps provided to stakeholders during the first participatory workshop. The figure displays the set of maps for one of the eight study sub-regions (see Figure S1 for the whole region studied), the Grenoble city area, used to facilitate discussions on landscape challenges. It includes (top left) a land-use cover map providing general landscape context, and three maps illustrating the spatial distribution of the (top right) Urban, (bottom left) Cultural, and (bottom right) Rural ecosystem service bundles. For the bundle maps, areas with the lowest 20% of ecosystem service provision (the lowest quintile) are highlighted in red as "coldspots", while areas with the highest 20% are shown in green. These coldspot maps were the primary tool given to stakeholders. Working in small groups based on their regional expertise, participants used these large-format maps to identify and delineate specific areas where they perceived conflicts or opportunities for intervention. This visual diagnostic served as the foundation for co-designing landscape management rules.

These refined rules, detailed in the following section, were then applied following two distinct spatial allocation logics to generate the scenarios (Fig. S7). The intervention polygons shown in the figure represent the zones of eligibility where the rules were allowed to operate. The actual land-use changes occurred only on specific target pixels within these polygons, as determined by the logic of each individual rule. For the 'Participatory' scenario, the eligibility zones were the final priority polygons co-defined with stakeholders, totalling 69,327 ha (Fig. S7A). For the 'Technical' scenario, the eligibility zones were the exterior structural buffers of border segments classified as 'Decreasing Gradient' (Type C) or 'Boundary Depression' (Type D) for at least one of the three ecosystem service bundles, totalling 104,333 ha (Fig. S7B). These figures denote the eligibility zones within which interventions could be applied, not the area finally modified: within each zone, the raster rules changed only those pixels whose land-use configuration met the rule conditions. As a result, both scenarios converged on a similar directly modified area (~14.5% of the landscape) despite starting from eligibility zones of contrasting size. Although the allocation logics are distinct, the two strategies overlapped substantially, sharing 23,605 ha of eligibility zones.

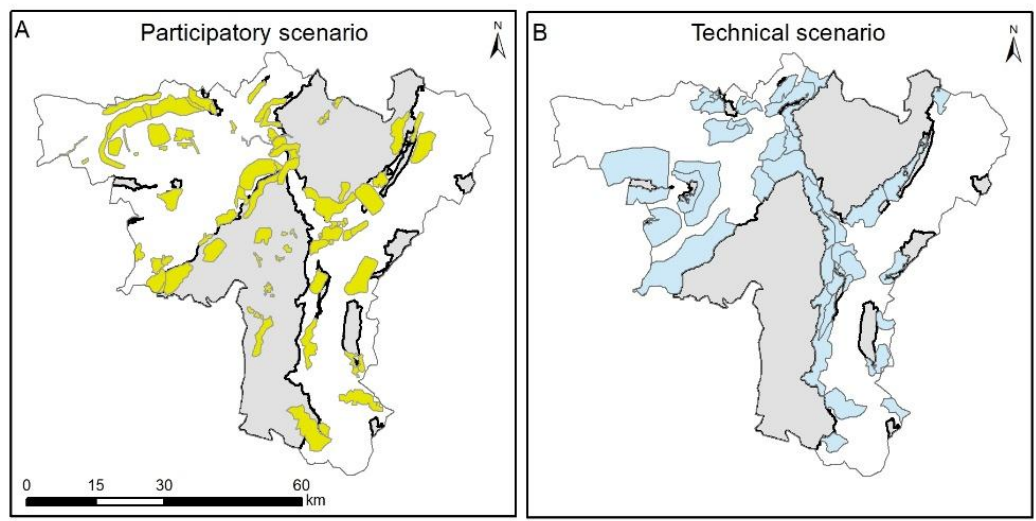

*Figure S 7. Maps of the intervention zones for the future scenarios. The figure illustrates the different spatial allocation strategies for the landscape interventions. Protected areas are shown in grey with a black outline. (A) The Participatory scenario, where the intervention zones (in yellow), totalling 69,327 ha, correspond to the priority polygons delineated by local stakeholders during participatory workshops. (B) The Technical scenario, where the eligibility zones (in light blue), totalling 104,333 ha, were selected by our diagnostic framework, targeting the exterior surroundings of all border segments classified as declining or discontinuous (Types C or D) for at least one ecosystem service bundle.*

##### a. Landscape intervention rules (Nature-based solutions implementation)

This section details the set of explicit, algorithmic land-use change rules developed to model the future scenarios. These rules translate the qualitative proposals from the stakeholder workshops into quantitative operations that modify the baseline land use/land cover map. Each rule identifies specific target pixels based on their current land use/land cover code and spatial context, and then changes their code to a new one. The rules were often applied sequentially to specific polygons representing different landscape management zones. The complete implementation of these rules, including all specific parameters, is available in the project's code repository (https://doi.org/10.5281/zenodo.17877768). The interventions and associated rules were conceptually consolidated into five main management action groups. Below is a technical description of each individual rule within these groups.

###### i. Contextual and preparatory rules

###### Urban vs. Non-Urban context definition

This rule processes the entire land use/land cover map to classify every pixel as having either an 'Urban Context' or a 'Non-Urban Context'. A pixel is assigned an 'Urban Context' if the majority (>50%) of its surrounding neighbourhood (a 5-pixel buffer) consists of urban land uses (e.g., settlement areas, industrial zones). The resulting Boolean raster layer is used as a conditional input for subsequent rules that operate differently in urban versus non-urban settings.

### Forest diversification for climate adaptation in public lands

Based on the input of a forestry expert participant in the workshop, this rule was implemented to reflect anticipated climate change adaptation strategies in public forests. It applies a baseline change to all forests managed by the French National Forest Office (ONF) across the entire study area. The rule modifies the tree species composition based on three altitudinal gradients:

• Lowland/Hill Zone (e.g., < 800 m): Forest composition is changed to 100% deciduous.
• Mountain Zone (e.g., 800 - 1600 m): Forest composition is adjusted to a 50% deciduous / 50% coniferous mix.
• Subalpine Zone (e.g., > 1600 m): Forest composition is modified to 80% coniferous / 20% deciduous.

This rule was applied as a foundational change to the baseline land use/land cover map for both the 'Participatory' and 'Technical' scenarios, creating a common, climate-adapted starting point before the application of scenario-specific interventions.

#### ii. Agricultural diversification

**Hedgerow creation:** This rule identifies the interior boundaries of large agricultural patches ($\geq 40$ pixels, i.e. 0.1 ha). It converts these boundary pixels to 'Tree cover in agricultural context' (31400), but only if none of their immediate exterior neighbours are already forested land uses, thus preventing the creation of redundant hedgerows along existing forest edges.

**Agricultural patch diversification:** This rule targets large, homogeneous patches of annual crops ($\geq 40$ pixels, land use/land cover code 21000). It reclassifies these patches by growing contiguous sub-patches of exactly 100 pixels (0.25 ha), each assigned a random crop code from a pool of cultivation types present within the local polygon.

**Targeted patch reclassification:** Within the largest meadow patch of a given polygon, this rule identifies all enclosed cultivation patches and reclassifies them to 'Broadleaf forest' (31103), effectively converting isolated farming areas within large grasslands into forest groves.

**Single meadow patch insertion:** This rule targets large agricultural patches ($\geq 40$ pixels; 0.1 ha) and randomly selects a single, contiguous group of exactly 20 pixels (0.05 ha) within them to be converted to 'Meadow' (23200).

**Scattered meadow patch insertion:** This rule iteratively places 10-pixel (0.025 ha) meadow patches (23200) within eligible agricultural and grassland areas, ensuring a minimum separation of 30 pixels (0.075 ha) between new patches to create a scattered pattern.

**Parallel lines in patches:** Within orchards (land use/land cover code 22200), this rule creates parallel lines of grassland (23200) by converting pixels at intervals of 4 (1 pixel of grass followed by 3 pixels of orchard, 5 m each 15 m), simulating the creation of grass strips between tree rows.

#### iii. Forest restoration & management

Scattered meadow patch insertion (in Forest): Within large forest patches, this rule iteratively creates small, 6-pixel meadow patches (23200) while ensuring a minimum separation of 20 pixels (0.05 ha) between them, simulating management for fire prevention and habitat diversification.

Forested wetland creation: This rule creates composite wetland-forest habitats. It first places wetland patches ($\geq 10$ pixels, 0.025 ha; land use/land cover code 41000) in eligible agricultural areas, ensuring 100m separation. It then reclassifies a subset of pixels ($\geq 8$, 0.02 ha) within each new wetland to 'Broadleaf forest' (31103).

Riparian forest restoration: This rule performs a mass conversion of all eligible non-protected pixels within specific riparian polygons to 'Broadleaf forest' (31103).

Mass conversion to forest (Rural context): This rule performs a mass conversion of all non-protected pixels within its target polygons to 'Broadleaf forest' (31103). Its operation is specifically restricted to areas previously identified as having a 'Non-Urban Context'.

##### iv. Urban greening

Linear tree addition (Non-Urban Roads): Along road pixels in 'Non-Urban Contexts', this rule converts adjacent, non-protected pixels to 'Tree cover in agricultural context' (31400) at spaced intervals of approximately 2 pixels (10 m).

Linear tree addition (Urban context roads): Along road pixels in 'Urban Contexts', this rule converts adjacent, non-protected pixels to 'Tree cover in urban context' (31450) at wider spaced intervals of approximately 4 pixels (20 m).

Urban tree patch (parking): This rule targets pixels of major roads/parking areas (12210) and iteratively grows small tree patches ($\geq 3$ pixels; 0.0075), land use/land cover code 31450), prioritizing locations near existing green areas and ensuring a 2-pixel separation (10m) between new patches.

Agricultural tree patch insertion: This rule places small tree patches ($\geq 4$ pixels, 0.01 ha; land use/land cover code 31400) in eligible agricultural and grassland areas, ensuring a minimum separation of 100m between them.

Industrial/Commercial tree patch: In industrial/commercial zones (12100), this rule iteratively places tree patches ($\geq 4$ pixels, 0.01 ha; land use/land cover code 31450), ensuring a minimum separation of 5 pixels (25m).

Road-focused greening (within connectivity zone): Along road pixels within a pre-defined "Urban connectivity zone," this rule converts adjacent, non-protected pixels to 'Tree cover in urban context' (31450) at spaced intervals of approximately 4 pixels (20 m).

Road-focused greening (Global urban context): Along road pixels located in any area identified as having an 'Urban context', this rule converts adjacent, non-protected pixels to 'Tree cover in urban context' (31450) at spaced intervals of approximately 4 pixels (20m).

##### v. River and wetland restoration

Random wetland patch: This rule iteratively places new wetland patches ($\geq 10$ pixels, 0.025 ha; land use/land cover code 41000) by growing them from random starting points within a pool of convertible agricultural and grassland pixels.

River course & channel creation: This rule identifies all existing water bodies within a polygon as connection points and uses a “skeletonization” algorithm to find a 1-pixel wide path (5m) connecting them through traversable terrain. It then converts this path to 'River network' (51100).

##### vi. Ecological corridors

Ecological corridor creation (Least-Cost Path /Zone): This rule uses a Least-Cost Path algorithm to create a 5-pixel wide (25m) ecological corridor (a mix of 31100 and 23200) connecting the two largest forest patches within a non-urban polygon.

##### b. Rule validation and refinement process

The initial set of algorithmic rules was not final but served as a technical interpretation of the stakeholders' proposals. These rules were presented back to the stakeholders during the second workshop for a crucial validation and refinement process. This collaborative review ensured that the technical parameters better reflected on-the-ground realities and local expertise. Figure S8 illustrates this iterative process with three examples, showing the landscape in its 'Present' state, after the application of our initial 'After first workshop' rule, and after the 'After second workshop' rule was refined with stakeholder feedback. For instance, Figure S8A exemplifies the refinement of Agricultural Diversification and Hedgerow Creation rules. Our initial interpretation applied a dense network of hedgerows and diversified large crop fields into very small sub-patches. Stakeholders considered this too intensive, so the final rule was adjusted to apply fewer hedgerows and create larger, more manageable 100-pixel (0.25 ha) sub-patches. Similarly, Figure S8B shows the refinement of the Agricultural Tree Patch Insertion rule. An initial proposal to insert a high density of very small tree patches into grasslands was adapted, following stakeholder feedback, to create fewer but larger tree groves that better align with practical pasture management. In other cases, such as the Urban Greening rule shown in Figure S8C, stakeholders validated our initial technical proposal without suggesting major changes, and the algorithm was maintained in its original form. This validation phase was fundamental, transforming the set of rules from a purely technical exercise into a co-designed tool that balanced ecological objectives with socio-economic feasibility.

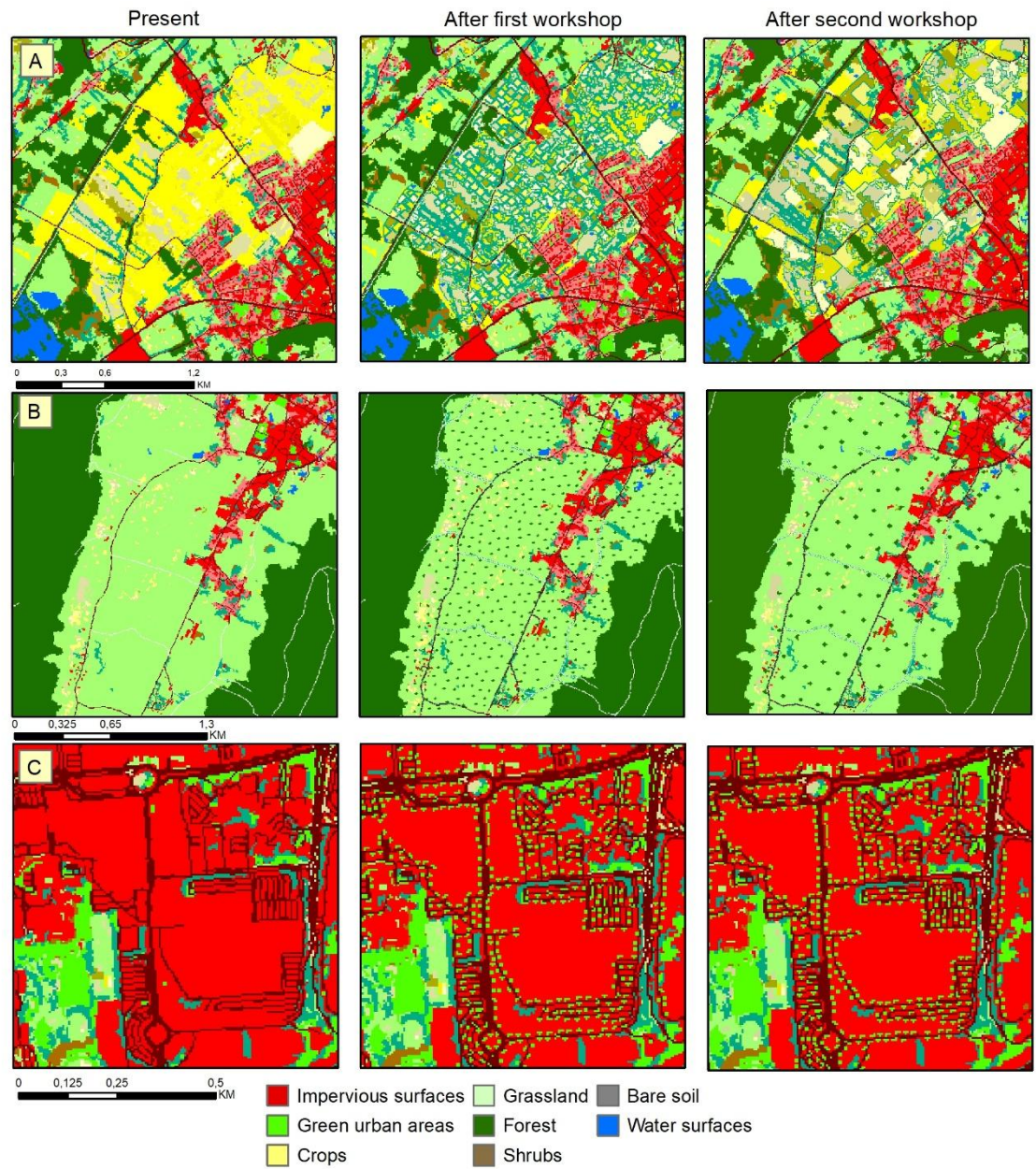

Figure S 8. Examples of the stakeholder-led rule refinement process. The figure illustrates how three initial intervention rules ('After first workshop' column) were modified based on feedback gathered during the second stakeholder workshop ('After second workshop' column). (A) Rules for Agricultural Diversification and Hedgerow Creation were adjusted from an intensive application to a more realistic mosaic of larger sub-patches and fewer hedgerows. (B) A rule for Agricultural Tree Patch Insertion was refined from creating many small, scattered tree patches to fewer, larger groves. (C) A rule for Urban Greening, designed to connect green spaces, was validated by stakeholders without significant changes. This iterative process ensured the final rules reflected both ecological goals and practical feasibility

#### c. Modifications to ecosystem service models for future scenarios

To better reflect the functional impacts of the landscape intervention rules, we introduced specific adjustments to the baseline ecosystem service models for the future scenarios. This section details the rationale and implementation of these changes, which primarily affect the Food Production model.

Several of the co-designed intervention rules, such as Hedgerow Creation and Agricultural Patch Diversification, are central components of agroecological management. While these rules imply a direct loss of productive area (e.g., a pixel of cropland converted to a hedgerow), a growing body of scientific evidence shows that agroecological practices can slightly boost the productivity of the remaining cultivated land (Van Cossel et al., 2025). This can be due to improved soil fertility, enhanced pollination and pest control, or more efficient resource use (e.g., Land equivalent ratios in agroforestry systems).

To account for this effect and avoid underestimating future food production, we applied a productivity modifier to specific land use/land cover classes within the intervention polygons where agroecological practices were assumed to be implemented. This adjustment ensures that our scenarios model not just the spatial rearrangement of land uses, but also the functional shift towards more productive, diversified agricultural systems. Based on a comprehensive literature review on the impacts of agroecological practices on crop and forage yields, we defined a set of percentage-based yield increases for relevant land use/land cover classes. These modifiers, detailed in Table S9, were applied to the base yield values of the target pixels within the intervention polygons of both the 'Participatory' and 'Technical' scenarios.

*Table S 9. Yield modifiers applied to land use/land cover classes under agroecological management assumptions.*

| LULC Class | Code Range | Assumed Agroecological Practice(s) | Yield Increase (%) | Rationale / Key References |
| --- | --- | --- | --- | --- |
| **Annual Crops (Wheat, Barley, etc.)** | 21211-21218 | Crop rotation, cover crops (legumes) | +15% | Low-to-moderate estimate, assuming general agroecological practices. (Allam et al., 2023; Marini et al., 2020; Lehmann et al., 2020; Bourgeois et al., 2022; EEA, 2022) |
| **Maize** | 21216 | Cover crops, diversified rotations | +15% | For temperate climates with legume precrops. (Bourgeois et al., 2022; Lehmann et al., 2020) |
| **Potatoes & Sugar beet** | 21221-21222 | Conservative estimate for non-insect pollinated crops | +5% | Conservative estimate assuming general soil health benefits (Lehmann et al., 2020) |
| **Sunflower & Rape** | 21231-21232 | Conservative estimate for insect pollinated crops | +10% | Conservative estimate assuming enhanced pollination from increased landscape diversity (Lehmann et al., 2020) |
| **Soya, Dry pulses & Fodder crops** | 21233-21250 | Benefits from agroecological soil management | +10% | Based on improved soil fertility and water retention. (FAO) |
| **Vineyards & Orchards** | 22100 & 22200 | Agroforestry (intercropping with crops, understory, grass) | +30% | Based on Land Equivalent Ratios (LER) for mixed systems. (EEA, 2022; Jäger & Herzog, 2021) |
| **Managed & Seminatural Grassland (Pastures & Meadows)** | 23100 & 23200 | Inclusion of agroecological practices (pasture diversification, etc.) | +20% | Assumes benefits from diversified species mixtures and improved soil fertility. (Orwin et al., 2022) |

### F. Stakeholders feasibility assessment (detailed methods)

To integrate socio-political feasibility into our analysis, we conducted a viability assessment during the second stakeholder workshop. The session began with a presentation of the preliminary results of the scenario modelling, followed by a detailed explanation of each of the algorithmic intervention rules. To facilitate in depth understanding, participants were provided with a comprehensive 50-page booklet containing a detailed description of each rule, its technical parameters, and visual examples of "before and after" landscape changes. Following this presentation, the 10 stakeholders were divided into two groups (5 participants per group). Each group was tasked with performing two evaluations for the five consolidated management action groups.

First, each group engaged in a deliberative process to collaboratively rank the five action groups from 1 (easiest to implement) to 5 (most difficult), reaching a consensus on the final order. There was a high degree of agreement between the two groups on this ranking. Second, for each action group, the groups evaluated four key viability domains: Financial, Social, Knowledge, and Governance (Bruley et al., 2025). Through discussion, they reached a consensus on a score from 1 (strong 'barrier') to 5 (strong 'enabler/lever'). During these discussions, stakeholders provided qualitative justifications for their scores. For example, Agricultural Diversification was ranked as most difficult due to "low social acceptance from farmers and restrictive Common Agricultural Policy (CAP)" (Governance barrier), while River and Wetland Restoration was ranked as easiest because of "available funding and strong technical knowledge" (Financial and Knowledge levers). The final scores used in the main analysis represent the average of the consensus scores from the two groups. The full, disaggregated results of this assessment are provided in Table S10.

*Table S 10. The table shows the consensus scores from the two stakeholder groups (G1, G2) for the perceived difficulty of each action group (1 = Easy, 5 = Hard) and for four viability domains (1 = Strong Barrier, 5 = Strong Lever). The final scores used in the main manuscript's analysis are the averages of the G1 and G2 scores. (a) They considered the availability of financing from the Common Agricultural Policy a lever, but noted under the social domain (b) that it can be either very well accepted or very poorly accepted depending on the stakeholder group. (c) This double score highlights the trade-off described by participants regarding the complexity of integrating green areas into urban spaces, which are already saturated. The four viability domains (Financial, Social, Knowledge, Governance) are qualitative and provided for context only; they were not aggregated into a numerical average and do not enter the Area-Weighted Difficulty Score, which is based solely on the consensus Difficulty ranking. Where the two groups expressed opposing views within a domain (shown as "1 or 5" or "2 or 4"), both values are reported to preserve that disagreement rather than averaging it away.*

| Action Group | Group | Difficulty | Financial | Social | Knowledge | Governance |
| --- | --- | --- | --- | --- | --- | --- |
| Agricultural Diversification | G1 | 5 | 4a | 1 or 5b | 4.5 | 1 |
| Agricultural Diversification | G2 | 5 | 4a | 1 | 5 | 2 |
| Ecological Corridors | G1 | 4 | 5 | 3 | 4.5 | 4 |
| Ecological Corridors | G2 | 4 | 1.5 | 3 | 3 | 1 |
| Forest Restoration & Management | G1 | 2 | 3 | 4 | 3.5 | 3 |
| Forest Restoration & Management | G2 | 3 | 4 | 3 | 2 | 3 |
| Urban Greening | G1 | 3 | 5 | 4 | 2.5 | 2 or 4c |
| Urban Greening | G2 | 2 | 2 | 4 | 4 | 3 |
| River and Wetland Restoration | G1 | 1 | 4 | 3 | 4 | 2 |
| River and Wetland Restoration | G2 | 1 | 4 | 4 | 3 | 4 |

During the viability assessment, stakeholders provided qualitative justifications for their scores.

For Agricultural Diversification, comments were oriented towards significant obstacles. Stakeholders consistently identified restrictive (Governance) frameworks, specifically the Common Agricultural Policy (CAP), and low social acceptance from farmers (Social) as major barriers to implementation.

For Ecological Corridors, perceptions were mixed. While there was a consensus on good social acceptance (Social) for such initiatives, stakeholders pointed to the lack of dedicated funding mechanisms for connectivity projects (Financial) as a significant barrier.

For Forest Restoration & Management, discussions highlighted the technical complexity and the need for long-term monitoring (Knowledge) as a challenge. Furthermore, potential conflicts between private land ownership and public objectives (Governance) were noted as a recurring issue.

For Urban Greening, comments were overwhelmingly positive. Stakeholders pointed to strong political willingness and its integration into urban planning (Governance), as well as high social demand for green spaces (Social), as powerful enablers.

Finally, for River and Wetland Restoration, stakeholders' comments focused on strong enabling factors, citing the availability of dedicated funding (Financial) and the existence of strong technical knowledge (Knowledge) in the region for these types of projects.

### G. Statistical analysis

#### a. Landscape-level spatial analysis

This section provides the supporting quantitative data for the spatial leverage analysis discussed in Section 3.1 of the main text. Table S11 details the total percentage of the landscape affected by changes in ecosystem service provision and calculates the specific leverage multiplier for each ecosystem service relative to the directly modified area of the Participatory (14.29%) and Technical (14.79%) scenarios.

*Table S 11. Spatial extent and directional leverage of ecosystem service provision. The table details the directional impact of the Participatory (P) and Technical (T) scenarios on the 12 modelled ecosystem services. "Area improved" and "Area declined" denote the percentage of the total study area exhibiting an increase or decrease in ecosystem service provision relative to the baseline. Multipliers quantify the spatial leverage effect, calculated as the ratio between the specific area of change (improvement or decline) and the total directly modified area (14.29% for the Participatory scenario and 14.79% for the Technical scenario). The Positive multiplier indicates the spatial amplification of benefits (area of improvement per unit of land intervention), while the Negative multiplier quantifies the spatial scale of ecosystem service reduction associated with the interventions. Values >1 indicate a propagation of effects beyond the direct directly modified area, characteristic of connectivity-dependent ecosystem services.*

| Ecosystem service | Scenario | Positive multiplier | Negative multiplier | Area improved (%) | Area declined (%) |
| --- | --- | --- | --- | --- | --- |
| Biological Control | P | 1.44 | 0.51 | 20.61 | 7.31 |
|  | T | 1.39 | 0.5 | 20.54 | 7.37 |
| Carbon Storage | P | 0.91 | 0.08 | 13.04 | 1.2 |
|  | T | 0.9 | 0.1 | 13.27 | 1.52 |
| Emblematic Species | P | 4.66 | 1.57 | 66.54 | 22.38 |
|  | T | 4.41 | 1.58 | 65.2 | 23.35 |
| Erosion Control | P | 0.14 | 0.49 | 2.07 | 7.0 |
|  | T | 0.17 | 0.47 | 2.48 | 6.98 |
| Flood Regulation | P | 0.69 | 0.28 | 9.82 | 4.05 |
|  | T | 0.7 | 0.28 | 10.28 | 4.12 |
| Food Production | P | 0.32 | 0.09 | 4.62 | 1.32 |
|  | T | 0.47 | 0.12 | 6.88 | 1.84 |
| Heatwave Mitigation | P | 0.89 | 1.38 | 12.73 | 19.73 |
|  | T | 0.89 | 1.32 | 13.19 | 19.45 |
| Hunting Value | P | 3.02 | 1.11 | 43.22 | 15.81 |
|  | T | 2.93 | 1.06 | 43.32 | 15.66 |
| Mosquito Control | P | 3.58 | 3.33 | 51.16 | 47.65 |
|  | T | 1.71 | 5.04 | 25.32 | 74.48 |
| Pollination | P | 0.43 | 0.4 | 6.11 | 5.74 |
|  | T | 0.45 | 0.39 | 6.62 | 5.74 |
| Outdoor Recreation | P | 2.9 | 0.22 | 41.44 | 3.08 |
|  | T | 2.81 | 0.21 | 41.54 | 3.08 |
|  | P | 3.56 | 2.07 | 50.93 | 29.56 |
| **Seed Dispersal** | T | 3.53 | 1.91 | 52.24 | 28.28 |

b. Modulation of consistent gradient shapes (Linear and Non-linear)

This section provides the detailed statistical evidence supporting the findings presented in Section 3.4 of the main text, specifically concerning the modulation of consistent gradient shapes. To analyze these changes, each gradient is modeled as a regression profile fitted across a series of spatial buffers that span from the interior of the protected area to the exterior matrix. Depending on the gradient typology, the profile is described by either a linear or a quadratic equation:

• Linear Gradients (Types B and C): Modeled as $y = ax + b$, where 'a' represents the slope (rate of change) and 'b' represents the intercept (vertical position). Figure S9 illustrates the sensitivity of these coefficients for stable gradients.
• Non-linear Gradients (Types D and E): Modeled as $y = ax^2 + bx + c$, where the quadratic coefficient 'a' represents the concavity (shape/depth of the valley or peak) and the constant term 'c' represents the vertical intercept (overall magnitude). Figure S10 displays the changes in these parameters for non-linear patterns.

The analysis focuses on the change in these key coefficients between the Present baseline and the future scenarios. Paired t-tests or Wilcoxon signed-rank tests were used to determine if the mean change in these coefficients was statistically different from zero.

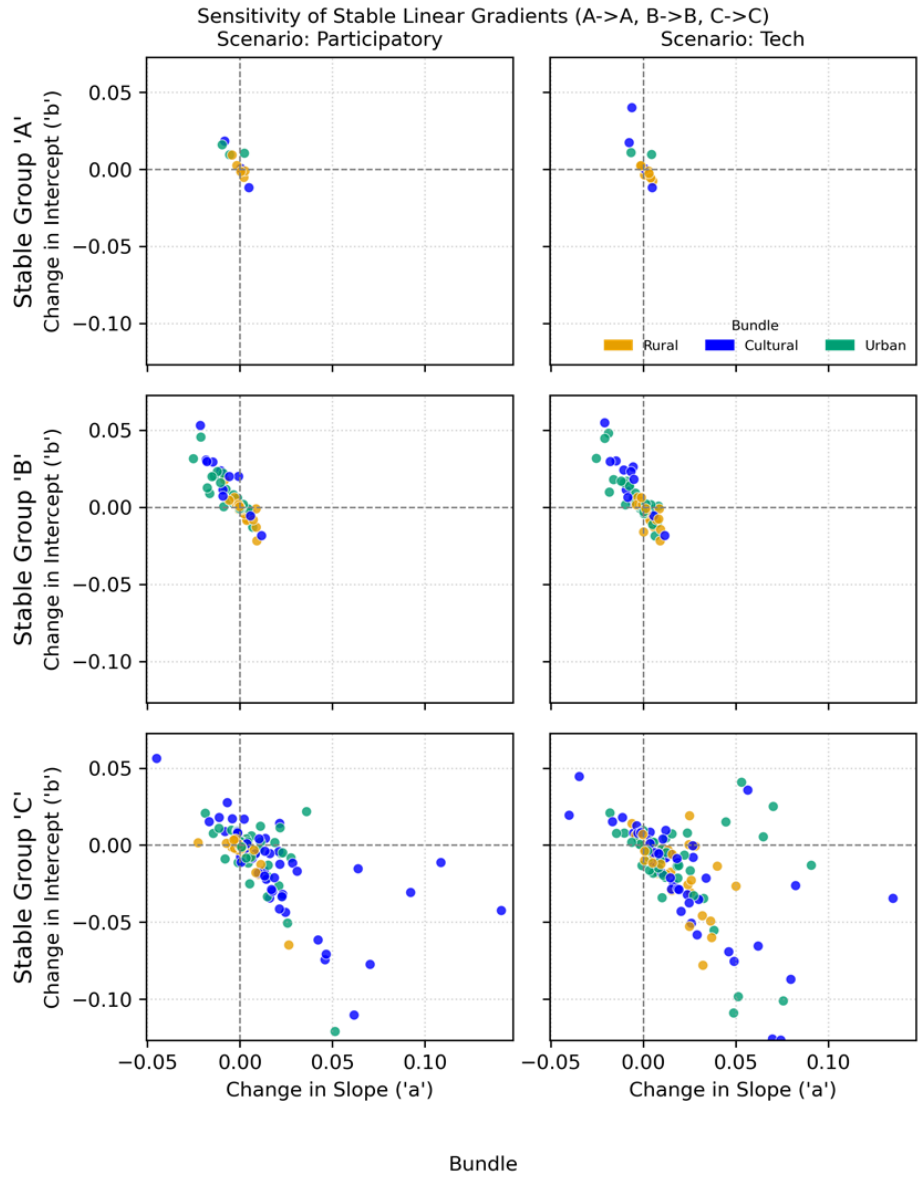

Figure S 9. Sensitivity of consistent linear gradients (Types B and C). The plots show the change in slope versus the change in intercept for border segments that remained in their linear category. The x-axis represents the change in the linear coefficient (slope 'a'), while the y-axis shows the change in the intercept ('b') of the regression $y = ax + b$. For C patterns, a positive 'a' change indicates a softening of the decline. Points in the lower-right quadrant represent a trade-off where the slope is softened at the cost of a lower intercept.

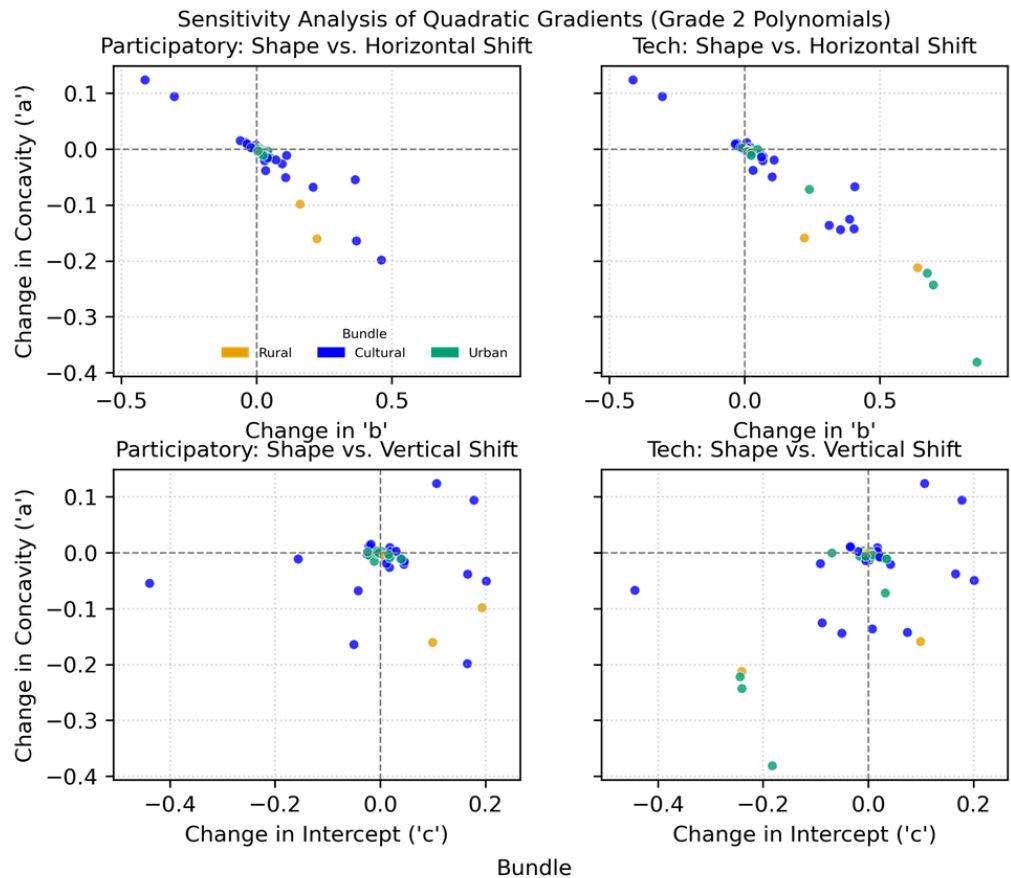

Figure S 10. Sensitivity of consistent non-linear gradients (Types D and E). The plots show the change in concavity versus the change in the vertical intercept for border segments that remained in their non-linear category. The axes represent changes in the coefficients of the quadratic regression $y = ax^2 + bx + c$. The x-axis shows the change in concavity ('a'), while the y-axis shows the change in the vertical intercept ('c'). For 'Boundary Depression' patterns (Type D, where $a > 0$), a negative 'a' change indicates a desirable flattening of the valley shape.

The results for stable Decreasing Gradients (Type C) are presented in Table S12. For these gradients, which decline from the inside out, a positive change in the slope ($a_{diff} > 0$) signifies that the rate of decline becomes less steep. This is a softening of the slope. The findings are consistent with the trade-off mechanism identified for the Technical scenario. For all three bundles under this scenario, the analysis shows a statistically significant softening of the slope ($p < 0.001$). This occurs alongside a statistically significant negative change in the intercept ($b_{diff} < 0$, $p < 0.01$), indicating that the entire fitted profile is shifted to a lower starting point. In contrast, the Participatory scenario shows a more mixed effect. While it also softens the slope for the Cultural and Urban bundles ($p < 0.01$), it does not induce a statistically consistent change in the intercept.

Table S 12. Statistical significance of coefficient changes for Stable Decreasing Gradients (C→C). The table shows the results from one-sample t-tests comparing the mean change in the slope (Mean\_a\_diff) and intercept (Mean\_b\_diff) coefficients to zero. The analysis includes all border segments that remained classified as Decreasing Gradients (Type C). The p-value for each coefficient indicates whether the observed mean change is statistically different from zero ($p < 0.05$). For these gradients, a positive Mean\_a\_diff represents a softening of the slope, while a negative Mean\_b\_diff indicates a downward shift of the profile's starting point.

| Scenario | Bundle | N | Mean_a_diff | P value | Mean_b_diff | P value |
| --- | --- | --- | --- | --- | --- | --- |
| **Tech** | Rural | 53 | 0.0103 | 0.0000 | -0.0100 | 0.0003 |
| **Tech** | Cultural | 58 | 0.0313 | 0.0001 | -0.0222 | 0.0007 |
| **Tech** | Urban | 47 | 0.0175 | 0.0000 | -0.0118 | 0.0077 |
| **Participatory** | Rural | 54 | 0.0006 | 0.4650 | -0.0020 | 0.1521 |
| **Participatory** | Cultural | 58 | 0.0159 | 0.0005 | -0.0113 | 0.0065 |
| **Participatory** | Urban | 49 | 0.0072 | 0.0002 | -0.0055 | 0.0810 |

Table S13 details the modulation of stable Increasing Gradients (Type B). The results here support the conclusion that both scenarios apply a similar moderating effect, particularly for the Urban bundle. For this bundle, both scenarios induced a significant reduction in the slope ($a_{diff} < 0$) and an increase in the intercept ($b_{diff} > 0$), with $p < 0.05$ for both coefficients in the Technical scenario and $p < 0.01$ in the Participatory. This pattern suggests a re-adjustment of the gradient shape, which elevates the profile near the border interface rather than simply making the overall gradient steeper.

*Table S 13. Statistical significance of coefficient changes for Stable Increasing Gradients (B→B). The table shows the results from one-sample t-tests comparing the mean change in the slope (Mean\_a\_diff) and intercept (Mean\_b\_diff) coefficients to zero. The analysis includes all border segments that remained classified as Increasing Gradients (Type B). The p-value for each coefficient indicates whether the observed mean change is statistically different from zero ($p < 0.05$). For these gradients, a negative Mean\_a\_diff represents a moderation of the slope, while a positive Mean\_b\_diff indicates an upward shift of the profile's starting point.*

| **Scenario** | **Bundle** | **N** | **Mean_a_diff** | **P value** | **Mean_b_diff** | **P value** |
| --- | --- | --- | --- | --- | --- | --- |
| **Tech** | Rural | 31 | 0.0017 | 0.0164 | -0.0029 | 0.0180 |
| **Tech** | Cultural | 16 | -0.0048 | 0.0512 | 0.0122 | 0.0182 |
| **Tech** | Urban | 31 | -0.0040 | 0.0171 | 0.0062 | 0.0253 |
| **Participatory** | Rural | 30 | 0.0010 | 0.1493 | -0.0014 | 0.2508 |
| **Participatory** | Cultural | 15 | -0.0053 | 0.0625 | 0.0129 | 0.0191 |
| **Participatory** | Urban | 31 | -0.0052 | 0.0010 | 0.0075 | 0.0012 |

#### c. Analysis of gradient modulation using landscape metrics

This section details the landscape metrics used to explore the biophysical drivers behind the categorical transformations (Section 3.3) and the gradient modulations (Section 3.4). These metrics were calculated for the aggregated areas Inside (buffers -1 to -5) and Outside (buffers 1 to 5) the protected area boundary for each border segment, following the methodology described in González-García et al. (2026).

i. Landscape metrics and calculation methodology

We analysed a suite of metrics representing three key components of landscape pattern, comparing the values inside and outside the protected area using the paired Wilcoxon signed-rank test (see the full model specification in the provided documents).

• **Composition Metrics:** Quantify the proportion of land-cover macro-categories:
  ○ Proportion of natural habitat: Aggregates land use/land cover codes for forests, natural grasslands, and wetlands.
  ○ Proportion of semi-natural habitat: Primarily aggregates land use/land codes for agricultural lands.
  ○ Proportion of artificial habitat: Aggregates land use/land codes for urban areas and infrastructure.
• **Structure/Diversity metric:** Quantifies the variety of land-cover types:
  ○ Shannon diversity: Calculated using the Shannon Index across the full, detailed land-use code classification.
• **Configuration/Fragmentation metric:** Quantifies the spatial arrangement:
  ○ Spatial autocorrelation (Moran's I): Used as a proxy for fragmentation. Lower values indicate a more fragmented landscape, while higher values suggest a more continuous one.

ii. Analysis for key categorical transitions (Supporting section 3.3)

This analysis provides the landscape-level explanation for the most transformative outcomes, such as the smoothing of Boundary Depressions ($D \rightarrow C$) (Fig. S11) and Decreasing Gradient ($C \rightarrow A$) (Fig. S12). Due to the low number of segments experiencing these transitions, this is an exploratory, descriptive analysis focused on characterizing the dominant change mechanism in terms of landscape metrics.

### Landscape Change ( $\Delta$ ) Driving D → C Transitions

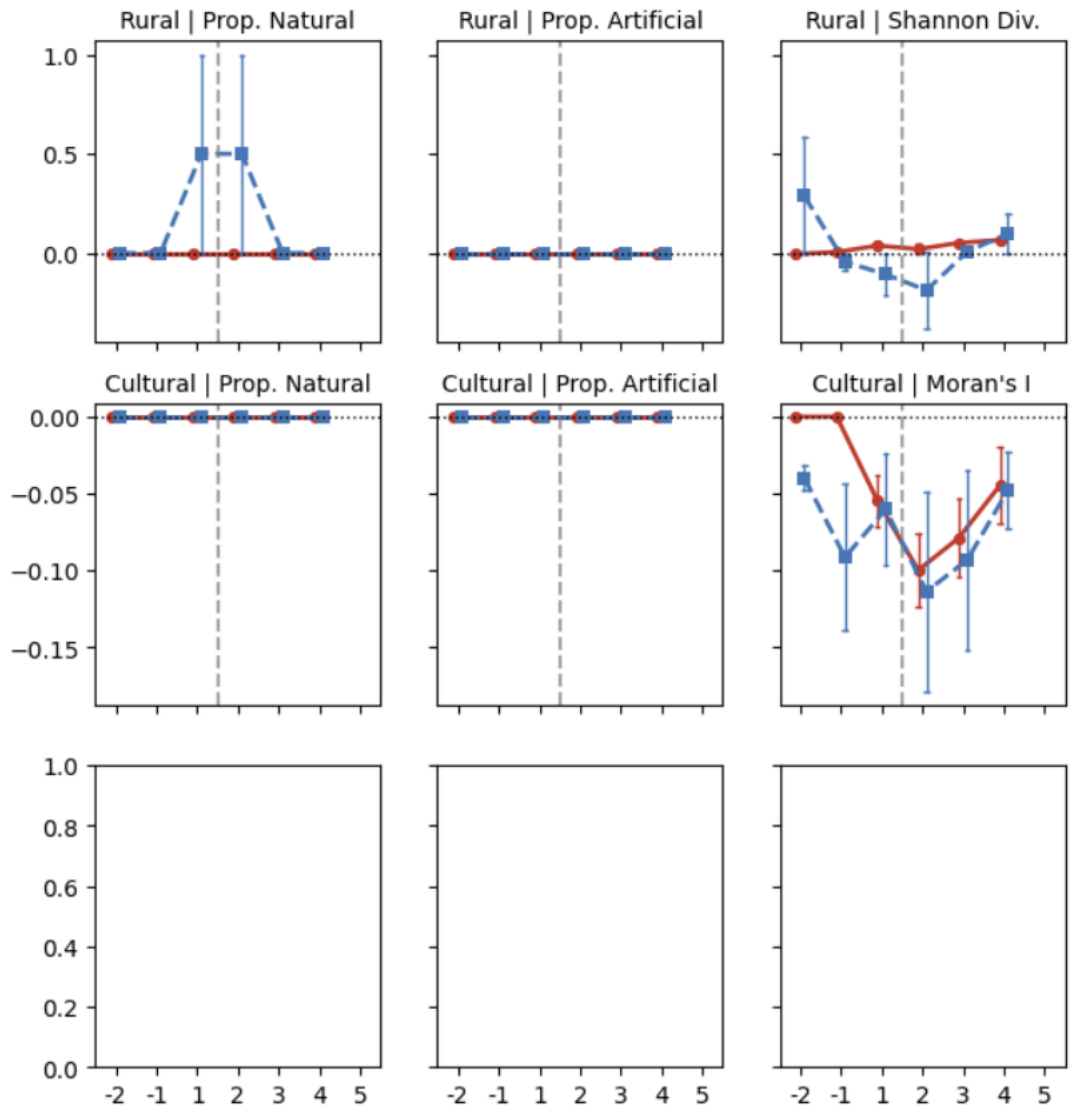

Figure S 11. This figure illustrates the difference in the change in Shannon Diversity and Proportion of Natural Habitat (Rural Bundle) and Moran's I (Cultural Bundle) that drives the smoothing of the undesirable Type D pattern into a stable C-gradient. The profile shows the distinct biophysical drivers for the correction: a strong, localized increase in Natural Habitat proportion in the outer matrix for the Rural Bundle, versus a system-wide decrease in Moran's I (increased fragmentation) for the Cultural Bundle.

### Landscape Change ( $\Delta$ ) Driving C→A Transitions

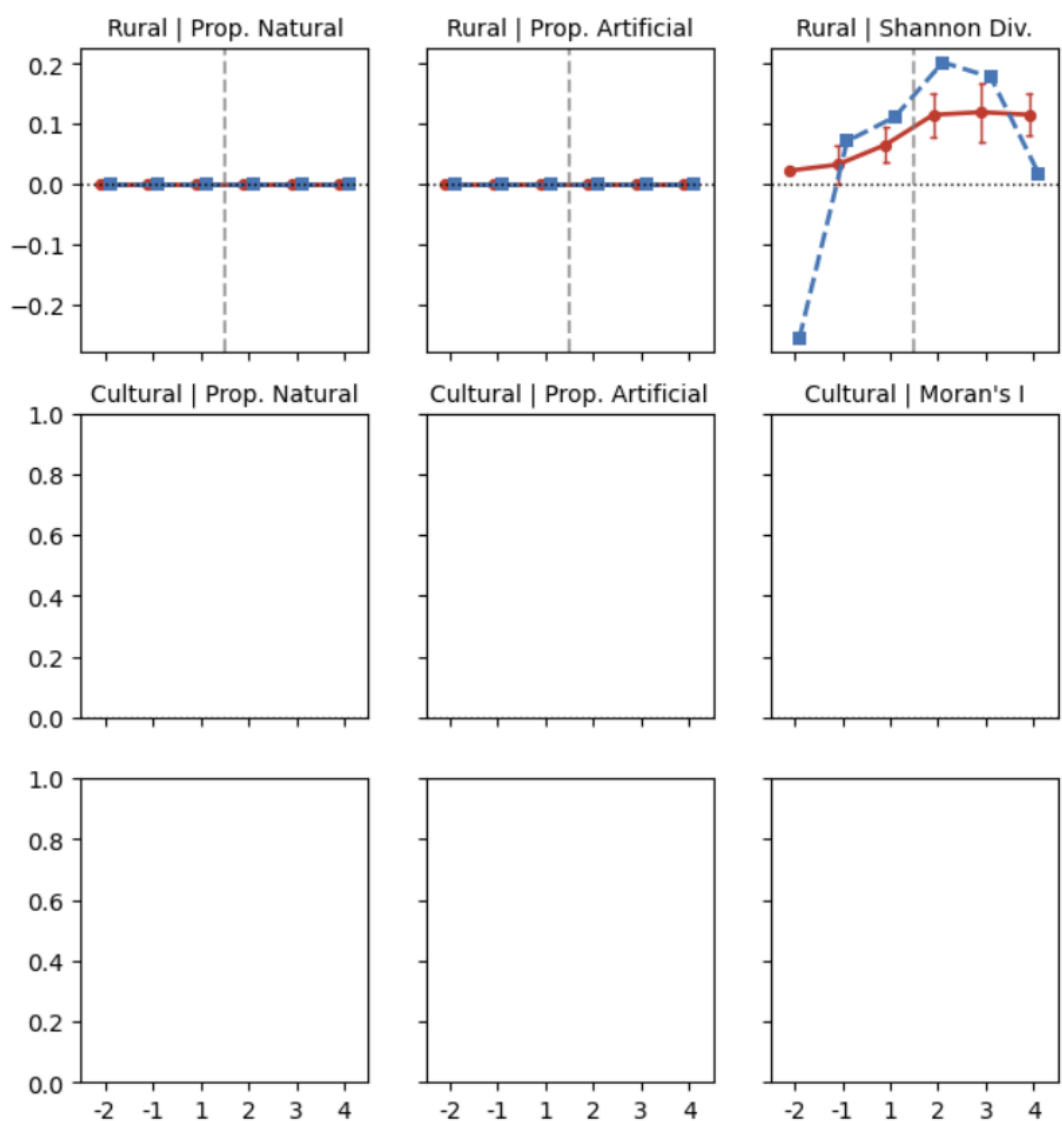

Figure S 12. This figure provides the profile for segments that achieved a Seamless Transfer (Type A), a unique outcome for the Technical scenario. The profile shows the change in the Shannon Diversity metric for the Rural Bundle, the only bundle observed to successfully transition to Type A. The finding confirms that achieving this is tied to a significant and sustained increase in the Shannon Diversity across the entire outer matrix.

#### iii. Metrics of landscape change for stable gradients (Supporting section 3.4)

This section provides the visual evidence and statistical underpinning for the landscape re-structuring that drives the systematic modulation of stable linear gradients (Types B and C), as discussed in Section 3.4 of the main text.

Figure S13 illustrates the biophysical mechanism associated with the modulation of Type C gradients. The plots show the Average Change ($\Delta$) in key landscape metrics (calculated as the mean value of the metric in the scenario minus the mean value in the Present baseline) across the structural buffers from the protected area interior (I) to the exterior matrix (O).

For the Cultural and Urban bundles, the Technical scenario (red line) consistently induces a significantly greater reduction in the Moran's I index (increased fragmentation) in the outer matrix (O1-O5) compared to the Participatory scenario (blue line). This strong, non-local reduction in homogeneity is the biophysical cause of the Technical scenario's systematic slope softening and subsequent intercept reduction (as validated in Table S14). In the Rural bundle, the scenario effects on composition are minimal, but the Technical approach achieves a greater, more consistent increase in Shannon Diversity in the outer buffers, correlating with the successful slope softening.

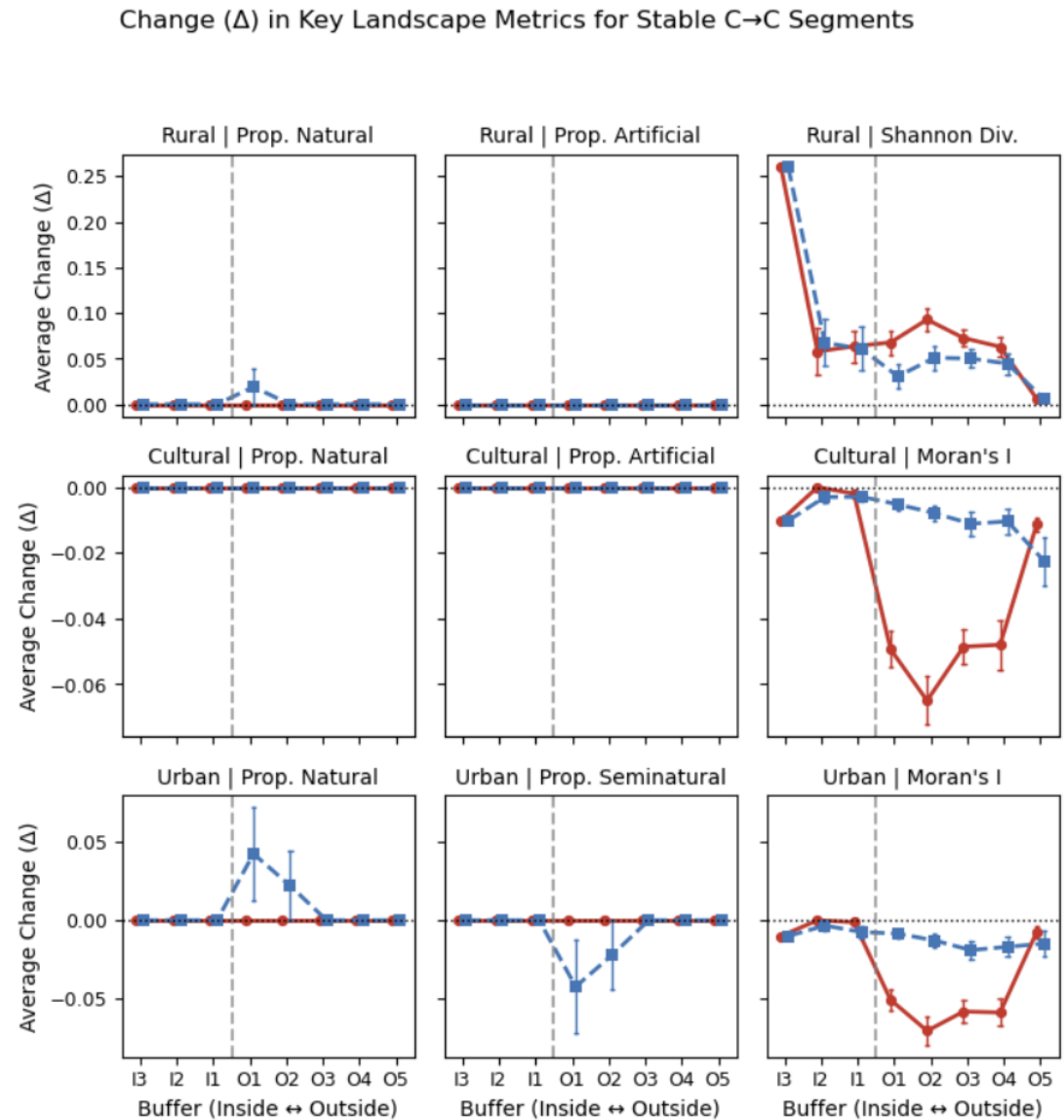

Figure S 13. Average Change ($\Delta$) in Key Landscape Metrics for Stable Decreasing Gradients (C→C) Segments. The plots show the mean change (scenario - baseline) in key landscape metrics for each bundle along the buffer gradient (I3 to O5) for segments that remained classified as Type C. The Technical scenario (red line) is compared to the Participatory scenario (blue line). A negative change in Moran's I indicates increased fragmentation, while a positive change in Shannon Diversity indicates increased landscape heterogeneity. Buffers are numbered from the interior (I) to the exterior (O).

Table S 14. Comparison of Changes in Key Landscape Metrics (Outside vs. Inside) for Stable Decreasing Gradients (C→C) between Technical and Participatory Scenarios. The change (Outside - Inside) in a given metric (e.g., Shannon Diversity) is compared between the Technical and Participatory scenarios for each buffer and bundle. The table includes all cases where a statistically significant difference was detected ($p < 0.05$). Buffers are denoted as 'O'

(Outside) followed by the buffer number (O1, O2, O3, O4), which correspond to spatial zones extending outwards from the protected area border. 'N' is the number of border segments included in the comparison for that specific buffer and bundle.

| Bundle | Metric | Buffer | N | Mean_Diff_(T-P) | Shapiro-p | p_value |
| --- | --- | --- | --- | --- | --- | --- |
| Rural | shannon_diversity | O1 | 52 | 0.037202 | 2.496217e-03 | 2.599397e-08 |
| Rural | shannon_diversity | O2 | 50 | 0.041822 | 5.225674e-07 | 1.709330e-07 |
| Rural | shannon_diversity | O3 | 45 | 0.022526 | 9.090658e-05 | 3.291596e-07 |
| Rural | shannon_diversity | O4 | 37 | 0.018365 | 1.798938e-06 | 2.067488e-04 |
| Cultural | morans_i | O1 | 58 | -0.044113 | 5.401158e-05 | 7.887818e-10 |
| Cultural | morans_i | O2 | 56 | -0.057061 | 2.619855e-05 | 2.701218e-09 |
| Cultural | morans_i | O3 | 51 | -0.037421 | 4.583408e-03 | 6.770535e-08 |
| Cultural | morans_i | O4 | 42 | -0.037560 | 1.664879e-04 | 7.999734e-06 |
| Urban | morans_i | O1 | 47 | -0.042400 | 2.407123e-05 | 3.632141e-08 |
| Urban | morans_i | O2 | 45 | -0.057937 | 5.171731e-05 | 9.053483e-08 |
| Urban | morans_i | O3 | 40 | -0.039310 | 1.079798e-02 | 1.756056e-06 |
| Urban | morans_i | O4 | 33 | -0.042069 | 1.922614e-03 | 1.488691e-05 |

Figure S14 illustrates the landscape change that underlies the "re-adjustment" of Type B gradients. For the Cultural and Urban bundles, both scenarios drive a decrease in the Moran's I index across the matrix, confirming that the re-adjustment of the gradient (slope moderation and intercept increase, as validated in Table S15) is achieved by breaking up the spatial continuity of the existing landscape patterns. This suggests concentration of ecosystem service benefits closer to the border through increased fragmentation. The Rural bundle under the Technical scenario shows a small but distinct decrease in Proportion of Natural Habitat in the outer buffers (O1, O2), accompanied by a slight, non-significant decrease in Shannon Diversity in the inner buffer (I1). This internal pattern points toward a localized trade-off that occurs near the boundary of the protected area for the Rural bundle, which the main text attributes to potential in-situ interventions.

#### Change ($\Delta$) in Key Landscape Metrics for Stable B→B Segments

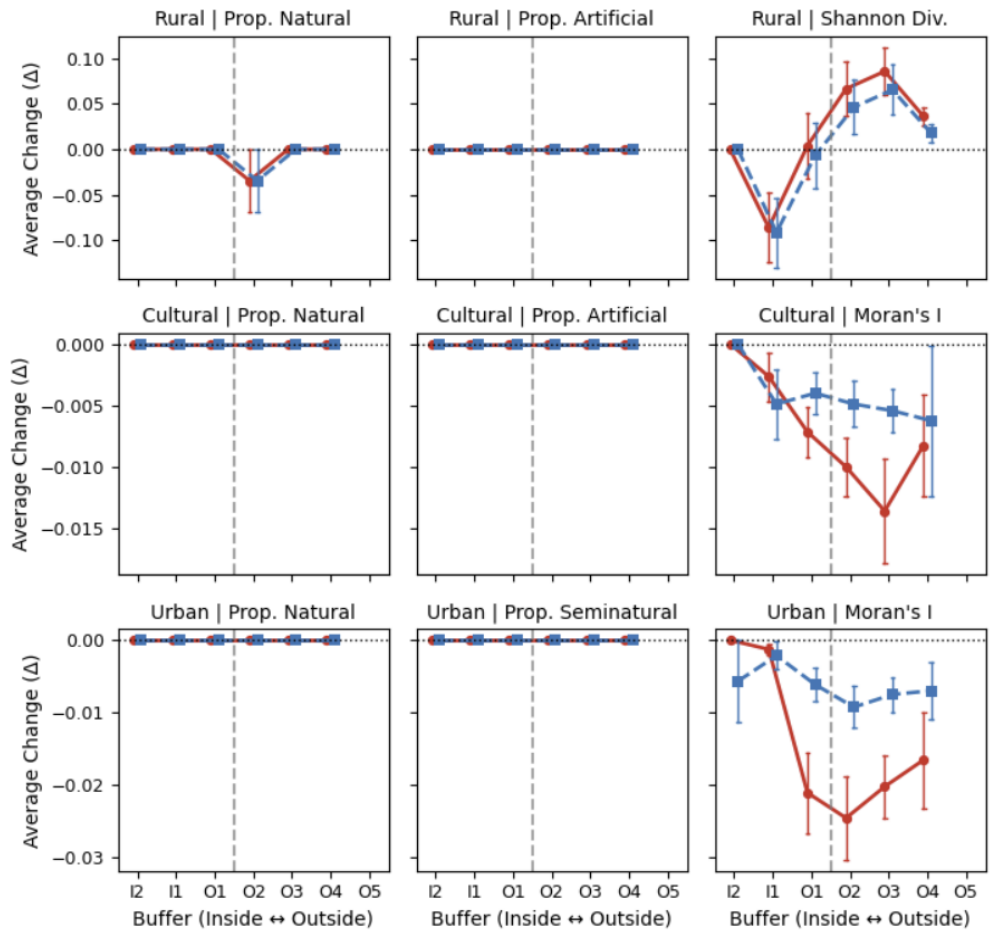

Figure S 14. Average Change ($\Delta$) in Key Landscape Metrics for Stable Increasing Gradients (B→B) Segments. The plots show the mean change (scenario - baseline) in key landscape metrics for each bundle along the buffer gradient (I2 to O4) for segments that remained classified as Type B. The Technical scenario (red line) is compared to the Participatory scenario (blue line). The profiles illustrate that the modulation in the Urban and Cultural bundles is driven by a reduction in Moran's I (increased fragmentation), suggesting a targeted strategy to enhance the interface.

Table S 15. Comparison of Changes in Key Landscape Metrics (Outside vs. Inside) for Stable Increasing Gradients (B→B) between Technical and Participatory Scenarios. The table presents the statistical comparison of the mean difference in landscape metrics between the Technical and Participatory scenarios for segments that remained classified as Increasing Gradients (Type B). The analysis covers Shannon Diversity and Moran's I across buffers.

| Bundle | Metric | Buffer | N | Mean_Diff_(T-P) | Shapiro-p | p_value |
| --- | --- | --- | --- | --- | --- | --- |
| Rural | shannon_diversity | I1 | 30 | 0.005413 | 3.744245e-11 | 0.262618 |
| Rural | shannon_diversity | O1 | 30 | 0.010624 | 5.658962e-05 | 0.001116 |
| Rural | shannon_diversity | O2 | 29 | 0.020352 | 2.102371e-07 | 0.000293 |
| Rural | shannon_diversity | O3 | 29 | 0.020321 | 9.757335e-06 | 0.000531 |
| **Rural** | shannon_diversity | O4 | 20 | 0.018665 | 2.800731e-04 | 0.021824 |
| **Cultural** | morans_i | O1 | 15 | -0.003157 | 6.693799e-03 | 0.049950 |
| **Cultural** | morans_i | O2 | 15 | -0.005124 | 1.158629e-04 | 0.011719 |
| **Cultural** | morans_i | O3 | 15 | -0.008178 | 1.632554e-05 | 0.017960 |
| **Urban** | morans_i | O1 | 31 | -0.014925 | 4.180744e-06 | 0.001019 |
| **Urban** | morans_i | O2 | 30 | -0.015301 | 2.119394e-04 | 0.002647 |
| **Urban** | morans_i | O3 | 30 | -0.012706 | 1.710674e-05 | 0.002263 |
| **Urban** | morans_i | O4 | 23 | -0.009567 | 1.521512e-06 | 0.099481 |

##### H. Complementary analysis: Characterizing gradient transitions through absolute provision values

To complement the gradient typology analysis, which focuses on the shape of the ecosystem service profiles relative to the transect mean, we performed a quantitative assessment using absolute provision values. This approach allows us to determine the physical magnitude of the changes associated with categorical transitions (e.g., determining if a transition from a 'Boundary Depression' Type D to a 'Decreasing Gradient' Type C is driven by filling the depression or by lowering the internal values). We used globally normalized provision values (scale 0–1) to ensure comparability across scenarios.

For each valid border segment, we calculated the mean provision value within each structural buffer under the Present, Technical, and Participatory scenarios. We then computed the absolute provision differential ($\Delta$) (Future Scenario minus Baseline) aggregated into three key functional zones:

**Interior change ($\Delta_{\text{interior}}$):** The mean change across the interior buffers (from -5 to -1). Non-negative values confirm that landscape interventions did reduce the ecosystem services within the protected area.

**Border change ($\Delta_{\text{border}}$):** The net change in the first exterior buffer (+1), immediately adjacent to the protected area boundary. This metric quantifies the direct improvement in ecosystem services at the border.

**Exterior change ($\Delta_{\text{exterior}}$):** The mean change across distant exterior buffers (from +3 to +5), used to assess the spatial propagation of benefits into the broader working landscape.

Finally, we grouped these metrics by ecosystem service bundle and transition type. This analysis verifies that improvements in gradient classification correspond to a net increase in ecosystem service provision and quantifies the "uplift" effect of the interventions.

Table S 16. Quantitative changes in absolute ecosystem service provision associated with gradient transitions. This table summarizes the mean difference in ecosystem service provision between Future Scenarios and the Baseline, stratified by Ecosystem Service Bundle, Scenario, and Transition Type. Calculations are based on absolute provision values (globally scaled 0–1) prior to the normalization by transect average used for gradient typology. *N* denotes the number of border segments exhibiting each transition. The metrics quantify changes across three functional zones: $\Delta_{Interior}$ represents the mean change across interior buffers (from -5 to -1), where non-negative values indicate that the intervention maintained ecosystem services within the protected area; $\Delta_{border}$ reflects the net change in the first exterior buffer (+1), quantifying direct improvements at the protected area boundary; and $\Delta_{exterior}$ indicates the mean change across distant exterior buffers (from +3 to +5), assessing the spatial propagation of benefits into the broader working landscape. Only transitions with a frequency of $N \geq 3$ are included.

| Bundle | Scenario | Transition | N | $\Delta_{Interior}$ | $\Delta_{border}$ | $\Delta_{exterior}$ |
| --- | --- | --- | --- | --- | --- | --- |
| Cultural | Technical | C->C | 58 | 0.03 | 0.04 | 0.04 |
|  |  | B->B | 16 | 0.04 | 0.03 | 0.03 |
|  |  | D->D | 13 | 0.03 | 0.03 | 0.03 |
|  |  | D->C | 5 | 0.04 | 0.06 | 0.05 |
|  |  | A->A | 4 | 0.04 | 0.03 | 0.04 |
|  |  | E->E | 4 | 0.04 | 0.05 | 0.04 |
|  | Participatory | C->C | 58 | 0.03 | 0.03 | 0.03 |
|  |  | B->B | 15 | 0.04 | 0.03 | 0.03 |
|  |  | D->D | 15 | 0.04 | 0.04 | 0.03 |
|  |  | E->E | 4 | 0.05 | 0.04 | 0.03 |
|  |  | A->A | 3 | 0.03 | 0.03 | 0.03 |
|  |  | D->C | 3 | 0.08 | 0.09 | 0.06 |
| Rural | Technical | C->C | 53 | 0.0005 | 0.0046 | 0.0064 |
|  |  | B->B | 31 | -0.0003 | 0.0019 | 0.0022 |
|  |  | A->A | 9 | 0.0006 | 0.0028 | 0.0017 |
|  |  | D->D | 6 | 0.0007 | 0.0001 | 0.0011 |
|  | Participatory | C->C | 54 | 0.0025 | 0.0017 | 0.0031 |
|  |  | B->B | 30 | 0.0004 | 0.001 | 0.0019 |
|  |  | A->A | 7 | 0.0003 | 0.0017 | 0 |
|  |  | D->D | 6 | 0.0071 | 0.0048 | 0.0005 |
| Urban | Technical | C->C | 47 | 0 | 0 | 0 |
|  |  | B->B | 31 | 0 | 0 | 0 |
|  |  | D->D | 8 | 0 | 0 | 0 |
|  |  | E->E | 8 | 0 | 0 | 0 |
|  |  | D->C | 3 | 0 | 0 | 0 |
|  | Participatory | C->C | 49 | 0.0 | -0.0104 | -0.0111 |
|  |  | B->B | 31 | 0.0029 | -0.0028 | -0.0031 |
|  |  | D->D | 12 | 0.0039 | -0.0077 | -0.0026 |
|  |  | E->E | 8 | -0.0015 | -0.0102 | -0.0115 |
|  |  | A->A | 3 | 0.0044 | 0.0015 | -0.0046 |

##### References:

- Aartsma P, Asplund J, Odland A, Reinhardt S, Renssen H (2020) Surface albedo of alpine lichen heaths and shrub vegetation. *Arctic, Antarctic, and Alpine Research* 52, 312-322.
- Al Fahdawi YMN, Al Ramahi FKM, Alfalahi ASH (2021) Measurement Albedo Coefficient for Land Cover (LC) and Land Use (LU), using remote sensing techniques, a study case: Fallujah City. *Journal of Physics: Conference Series* 1829, 012003.

- Allam, M., Radicetti, E., Ben Hassine, M., Jamal, A., Abideen, Z., & Mancinelli, R. (2023). A meta-analysis approach to estimate the effect of cover crops on the grain yield of succeeding cereal crops within European cropping systems. *Agriculture*, 13(9), 1714.
- Ambrosi L, Berger V, Rainer G, Obojes N, Tappeiner U, Tasser E, Leitinger G (2024) Spatiotemporal variability of evapotranspiration in Alpine grasslands and its biotic and abiotic drivers. *Ecohydrology*, e2633.
- Andréasson V (2023) Evaluating the impact of agriculture on albedo using Sentinel-2 data in southern Sweden. Master's Thesis.
- Attarod P, Aoki M, Bayramzadeh V (2009) Measurements of the actual evapotranspiration and crop coefficients of summer and winter seasons crops in Japan. *Plant Soil Environ* 55, 121-127.
- Bahn M, et al. (2008) Soil respiration in European grasslands in relation to climate and assimilate supply. *Ecosystems* 11, 1352-1367.
- Blumthaler M, Ambach W (1988) Solar UVB-albedo of various surfaces. *Photochemistry and Photobiology* 48, 85-88.
- Bourgeois, B., Charles, A., Van Eerd, L. L., Tremblay, N., Lynch, D., Bourgeois, G., ... & Vanasse, A. (2022). Interactive effects between cover crop management and the environment modulate benefits to cash crop yields: a meta-analysis. *Canadian Journal of Plant Science*, 102(3), 656-678.
- Bsaibes A, et al. (2009) Albedo and LAI estimates from FORMOSAT-2 data for crop monitoring. *Remote sensing of environment* 113, 716-729.
- Byczek C, Longaretti P-Y, Renaud J, Lavorel S (2018) Benefits of crowd-sourced GPS information for modelling the recreation ecosystem service. *PLoS ONE* 13(10): e0202645.
- Cancela JJ, Fandiño M, Rey BJ, Pereira LS (2010) Calibration of vineyard crop coefficients and yield response factor to support precision irrigated viticulture. *International Conference on Agricultural Engineering - AgEng 2010: towards environmental technologies*, 339.
- Cardinael R, et al. (2017) Increased soil organic carbon stocks under agroforestry: A survey of six different sites in France. *Agriculture, Ecosystems & Environment* 236, 243-255.
- Chen Y, Wang B, Pollino CA, Cuddy SM, Merrin LE, Huang C (2014) Estimate of flood inundation and retention on wetlands using remote sensing and GIS. *Ecohydrology* 7, 1412-1420.
- Chiti T, Gardin L, Perugini L, Quarantino R, Vaccari FP, Miglietta F, Valentini R (2012) Soil organic carbon stock assessment for the different cropland land uses in Italy. *Biology and Fertility of Soils* 48, 9-17.
- Cronshey R (1986) Urban hydrology for small watersheds (No. 55). US Department of Agriculture, Soil Conservation Service, Engineering Division.

- Dorendorf J, Eschenbach A, Schmidt K, Jensen K (2015) Both tree and soil carbon need to be quantified for carbon assessments of cities. *Urban Forestry & Urban Greening* 14, 447-455.
- EEA, 2022: https://forest.eea.europa.eu/topics/society/agroforestry?utm_source
- FAO (2016) *Soils and pulses: Symbiosis for life*. Food and Agriculture Organization of the United Nations, Rome.
- Foissard X, Rome S, Bigot S, Rousset E, Fouvet AC (2024) A new high spatial density temperature dataset in the Grenoble alpine valley (France) for urban heat island investigation and climate services dedicated to municipalities purposes. *Data in Brief* 55, 110553.
- Fonseca F, de Figueiredo T, Bompastor Ramos MA (2012) Carbon storage in the Mediterranean upland shrub communities of Montesinho Natural Park, northeast of Portugal. *Agroforestry Systems* 86, 463-475.
- Galleguillos M, Jacob F, Prévot L, French A, Lagacherie P (2011) Comparison of two temperature differencing methods to estimate daily evapotranspiration over a Mediterranean vineyard watershed from ASTER data. *Remote Sensing of Environment* 115, 1326-1340.
- González-García A, Neyret M, López-Tejedor A, Prima MC, Si-Moussi S, Renaud J, Gueguen M, Lavorel S (2026) Ecosystem service gradients at protected area borders reveal multiple patterns and prevalent management conflicts. *Conservation Biology*.
- González-García A, Palomo I, González JA, López CA, Montes C (2020) Quantifying spatial supply-demand mismatches in ecosystem services provides insights for land-use planning. *Land use policy* 94, 104493.
- Goswami, B.N., Venugopal, V., SenGupta, D., Madhusoodanan, M.S., and Xavier, P.K. (2006). Increasing trend of extreme rain events over India in a warming environment. *Science* 314, 1442–1445.
- Guidi C, Vesterdal L, Gianelle D, Rodeghiero M (2014) Changes in soil organic carbon and nitrogen following forest expansion on grassland in the Southern Alps. *Forest ecology and management* 328, 103-116.
- INFC (2005) *Inventario Nazionale delle Foreste e dei Serbatoi Forestali di Carbonio*. Ministero delle Politiche Agricole Alimentari e Forestali, Ispettorato Generale - Corpo Forestale dello Stato.
- IPCC (2006) *2006 IPCC Guidelines for National Greenhouse Gas Inventories*. Vol. 4: Agriculture, Forestry and Other Land Use. IGES, Japan.
- Jaafar HH, Ahmad FA, El Beyrouthy N (2019) GCN250, new global gridded curve numbers for hydrologic modeling and design. *Scientific data* 6, 145.
- Jäger, M., & Herzog, F. (2021). *Agroforestry with standard fruit trees in Switzerland. Improving production and enhancing biodiversity*.

- Kalitin NN (1930) The measurements of the albedo of a snow cover. *Monthly Weather Review* 58, 59-61.
- Kang S, Gu B, Du T, Zhang J (2003) Crop coefficient and ratio of transpiration to evapotranspiration of winter wheat and maize in a semi-humid region. *Agricultural water management* 59, 239-254.
- Karger DN, Lange S, Hari C, Reyer CPO, Zimmermann NE (2021) CHELSA-W5E5 v1.0: W5E5 v1.0 downscaled with CHELSA v2.0. *ISIMIP Repository*.
- Klein AM, et al. (2007) Importance of pollinators in changing landscapes for world crops. *Proceedings of the royal society B: biological sciences* 274, 303-313.
- Lasseur R, Vannier C, Lefebvre J, Longaretti PY, Lavorel S (2018) Landscape-scale modeling of agricultural land use for the quantification of ecosystem services. *Journal of applied remote sensing* 12, 046024.
- Lehmann, L. M., Smith, J., Westaway, S., Pisanelli, A., Russo, G., Borek, R., ... & Ghaley, B. B. (2020). Productivity and economic evaluation of agroforestry systems for sustainable production of food and non-food products. *Sustainability*, 12(13), 5429.
- Lhomme JP, Mougou R, Mansour M (2009) Potential impact of climate change on durum wheat cropping in Tunisia. *Climatic Change* 96, 549-564.
- Marini, L., St-Martin, A., Vico, G., Baldoni, G., Berti, A., Blecharczyk, A., ... & Bommarco, R. (2020). Crop rotations sustain cereal yields under a changing climate. *Environmental Research Letters*, 15(12), 124011.
- Marsoner T, Simion H, Giombini V, Egarter Vigl L, Candiago S (2023) A detailed land use/land cover map for the European Alps macro region. *Scientific Data* 10, 468.
- McRae BH, et al. (2016) Conserving nature's stage: mapping omnidirectional connectivity for resilient terrestrial landscapes in the pacific northwest. *The Nature Conservancy, Portland, Oregon*.
- Mila AJ, Akanda AR, Biswas SK, Ali MH (2016) Crop co-efficient values of sunflower for different growth stages by lysimeter study. *British Journal of Environment and Climate Change* 6, 53-63.
- Moteva M, Kazandjiev V, Zhivkov Z, Kireva R, Mladenova B, Matev A, Kalaydzhieva R (2014) Estimation of crop evapotranspiration in Bulgarian climate conditions.
- Nandi R, Mudi DK, Singh KC, Saha M, Bandyopadhyay PK (2024) Partitioning of Evapotranspiration and Crop Coefficients of Lentil Under Conserved Soil Moisture Conditions. *Journal of Soil Science and Plant Nutrition* 24, 435-450.

- O'Connor LM, Renaud J, Dou Y, Karger DN, Maiorano L, Verburg PH, Thuiller W (2024) Habitat Suitability of European Land Systems for Terrestrial Vertebrates. *Global Ecology and Biogeography* 33, e13903.
- O'Connor L, et al. (in revision) Threats to nature's contributions to people provided by terrestrial vertebrates in Europe.
- Orwin, K. H., Mason, N. W., Berthet, E. T., Grelet, G., Mudge, P., & Lavorel, S. (2022). Integrating design and ecological theory to achieve adaptive diverse pastures. *Trends in Ecology & Evolution*, 37(10), 861-871.
- Panagos P, et al. (2017) Global rainfall erosivity assessment based on high-temporal resolution rainfall records. *Scientific Reports* 7, 4175.
- Panagos P, Meusburger K, Ballabio C, Borrelli P, Alewell C (2014) Soil erodibility in Europe: A high-resolution dataset based on LUCAS. *Science of Total Environment* 479-480, 189-200.
- Paredes P, D'Agostino D, Assif M, Todorovic M, Pereira LS (2018) Assessing potato transpiration, yield and water productivity under various water regimes and planting dates using the FAO dual Kc approach. *Agricultural water management* 195, 11-24.
- Prima MC, et al. (2024) A comprehensive framework to assess multi-species landscape connectivity. *Methods in Ecology and Evolution* 15, 2385-2399.
- Ross CW, Prihodko L, Anchang JY, Kumar SS, Ji W, Hanan NP (2018) Global Hydrologic Soil Groups (HYSOGs250m) for Curve Number-Based Runoff Modeling. ORNL DAAC, Oak Ridge, Tennessee, USA.
- Rosset M, Montani M, Tanner M, Fuhrer J (2001) Effects of abandonment on the energy balance and evapotranspiration of wet subalpine grassland. *Agriculture, ecosystems & environment* 86, 277-286.
- Schulp CJE, Lautenbach S, Verburg PH (2014) Quantifying and mapping ecosystem services: Demand and supply of pollination in the European Union. *Ecological Indicators* 36, 131-141.
- Schwaab J, et al. (2015) Carbon storage versus albedo change: radiative forcing of forest expansion in temperate mountainous regions of Switzerland. *Biogeosciences* 12, 467-487.
- Sharp R, Chaplin-Kramer R, Wood S, Guerry A, Tallis H, Ricketts T (2020) InVEST 3.2.0 User's Guide. The Natural Capital Project.
- Sherpa S, Renaud J, Guéguen M, Besnard G, Mouyon L, Rey D, Després L (2020) Landscape does matter: Disentangling founder effects from natural and human-aided post-introduction dispersal during an ongoing biological invasion. *Journal of Animal Ecology* 89, 2027-2042.

- Sieber P, Ericsson N, Hammar T, Hansson PA (2022) Albedo impacts of current agricultural land use: Crop-specific albedo from MODIS data and inclusion in LCA of crop production. *Science of The Total Environment* 835, 155455.
- Si-moussi S, Thuiller W (2024) Species habitat suitability of European terrestrial vertebrates for contemporary climate and land use (Version 1) [Dataset]. German Centre for Integrative Biodiversity Research. doi: 10.25829/wpfn43.
- Sørensen MV, Strimbeck R, Nystuen KO, Kapas RE, Enquist BJ, Graae BJ (2018) Draining the pool? Carbon storage and fluxes in three alpine plant communities. *Ecosystems* 21, 316-330.
- Srivastava N, Das N, Victor U, Kumar PV (1998) Efficiency of Sunflower (*Helianthus annus* L.). *Indian Journal of Dryland Agriculture Research and Development* 13, 55-63.
- Tedela NH, et al. (2012) Runoff curve numbers for 10 small forested watersheds in the mountains of the eastern United States. *Journal of Hydrologic Engineering* 17, 1188-1198.
- Thuiller W, Georges D, Engler R, Breiner F, Georges MD, Thuiller CW (2016) Package 'biomod2'. Species distribution modeling within an ensemble forecasting framework. Version 3.3-7.
- Tian L, Zhang Y, Zhu J (2014) Decreased surface albedo driven by denser vegetation on the Tibetan Plateau. *Environmental Research Letters* 9, 104001.
- Tonolli S, Salvagni F (2007) InfoCarb Inventario Forestale del Carbonio della Provincia di Trento. Centro di Ecologia Alpina, Trento.
- Trlica A, Hutyra LR, Schaaf CL, Erb A, Wang JA (2017) Albedo, land cover, and daytime surface temperature variation across an urbanized landscape. *Earth's Future* 5, 1084-1101.
- Vannier, C., Lasseur, R., Crouzat, E., Byczek, C., Lafond, V., Cordonnier, T., ... & Lavorel, S. (2019). Mapping ecosystem services bundles in a heterogeneous mountain region. *Ecosystems and People*, 15(1), 74-88.
- Vannier, C., Bierry, A., Longaretti, P. Y., Nettièr, B., Cordonnier, T., Chauvin, C., ... & Lavorel, S. (2019). Co-constructing future land-use scenarios for the Grenoble region, France. *Landscape and Urban Planning*, 190, 103614.
- Von Cossel, M., Scordia, D., Altieri, M., & Gresta, F. (2025). Spotlight on agroecological cropping practices to improve the resilience of farming systems: a qualitative review of meta-analytic studies. *Frontiers in Agronomy*, 7, 1495846.
- Weiss A, Arkebauer TJ, Walter-Shea EA (2001) Evaluation of an algorithm for predicting albedo in heliotropic crops. *Agricultural Systems* 68, 137-150.
